# Human PRDM9 can bind and activate promoters, and other zinc-finger proteins associate with reduced recombination in *cis*

**DOI:** 10.1101/144295

**Authors:** Nicolas Altemose, Nudrat Noor, Emmanuelle Bitoun, Afidalina Tumian, Michaël Imbeault, J. Ross Chapman, A. Radu Aricescu, Simon R. Myers

## Abstract

Across mammals, PRDM9 binding localizes almost all meiotic recombination hotspots. However, most PRDM9 motif sequence matches are not bound, and most PRDM9-bound loci do not become hotspots. To explore factors that affect binding and subsequent recombination outcomes, we mapped human and chimp PRDM9 binding sites in a human cell line, and measured PRDM9-induced H3K4me3 and gene expression changes. These data revealed varied DNA-binding modalities of PRDM9, and histone modifications that predict binding. At sites where PRDM9 binds, specific *cis* sequence motifs associated with TRIM28 recruitment, and histone modifications, predict whether recombination subsequently occurs. These results implicate the large family of KRAB-ZNF genes in consistent, localized meiotic recombination suppression. PRDM9 affects gene expression for a small number of genes including *CTCFL* and *VCX*, by binding nearby. Finally, we show that PRDM9’s DNA-binding zinc finger domain strongly impacts the formation of multimers, with a pair of highly diverged alleles multimerizing less efficiently.

## Introduction

In humans and mice, PRDM9 determines the locations of meiotic recombination hotspots (***Baudat et al., 2010; Myers et al., 2010; Parvanov et al., 2010***). PRDM9 is expressed early in meiotic prophase (***Sun et al., 2015***), during which its C2H2 Zinc-Finger (ZF) domain binds DNA at particular motifs and its PR/SET domain trimethylates surrounding histone H3 proteins at lysine 4 (H3K4me3; ***Hayashi et al., 2005***), a mark also found at the promoters of transcribed genes (***Santos-Rosa et al., 2002***), and also at lysine 36 (H3K36me3; ***Wu et al., 2013; Eram et al., 2014; Powers et al., 2016; Davies et al., 2016; Grey et al., 2017***). At a subset of PRDM9 binding sites, SPO11 is recruited to form Double Strand Breaks (DSBs) (***Neale and Keeney, 2006; Smagulova et al., 2011***). These DSBs undergo end resection and the resulting single-stranded DNA ends are decorated with the meiosis-specific protein DMC1 (***Neale and Keeney, 2006***).

*In vivo* experiments to date have mapped the locations of intermediate events in recombination by performing Chromatin ImmunoPrecipitation with high-throughput sequencing (ChIP-seq) against the H3K4me3 mark and the DMC1 mark in testis tissue from mice and humans (***Baker et al., 2014; Smagulova et al., 2011; Brick et al., 2012; Pratto et al., 2014; Davies et al., 2016***), or by sequencing DNA fragments that remain attached to Spo11 after DSB formation (***Lange et al., 2016***). Recent studies have also published direct PRDM9 ChIP-seq results using a custom antibody in mouse testes (***Baker et al., 2015a; Walker et al., 2015; Grey et al., 2017***). To study the DNA-binding properties of mouse PRDM9, one study sequenced genomic DNA fragments bound *in vitro* by recombinant proteins containing only the PRDM9 ZF array (***Walker et al., 2015***). In humans, indirect binding maps, as well as recombination hotspots identified by Linkage Disequilibrium (LD) maps (***Myers et al., 2005***), have enabled the discovery of human PRDM9 binding motifs (***Myers et al., 2008, 2010; Hinch et al., 2011; Pratto et al., 2014; Davies et al., 2016***). However, these published motifs are neither sufficient nor necessary to predict genome-wide PRDM9 binding, DSB formation, or recombination events (***Myers et al., 2010; Pratto et al., 2014***), and it has been suggested that binding might be influenced by chromatin features in ***cis (Walker et al., 2015***). Moreover, not all PRDM9 binding sites become hotspots (***Baker et al., 2014; Grey et al., 2017***), and the reasons for this remain unclear. In particular, apart from PRDM9 motifs themselves there are no specific DNA sequence features that have been shown to modulate recombination rate in *cis* in mammals, nor epigenetic modifications shown to play a causal role genome-wide.

PRDM9 has been hypothesized to play a role in meiotic gene regulation given its H3K4 trimethylase activity (***Hayashi et al., 2005; Mihola et al., 2009***). In fact, PRDM9 was shown to be transcriptionally activating in an early reporter gene assay (***Hayashi et al., 2005***). However, this model for PRDM9’s function in meiosis has fallen out of favor given recent experiments that demonstrate full fertility in transgenic mice with completely remodeled PRDM9 binding landscapes (***Baker et al., 2014; Davies et al., 2016***). This does not preclude the possibility that PRDM9 may play a secondary gene regulatory role in meiosis. PRDM9 has also been shown to bind to itself and form multimers, and its DNA-binding and trimethylation behaviors remain active in PRDM9 multimers (***Baker et al., 2015b***). However, it is not known which domains of PRDM9 mediate this multimer formation activity and whether different combinations of PRDM9 alleles may form hetero-multimers with different efficiencies.

To investigate the properties of PRDM9’s zinc-fingers in humans and chimpanzees as they relate to the questions posed above, we expressed various engineered versions of PRDM9 in a mitotic human cell line (HEK293T), then performed multimodal high-throughput sequencing analyses. While this approach will fail to reproduce cell-type-specific phenomena found only in spermatocytes and oocytes, it enables us to infer the fundamental rules governing the behavior of PRDM9 in the nucleus, and as we describe below replicates many of the key properties of PRDM9 binding *in vivo*. In these cells, we performed ChIP-seq against human PRDM9, H3K4me3, H3K36me3, and chimp PRDM9, as well as ATAC-seq (Assay for Transposase-Accessible Chromatin with high-throughput sequencing) to examine nucleosome positioning and DNA accessibility, and RNA-seq to examine gene expression (all samples listed in Methods and Materials). Importantly, by comparing data from transfected and untransfected cells (in which there is weak to no endogenous *PRDM9* expression), we can observe the same genomic sites with and without the effects of PRDM9 expression. This approach also allows us to rapidly engineer and test various different alleles and truncations of PRDM9 to explore the properties of its individual domains. Further, our results are complemented by previously published data on LD-based recombination crossover hotspots (***Myers et al., 2005***), DSB hotspots decorated by DMC1 (***Pratto et al., 2014***), H3K4me3 in human testes (***Pratto et al., 2014***), and histone modifications across human cell types (***ENCODE, 2012; Kundaje et al., 2015***). As described below, the results also implicate a widespread role for ZF-array binding by a host of *other* zinc-finger containing genes (***Imbeault et al., 2017***), in suppressing, rather than activating, recombination in humans.

## Results

### A map of direct PRDM9 binding in the human genome

We performed ChIP-seq in HEK293T cells transfected with the human PRDM9 reference allele (the “B” allele) containing an N-terminal YFP tag, which was targeted for immunoprecipitation. To identify regions bound by PRDM9, we modeled binding enrichment relative to a measure of local background coverage at each position in the genome, then performed a likelihood ratio test for evidence of binding above background (detailed in Methods and Materials). This yielded 170,198 PRDM9 binding peaks across the genome (p<10^-6^), demonstrating that PRDM9 can bind with some affinity to many sites outside of recombination hotspots, which number in the tens of thousands (***Myers et al., 2005; Pratto et al., 2014***), similar to findings in mice (***Baker et al., 2014; Walker et al., 2015***). This large number of peaks likely results from the high expression level of PRDM9 in this system, providing sensitivity to detect even weak binding interactions. Weak PRDM9 binding interactions such as these may help to explain the ~40% of DSB events that occur outside known hotspots in mice (***Lange et al., 2016***).

We compared our ChIP-seq data with a set of 18,343 published *in vivo* human DSB hotspot peaks from DMC1 ChIP-seq experiments in testis samples (***Pratto et al., 2014***). We found evidence for binding at up to 74% of DSB hotspots (at p<10^-3^) after correcting for chance overlaps, demonstrating that even in a completely different cell type and expression system, PRDM9 binds the majority of hotspots. The proportion bound in our system is greater (up to 82%) at DSB hotspots not subject to the telomere effect, which substantially increases the probability of DSB formation within roughly 15 Mb of each telomere in human male meiosis (***Pratto et al., 2014***; ***Figure 1***-S1a). The probability of overlapping DSB hotspots and testis H3K4me3 ChIP-seq peaks (***Pratto et al., 2014***) also increases with the strength of PRDM9 binding in our system (***Figure 1***b), and conversely the probability of overlap increases for hotter DMC1 peaks, especially in non-telomeric regions (***Figure 1***-S1b).

**Figure 1.**
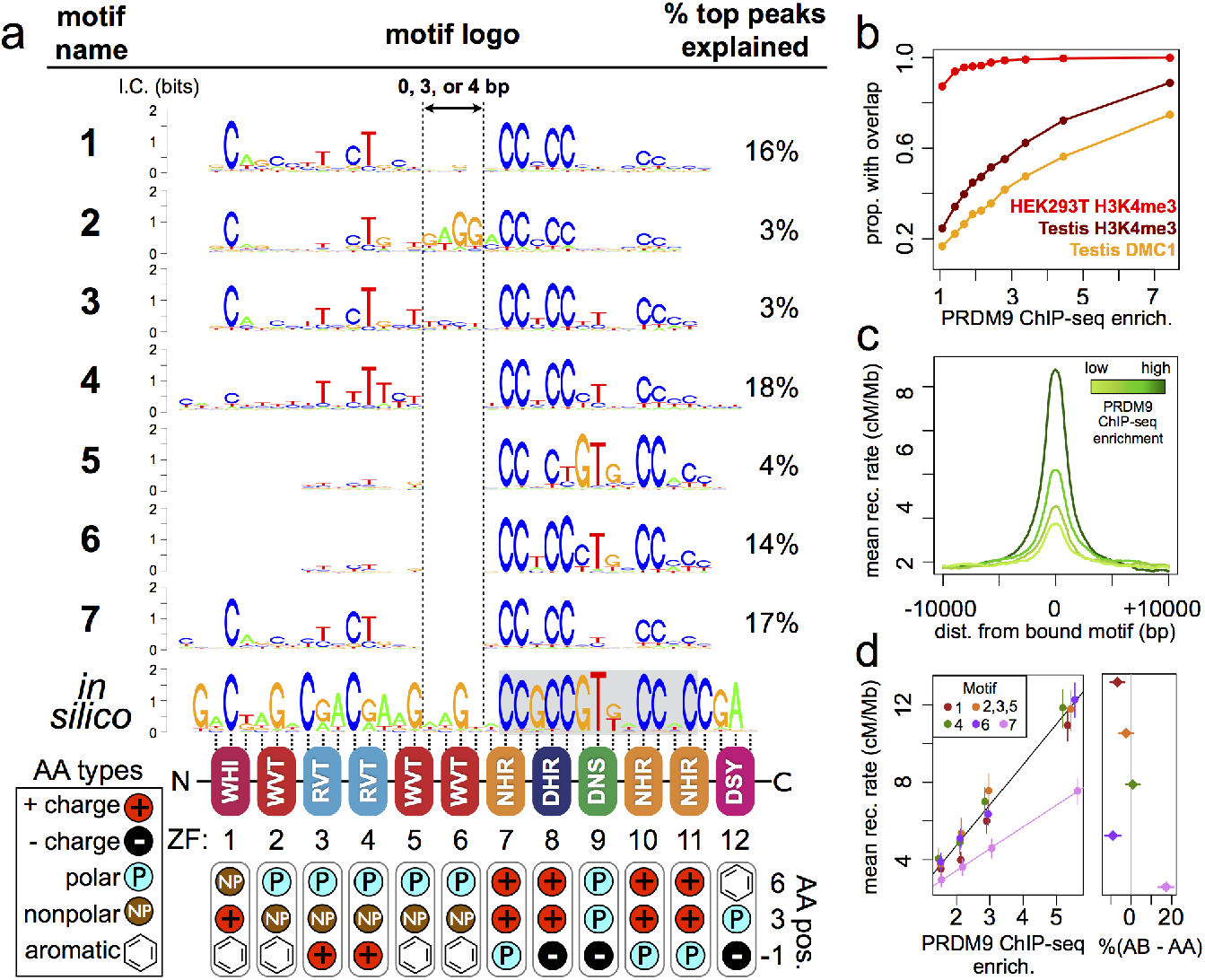
Comparison of seven distinct motifs bound by human PRDM9 (B allele). **a**: Seven motif logos produced by our motif-finding algorithm (applied to the top 5,000 PRDM9 binding peaks ranked by enrichment, after filtering out repeat-masked sequences) were aligned to each other and to an *in silico* binding prediction (***Myers et al., 2010***; ***Persikov et al., 2009; Persikov and Singh, 2014***), to maximize alignment of the most information-rich bases. The position of the published hotspot 13-mer is indicated by the gray box overlapping the *in silico* motif (***Myers et al., 2008***). The right side lists the percent of the top 1000 peaks ranked by enrichment (without further filtering) containing each motif type. Zinc finger residues at DNA-contacting positions (labeled −1, 3, 6) are illustrated below each zinc finger position, classified by polarity, charge, and presence of aromatic side chains. Zinc fingers 5 and 6 lack positively charged amino acids and contain aromatic tryptophan residues, and they coincide with a variably spaced motif region (indicated by vertical dotted lines). Motif 4 is truncated. **b**: H3K4me3 ChIP-seq data from PRDM9-transfected HEK293T cells (this study) and H3K4me3/DMC1 data from testes (***Pratto etal., 2014***) were force-called in a 1-kb window centered on each PRDM9 binding peak center (p<10^-6^, minimum peak separation 1000 bp) to provide a p-value for enrichment of each H3K4me3/DMC1 sample at each PRDM9 peak. In our parameterization, the enrichment value represents the fold enrichment over background, minus 1, at the base with the smallest p-value within each peak region. Peak windows with fewer than 5 input reads from cells or testes were filtered out, to improve enrichment estimates, and windows with excessive genomic coverage (in the top 0.1%ile) or IP coverage (>500 combined fragments) were removed to avoid outliers due to mapping errors. PRDM9 peaks overlapping H3K4me3 peaks from untransfected cells were removed, leaving 37,188 peaks passing all filters. Peaks were split into deciles according to their PRDM9 enrichment values, and the proportion of peaks with a force-called H3K4me3 or DMC1 p-value <0.05 is plotted within each decile. **c**: Peaks were stratified into quartiles based on increasing PRDM9 enrichment (light green to dark green) after filtering out promoters. Mean recombination rates (from the HapMap LD-based recombination map ***HapMap, 2007***) at each base in the 20-kb region centered on each bound motif are plotted for each quartile, with smoothing (ksmooth, bandwidth 25). **d**: Peak enrichment quartiles (filtered to remove promoters as in c) were separated by motif type (motifs 2, 3, and 5 were combined due to low abundance), and the mean HapMap recombination rate overlapping peak centers was plotted against median PRDM9 enrichment in each quartile, with lines of best fit added for Motif 7 versus all other motifs (left plot). Plot showing the difference in the percentage of AB-only DMC1 peaks versus AA-only DMC1 peaks (***Pratto etal., 2014***) containing each motif type (right plot). Error bars indicate two standard errors of the mean (left plot) or 95% bootstrap confidence intervals (right plot). **Figure 1-Figure supplement 1**. See Figure Supplements **Figure 1-Figure supplement 2**. See Figure Supplements **Figure 1-Figure supplement 3**. See Figure Supplements **Figure 1-Figure supplement 4**. See Figure Supplements

To investigate the histone methylation activity of PRDM9 and to provide an additional marker of PRDM9 binding, we also performed ChIP-seq against the H3K4me3 mark in both transfected and untransfected cells by the same method. After subtracting sites overlapping “pre-existing” H3K4me3 peaks (those present in untransfected cells), we found that 95% of PRDM9 binding peaks show H3K4me3 following transfection (p<0.01), and this proportion increases to 100% with increasing PRDM9 binding enrichment (see *Figure 1*b). That is, PRDM9 makes the H3K4me3 mark essentially everywhere it binds, regardless of the pre-existing chromatin substrate, and the strength of the H3K4me3 signal correlates with the strength of PRDM9 binding (*r* = 0.48, ***Figure 1***-S2). As observed in mice (***Davies et al., 2016; Powers et al., 2016; Grey et al., 2017***), we also observe localized H3K36me3 deposition at bound sites (see ***Figure 1***-S1d). Further, full-length PRDM9 preferentially binds more open chromatin, and appears to phase surrounding nucleosomes (see ***Figure 1***-S1h), again as seen in mice (***Baker et al., 2014***). However, the zinc finger domain by itself appears unable to phase nucleosomes (see ***Figure 1***-S1g).

Next, we compared enrichment values for PRDM9 and H3K4me3 in our cells with *in vivo* testis H3K4me3 and DMC1 enrichment values computed from published raw data (***Pratto et al., 2014***) (see Methods and Materials). PRDM9 enrichment in our HEK293T cells correlates with testis H3K4me3 enrichment (*r* = 0.50), but shows a much lower raw correlation with testis DMC1 enrichment (*r* = 0.21), consistent with a layer of DSB regulation occurring downstream of PRDM9 binding and H3K4me3 marking (***Figure 1***-S2), which we show below does indeed occur. Taken alone, the testis H3K4me3 data are a poor predictor of testis DMC1 heat, due to low signal in the dataset and a large number of peaks not overlapping DMC1 hotspots (***Pratto et al., 2014***). However, by measuring H3K4me3 enrichment *only* at PRDM9 peaks identified in our cells, we see a stronger correlation between testis H3K4me3 enrichment values and DMC1 heat (*r*=0.31, and up to 0.55 if we remove telomeres; ***Figure 1***-S2). This implies that some, though not all, of the differences between our peaks and hotspot occurrences reflect differences in PRDM9 binding *strength*, despite sharing of binding site positions, between HEK293T and meiotic cells.

Finally, LD-based recombination rates (***HapMap, 2007***) peak around our PRDM9 binding peak centers, and the local recombination rate increases with PRDM9 binding strength (***Figure 1***c-d). Thus, despite cell-type differences between our HEK293T expression system and the chromatin environment of early spermatocytes, our binding peaks capture the majority of biologically relevant recombination hotspots and reveal many additional non-hotspot sites bound by PRDM9.

### Binding motifs reveal multiple modes of PRDM9 binding

Next, we searched for sequence motifs occurring near PRDM9 binding sites using a Bayesian *de novo* motif finding algorithm (described in ***Davies et al., 2016*** and in Methods and Materials). Rather than a single motif described by a position weight matrix (PWM), this algorithm allows binding sites to be described by a mixture of multiple motifs. The algorithm identified seven distinct non-degenerate motifs each highly enriched in the central 100 bp of each PRDM9 ChIP-seq peak (***Figure 1***a; detailed in Methods and Materials). Together, they explain 75% of the top 1,000 binding peaks, falling to 53% of all peaks. The remaining peaks contain mostly degenerate, GC-rich sequences (***Figure 1***-S3), similar to DMC1 hotspots in transgenic mice containing this same allele (***Davies et al., 2016***) and interpretable as binding to clusters of individually weaker motif matches in mostly GC-rich regions.

While each of the seven motifs has a close internal match to the published 13-mer found in human recombination hotspots (***Myers et al., 2008***), each motif is much longer, with five motifs fully spanning the maximal possible ~36-bp expected binding footprint of PRDM9’s 12 canonical zinc fingers (***Figure 1***a). Therefore, the zinc fingers predicted to bind upstream of the published 13-mer are influential for binding and show high sequence specificity, and they explain a less specific extended motif reported in (***Myers et al., 2008***). Aligning these motifs to each other and to an *in-silico* motif prediction (***Myers et al., 2010; Persikov et al., 2009; Persikov and Singh, 2014***), shows that they differ mainly according to internal spacings within the motif (***Figure 1a***) although also somewhat in base-pair preferences (*e.g*. Motif 5). The region corresponding to ZF5 and ZF6 is predicted to span 6 bp, but in Motifs 4-7 this region spans only 2 bp, and in Motif 1 it spans only 5 bp. Interestingly, the expected 6-bp binding footprint is observed only for Motifs 2 and 3, which explain a relatively small proportion of peaks (6%). This alternative spacing cannot be captured in a single motif, possibly explaining why upstream zinc fingers have shown weak or no sequence specificity in previously published hotspot motifs (***Myers et al., 2010; Hinch et al., 2011; Pratto et al., 2014***).

Alternative spacing within motifs could explain how long zinc finger arrays like PRDM9’s are able to consecutively bind DNA despite theoretical physical constraints (***Persikov and Singh, 2011***), similar to multivalent CTCF binding (***Nakahashi et al., 2013***). Our results are also consistent with recent findings that truncated mouse PRDM9 alleles can stably bind discontinuous submotifs, though at reduced specificities, with subsets of zinc fingers (***Striedner et al., 2017***). ZF5 and ZF6, which overlap the variably spaced region, have large aromatic tryptophan residues at the DNA-contacting “-1” position (***Figure 1***a). They also lack the positively charged DNA-contacting residues found in the most sequence-specific zinc fingers in the array (consistent with an electrostatic attraction to the negatively charged DNA). We speculate that these bulky, uncharged middle zinc fingers fail to bind DNA strongly and may act more like a linker between the more strongly binding zinc fingers found upstream and downstream.

### Motif 7 represents a binding mode favored by the B allele of PRDM9

We next explored whether PRDM9 binding peaks containing different motifs might associate with different recombination outcomes. We observed a much lower mean recombination rate (***HapMap,*** 2007) around Motif 7 peaks, not explained by differences in PRDM9 binding enrichment, promoter overlap, repeat overlap, or H3K4me3 enrichment (***Figure 1***d, *Figure 1*-S4).

Previous work (***Pratto et al., 2014***) found no evidence of different binding specificities between the A and B alleles of PRDM9, in terms of distinct hotspots. Nonetheless, we hypothesized that Motif 7 might be a partially B-allele-specific motif underrepresented in LD-based recombination maps (***HapMap, 2007***), which are dominated by historical recombination events from the more common A allele of PRDM9. To test this, we searched for our motifs in DSB hotspots unique to an individual with an A/B PRDM9 genotype, then compared these to DSB hotspots found in homozygous A/A individuals (***Pratto et al., 2014***). We found that Motif 7 is 20% enriched in A/B-only hotspots relative to A/A hotspots, while all other motifs are found in fairly similar proportions between the two sets (***Figure 1***d). DSB hotspots containing Motif 7 also have lower relative DMC1 enrichment values in A/A relative to A/B testes (***Figure 1***-S4; ***Pratto et al., 2014***). Motif 7 also resembles, but extends, a motif present in A/B-only hotspots (***Figure 1***-S4; ***Pratto et al., 2014***). Therefore, the B allele binds Motif 7 with greater affinity than does the A allele, demonstrating distinguishable binding preferences between these alleles, which differ at a single DNA-contacting amino acid in ZF5 (***Baudat et al., 2010***).

### PRDM9 binding depends both on sequence and chromatin context

To examine how the primary DNA sequence affects the probability of PRDM9 binding, we identified matches to each of our motifs genome-wide using FIMO (***Bailey et al., 2015***). Although the probability of overlapping a PRDM9 binding peak increases linearly with motif match score, even the strongest 0.1% of motif matches have only a 50% chance of overlapping a binding peak (see ***Figure 2***-S2a). Given that binding cannot be reliably predicted by even this multivariate motif score alone, it must be influenced by the wider sequence and chromatin contexts of each motif match.

**Figure 2.**
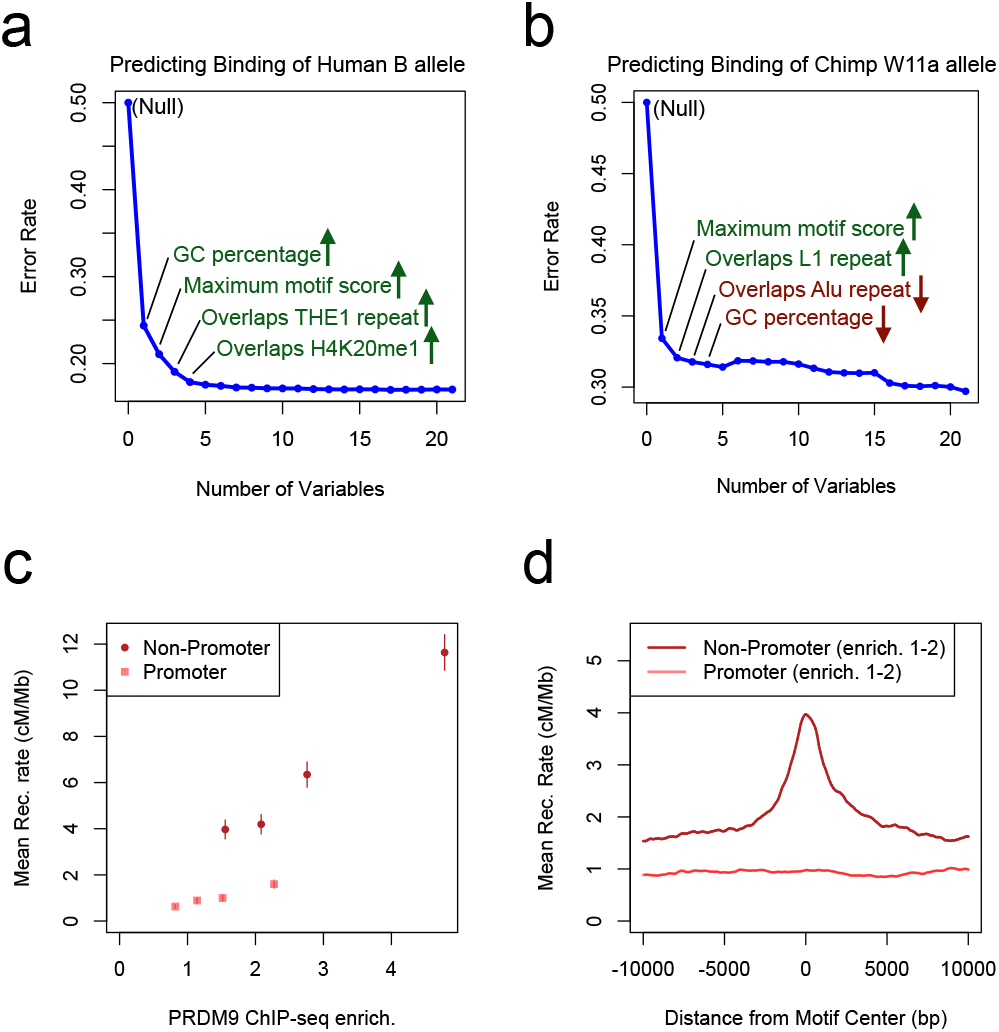
Factors predicting PRDM9 binding. **a**: A logistic regression model was trained on even sets of bound and unbound 100-bp bins across the autosomes for the human PRDM9 B allele ChIP-seq dataset, with 21 genomic and epigenomic annotations as explanatory variables. This plot shows the decrease in error rate on a held-out test set as each new feature is added by forward selection, with the identities of the first four ranking features labeled alongside. Arrows indicate the direction of the effect (green up arrows: positive association with binding; red down arrows: negative association with binding). “Motif Score” refers to the maximum FIMO (***Bailey et al., 2015***) motif score for any of the 7 human motifs in each bin. **b**: As in a, but for the chimp PRDM9 W11a allele ChIP-seq dataset. “Motif Score” refers to the maximum FIMO motif score for the chimp motif (see ***Figure 4***) in each bin. **c**: Mean HapMap recombination rates are reported for promoter (pink squares) and non-promoter (red circles) human PRDM9 peaks split into quartiles of PRDM9 enrichment (filtered to not overlap repeats or occur within 15 Mb of a telomere; error bars represent two standard errors of the mean). Both median enrichment values and recombination rates are greater for non-promoter peaks, even in overlapping ranges of enrichment. **d**: Mean recombination rate in 20-kb windows centered on bound motifs, for promoter (pink) and non-promoter (red) peaks further filtered only to include peaks with PRDM9 enrichment values between 1 and 2 (smoothing: ksmooth bandwidth 200). **Figure 2-Figure supplement 1**. See Figure Supplements **Figure 2-Figure supplement 2**. See Figure Supplements

**Figure 3.**
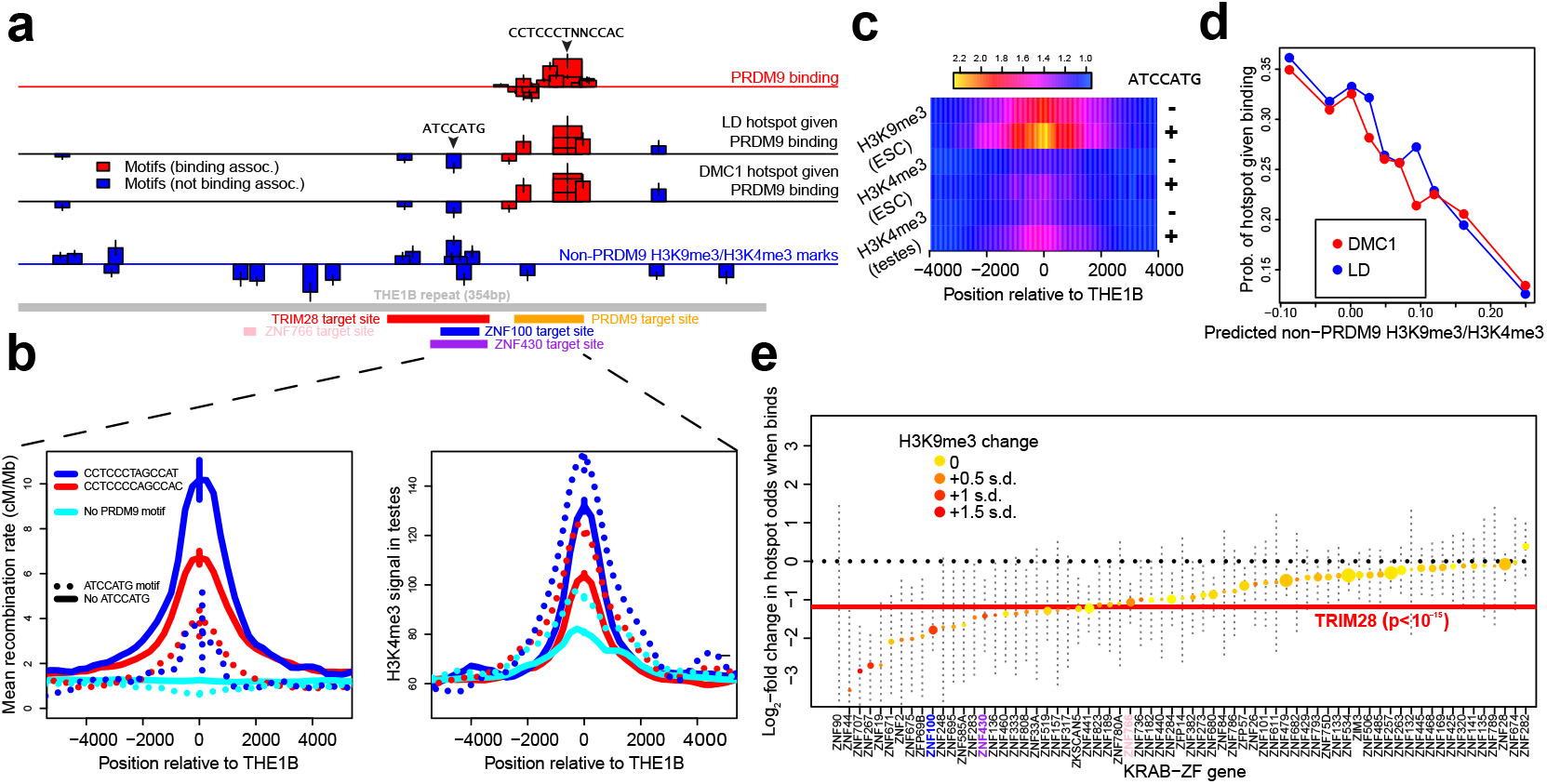
Inf uences on recombination in *cis* downstream of PRDM9 binding. **a**: Analysis of THE1B repeats shows the positions along the THE1B consensus (bottom, grey) of motifs influencing PRDM9 binding (top row), motifs influencing recombination hotspot occurrence at bound sites (middle two rows), and motifs influencing H3K4me3/H3K9me3 in testes and somatic cells (bottom row). Rectangle widths show motif size, and heights show log-odds-ratio or effect size (2 standard errors delineated). Rectangles below the lines have negative effects. Motifs associated with PRDM9 binding are in red; others in blue. Binding motifs for labeled proteins are at the plot base. **b**: Left plot shows LD-based recombination rates around the centers of THE1B repeats containing different approximate matches to the PRDM9 binding motif CCTCCC[CT]AGCCA[CT] (colors) and the motif ATCCATG (lines dotted if present). Right plot is the same but shows mean H3K4me3 in testes. ATCCATG presence reduces recombination and increases H3K4me3. **c**: Impact of ATCCATG presence (+) or absence (-) on normalized enrichment values around the centers of THE1B repeats, of H3K4me3 and H3K9me3 in different cells (labeled pairs of color bars, normalized to equal 1 at edges). H3K9me3 shows the strongest signal increase. **d**: Predicted non-PRDM9 H3K9me3/H3K4me3 versus probability DMC1-based or LD-based hotspots occur at PRDM9-bound sites. For the x-axis repeats were binned according to an additive DNA-based score, using the bottom row of part A and the combination of motifs they contained. **e**: Estimated impact on whether a hotspot occurs of co-binding by individual KRAB-ZNF proteins (labels; ***Imbeault et al., 2017***) near a PRDM9 binding peak. For each KRAB-ZNF protein, after peak filtering, a GLM was used to estimate the impact of KRAB-ZNF binding (binary regressor) on hotspot probability. We show the estimated log_2_-odds, with 95% CI’s. Colors indicate H3K9me3 enrichment increase at co-bound sites. Horizontal line shows the results for TRIM28. Features below the horizontal dotted line have a negative estimated impact on downstream recombination. **Figure 3-Figure supplement 1**. See Figure Supplements **Figure 3-Figure supplement 2**. See Figure Supplements **Figure 3-source data 1**. Detailed information on all THE1B motifs. file: http://www.stats.ox.ac.uk/~altemose/THE1B_Motifs.xlsx

**Figure 4.**
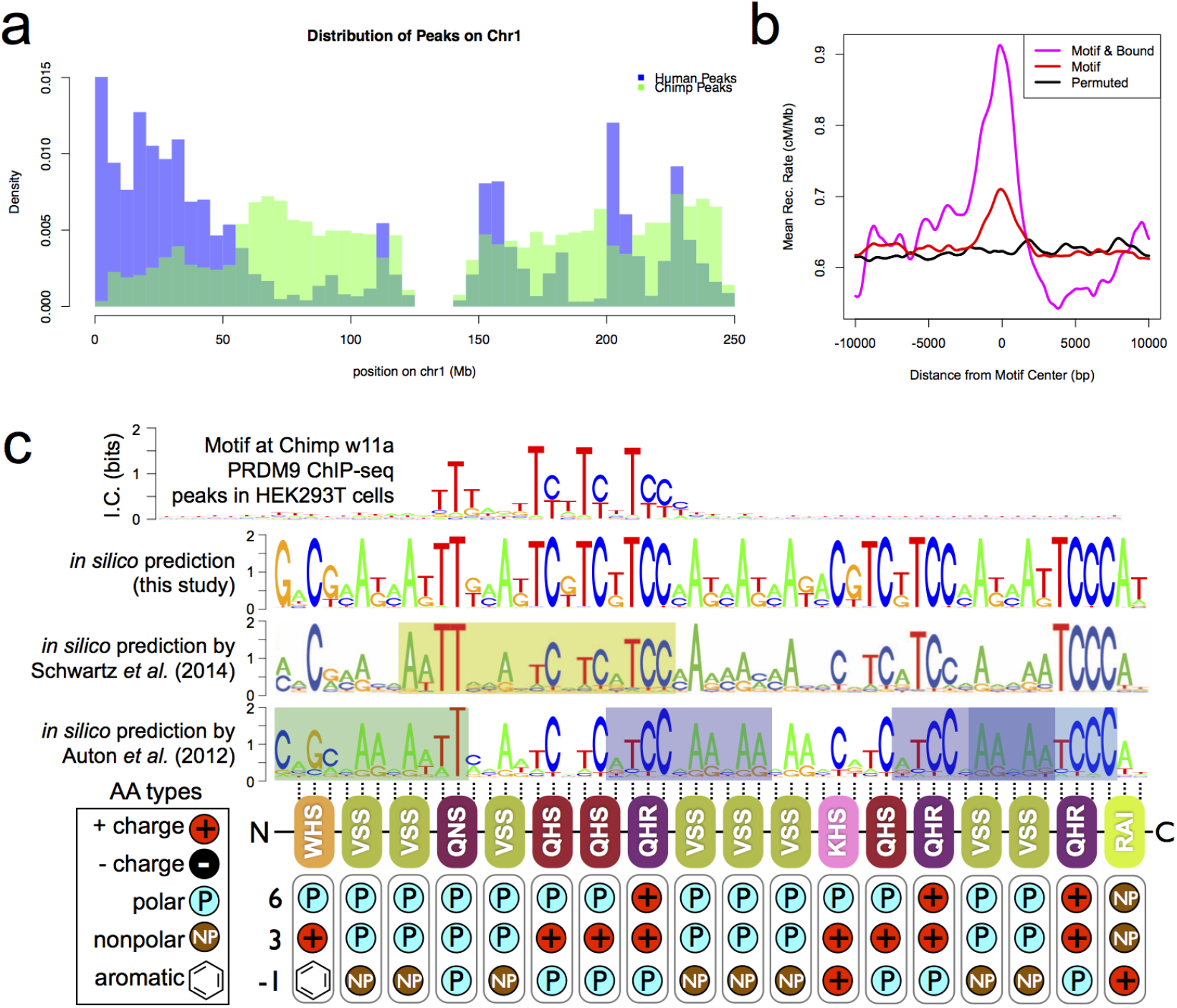
A novel Chimp PRDM9 binding motif. **a**: Comparison ofthe number of human (B allele; blue) and chimp (W11a allele; semitransparent green) PRDM9 ChIP-seq peaks in1-Mbbins across human Chr1. Consistent with their different motifs and other binding predictors (***Figure 2***), we observe very different binding landscapes across the genome. **b**: Profile plot showing the mean chimp recombination rate centered on either the strongest ~40,000 chimp motif matches in the genome (red), the subset ofthose matches that are among our binding peaks (5584 motifs, magenta), or a set of positions shifted at random uniformly in the range [-60000,60000] from the motif match locations (black), as a measure of background (smoothing: ksmooth, bandwidth 500). **c**: Top: the only non-degenerate motif returned by our motif-finding algorithm when trained on the top 5000 chimp PRDM9 peaks ranked by enrichment. Although the motif extends 53-bp, only a 17-bp core region shows high specificity, and this region overlaps and matches *in*-*silico* binding predictions (***Persikov et al., 2009; Persikov and Singh, 2014; Schwartz et al., 2014; Auton et al., 2012***), in particular a submotif (highlighted in yellow) shown to be common to many chimp PRDM9 alleles (reproduced from ***Schwartz et al., 2014***). Zinc finger residuesat DNA-contacting positions(labeled −1,3, 6)are illustrated below each zinc finger position, classified by polarity, charge, and presence ofaromatic side chains. In contrast to the human B allele, this chimp allele has 18 instead of 12 canonical zinc fingers, and they differ in amino acid types at the DNA-contacting positions.

To identify factors that predict whether any given region of the genome will be bound by PRDM9 at fine scales, we built a generalized linear model to predict the bound/unbound status of a set of 100-bp bins across the autosomes given a wide range of annotated genetic and epigenetic features (***Figure 2***a). We report the classification accuracy of the model on a held-out test set, successively adding new variables. The feature that provided the greatest decrease in classification error (from 50% to 24%) was GC content (positive), followed by the maximum motif FIMO score within each bin (positive), THE1 repeat overlap (positive), and H4K20me1 peak overlap (positive; ***Figure 2***a). With only these four features, the model achieves 82% classification accuracy on a held-out test set, compared to a null expectation of 50%, and none of the other variables when added individually or in combination produce significant further improvements in classification (83% accuracy with all 21 features considered; ***Figure 2***-S1). Recombination rates and human PRDM9 motifs are known to be enriched in THE1 elements (***Myers et al., 2005***), but the association with H4K20me1 binding has not been previously described. This mark is associated with DNA replication, DNA damage repair, and chromatin condensation (***Jørgensen et al., 2013***), and thus it may correlate with DNA accessibility to PRDM9 binding.

### Human PRDM9 is able to bind promoters genome-wide

A study in mice has shown that in the absence of PRDM9, DSBs localize to active promoters marked with H3K4me3. It has been suggested that PRDM9 may serve to provide alternative H3K4me3 sites to compete with and direct recombination away from promoters (***Brick et al., 2012***). However, our ChIP-seq data revealed that, surprisingly, of the 12,982 protein-coding genes with H3K4me3 surrounding their Transcription Start Site (TSS) in our untransfected cells (p<10^-5^), 81% have a PRDM9 binding peak center within 500 bp of the TSS, compared to only 6% expected by chance overlap.

Our power to detect binding at promoters is likely increased due to their overrepresentation among ChIP-seq reads (***Figure 2***-S2, ***Jain et al., 2015***). However, we see no promoter ChIP-seq enrichment for the chimp PRDM9 W11a allele, which does not bind GC-rich DNA (see below; only 3% of promoters are within 500 bp of a chimp PRDM9 peak, versus 9% expected by chance overlap). Furthermore, motif identification at human PRDM9’s promoter binding sites revealed the expected binding motifs at similar frequencies to non-promoter peaks, except for a 2-fold enrichment of Motif 7. Interestingly, Motif 7 is also the B-allele enriched motif, so PRDM9’s promoter affinity might also differ between common human alleles. We suggest that these motifs, together with accessible chromatin, allow the observed weak but consistent PRDM9 binding to these regions (***Figure 2c***,***Figure 2***-S2), which tend to have lower mean enrichment estimates across a range of motif FIMO scores (***Figure 2***-S2). Thus, the human B allele of PRDM9 can and does consistently bind to promoters, but more weakly than to non-promoter regions. A recent study of PRDM9 binding in mouse testes (***Grey et al., 2017***) found that mouse PRDM9 can also be present at a small number of promoter regions, but this recruitment depended on Spo11 (absent in HEK293T cells) and was hypothesized *not* to involve PRDM9’s zinc fingers. Therefore, different alleles of PRDM9 interact with promoters in different ways, and, as described below, recombination continues to be strongly suppressed at promoters, even if PRDM9 can bind them.

### Recombination outcomes depend on genomic context

Across all motifs, peaks overlapping promoters show little or no increase in recombination rate above the background rate of 1.1 cM/Mb (***Kong et al., 2002;*** see ***Figure 2***d). This effect cannot be explained by the weaker PRDM9 enrichment that we observe at promoter peaks; for similar enrichment values (strongly bound promoters versus weakly bound non-promoters), promoter peaks have much lower recombination rates and DMC1 enrichment (see ***Figure 2***, ***Figure 2***-S2). Although there is widespread human PRDM9 binding to promoters, PRDM9 seems utterly unable to induce recombination at these sites; however, in the absence of PRDM9, DSBs localize to promoters in mice (***Brick et al., 2012***). Thus, if competition with other PRDM9-bound loci explains why PRDM9 eliminates recombination at promoters, this competition must act downstream of PRDM9 binding and is not dependent only on H3K4me3 level. For example, promoters might contain a local chromatin environment that is much less favorable for DSB formation than other binding sites. In mice, *in vivo* recombination hotspot sites favor motif positions with lower H3K4me3 levels than genomic background (***Davies et al., 2016***), and this seems highly concordant with the results we report here.

To specifically examine the effect of local chromatin marks on recombination outcomes at PRDM9-bound sites, we annotated our binding peaks with whether they overlap ChIP-seq peaks reported for 9 histone variants or modifications reported by the ENCODE project: H3K9me1, H3K9me3, H3K9ac, H2az, H3K27ac, H3K27me3, H3K36me3, H3K79me2, and H4K20me1 (***ENCODE, 2012***). Because these data were collected in a different human cell line (K562), we can regard them only as an imperfect proxy for true chromatin states in HEK293T cells and in spermatocytes, relying on the fact that in comparisons across cell types, many or most chromatin mark locations are similar (***ENCODE, 2012***). Most of these chromatin marks are associated with active enhancers, promoters, and gene bodies, with the exceptions of H3K9me3 and H3K27me3, which mark heterochromatin (***ENCODE, 2012***). Interestingly, mean recombination rate decreases significantly across all chromatin marks tested (95% C.I. ranges −6% to −63%) suggesting repression as a dominant impact of chromatin modifications. The sole exception is H3K27me3, whose peaks shows a 28% increase above the mean rate for all peaks (95% C.I. 17-40%; see ***Figure 3***-S1a). That is, conditional on binding strength, PRDM9 binding sites overlapping facultative heterochromatin regions, which are typically transcriptionally repressed, appear to be more likely to become recombination hotspots. On the other hand, both active chromatin environments, and constitutive heterochromatin, consistently show reductions in hotspot probability. It is obviously challenging to conduct a comprehensive exploration of whether – and exactly how - these relationships might be causal or correlative, and the extent to which the chromatin environment reflects *cis* or *trans* factors. However, we were able to explore these questions in detail within a collection of hotspots that collectively account for around 5% of human recombination.

### Analysis of THE1B repeats reveals non-PRDM9 motifs for recombination hotspots

Although only a subset of PRDM9-bound sites in the genome become recombination hotspots, the only specific mammalian sequence feature so far identified as influencing either PRDM9 binding, or downstream recombination events, is the PRDM9 binding motif itself. Thus, it is uncertain which factors prevent or promote hotspot occurrence, whether these act in *cis* or *trans*, and what these might be.

One approach to address these questions is to search for sequence motifs that might influence PRDM9 binding and subsequent hotspot formation. Identified motifs are likely to have a causal influence, so they can help address whether *e.g*. particular histone modifications associated with those motifs have a genuinely causal influence. Although in general motif identification is complicated by hotspot background heterogeneity, one family of retrotransposon elements, called THE1B repeats, contribute a large fraction of human A and B-allele recombination (4.6% measured by DMC1 mapping; ***Pratto et al., 2014***) on a relatively homogenous genetic background. PRDM9 binds directly to a subset of THE1B repeat copies containing matches to its target motif (***Figure 3***a), in a known region of the repeat (***Myers et al., 2008***). Of 20,696 autosomal THE1B repeat copies, 21% (4,392) overlap our PRDM9 ChIP-seq peaks. These PRDM9-bound copies fully explain THE1B enrichment among recombination hotspots identified by DMC1 mapping (1155 hotspots; p<10^-15^ by FET; odds ratio 10.8; ***Pratto et al., 2014***), or LD mapping (1209 hotspots; ***HapMap, 2007***). Unbound THE1B repeats do not show significantly greater overlap with DMC1 hotspots than expected by chance (p=0.18 compared to a null set of THE1B repeat positions right-shifted 5 kb). Nevertheless, many strongly bound THE1B repeat copies still do not become hotspots.

Because THE1B repeats are spread throughout the genome and share highly similar sequences, perturbed by random mutations, we were able to precisely dissect the impact of particular sequence motifs occurring in subsets of these repeats on PRDM9 binding, and on downstream DSB formation (as measured by DMC1 mapping) and crossover activity (as measured by LD mapping). We first examined the relationship between PRDM9 binding and broad-scale recombination rate by partitioning THE1B repeats into quintiles of increasing recombination rate in the surrounding 1 Mb in males (independently measured by ***Kong et al., 2002;*** excluding the 20-kb region surrounding each repeat to avoid direct biasing of results). Peaks in mean DMC1 heat occur at THE1B repeats in all cases, but peak height increases strongly with broad-scale heat for both telomeric and non-telomeric regions (***Figure 3***-S2). Therefore, in broad “hotter” regions, more double-strand breaks occur, completely independently of the local sequence (which is similar in THE1B repeats genome-wide). This is not unexpected, given previous observations of similar broad-scale recombination rate patterns among differing PRDM9 alleles (***Pratto et al., 2014***). Although broad-scale correlations have unknown causes, one possible explanation *a priori* is that general broad-scale accessibility to PRDM9 binding differs between hot and cold regions, in a manner shared across alleles. To test this we also examined mean H3K4me3 signals in testes in the same way (***Figure 3***-S2), which should be reduced if PRDM9 binding is depressed in even a subset of colder regions. Strikingly, this revealed no difference whatsoever between hot and cold regions, or between telomeric and non-telomeric regions, implying >10-fold differences in mean recombination rate occur without any change in mean H3K4me3 enrichment at THE1B repeats. This proves that at least in human males, broad-scale recombination control operates without impacting PRDM9’s ability to bind and deposit H3K4me3, a property observed previously for elevated male recombination in telomeres (***Pratto et al., 2014***). Therefore, DSB formation has an additional layer of regulation, downstream of H3K4me3 deposition by PRDM9. The different recombination rates observed between the two sexes do not, then, necessarily imply differential binding by PRDM9.

Motivated by these broad-scale results, we now tested for *local* impacts of particular sequence motifs occurring in subsets of THE1B repeats on both PRDM9 binding, and downstream DSB formation (as measured by DMC1) and crossover activity (as measured by LD patterns). We used conditional association testing to identify collections of motifs that independently correlate with PRDM9 binding or recombination (see Methods and Materials).

Seventeen distinct motifs (***Figure 3***a) were found to influence PRDM9 binding in THE1B elements (***Figure 3***-source data 1). All map within the predicted PRDM9 binding region and span the entire region, confirming that all of PRDM9’s zinc fingers are involved in binding to THE1B copies. Motifs promoting PRDM9 binding were consistently associated with higher H3K4me3 in testes and increasing hotspot probability (Methods and Materials, ***Figure 3***-source data 1, ***Pratto et al., 2014***), so the same motifs operate *in vivo*. Importantly for the results described below, binding of PRDM9 does not associate strongly with any sequence motifs outside the directly bound region, so it might act as a local “pioneer” protein at least on this background, despite results in mice (***Grey et al., 2017***).

We then independently tested for the presence of motifs influencing recombination hotspot formation (requiring association with both DMC1 and LD-based hotspots) *conditional* on PRDM9 binding in HEK293T cells. We identified an initial 7 such motifs (Methods and Materials; ***Figure 3***a; ***Figure 3***-source data 1). Only three of these map within the PRDM9 binding region and correspond to stronger/weaker PRDM9 binding. The remaining four “non-PRDM9” recombination-influencing motifs show no association whatsoever with PRDM9 binding in HEK293T cells, and map well outside the PRDM9 binding motif (***Figure 3***a). The strongest signal is for the motif ATCCATG (joint p=2.8×10^-9^ for LD-hotspots, OR=0.32; p=2.5×10^-6^ for DMC1 hotspots), whose presence within a THE1B repeat produces a dramatic reduction in the surrounding recombination rate at PRDM9-bound THE1B repeats (***Figure 3***b). ATCCATG presence also reduces the local recombination rate below background in repeats containing no PRDM9 target motif and not bound by PRDM9, implying an impact beyond the THE1B repeat itself and not dependent on whether PRDM9 can bind the repeat. We examined H3K4me3 signal in testes (from ***Pratto et al., 2014***) around THE1B elements containing, and not containing, the motif ATCCATG, and conditional on the strength of match to the PRDM9 binding motif within the THE1B element (***Figure 3***b), to determine whether it might operate by preventing binding or H3K4me3 deposition in early meiosis. Strong H3K4me3 enrichment specific to THE1B repeats containing PRDM9 binding motifs occurred regardless of whether “ATCCATG” was present. Therefore, this motif does not suppress PRDM9 binding but instead acts downstream. In fact, presence of the modifier motif ATCCATG actually modestly *increased* the H3K4me3 signal, something returned to below. Similar results were observed for the other three non-PRDM9 recombination-influencing motifs.

### Motifs influencing local chromatin states in somatic and meiotic tissue types occur throughout THE1B repeats

To better understand how these motifs might functionally operate, we also performed independent *de novo* motif finding to identify motifs within THE1B elements associating with the occurrence of 15 previously identified chromatin states, and individual histone modifications (p<2.5×10^-8^, significant after Bonferroni correction), across each of 125 somatic cell types (***Kundaje et al., 2015***). This identified rich information: 67 clusters of similar motifs, collectively showing association signals for 8 chromatin states (***Figure 3***-source data 1), and spanning all 125 cell types. It is perhaps surprising that such a rich diversity of motifs is identified to (presumably) influence chromatin state between THE1B repeat copies, given that the THE1B sequence is only around 350 bp in size, although some such influences are subtle.

Strikingly, the motif ATCCATG is that most strongly positively associated with the “heterochromatin” state among all 2,021 seven-mers commonly present within THE1B repeats. Association with heterochromatin, marked by enriched H3K9me3, occurred across >50% of all ROADMAP-annotated cell types, with strongest signals observed in embryonic stem cells. Direct examination of histone modifications (Methods and Materials) revealed a strong localized increase in H3K9me3 within THE1B repeats containing ATCCATG (***Figure 3***c). More surprisingly a weak, but significant, increase in H3K4me3 signal (p=7.5×10^-13^) was also seen, even though this modification is more generally associated with active chromatin regions including promoters. The same weak H3K4me3 peak was also seen in testes, after restricting analysis to THE1B repeats not bound by PRDM9, indicating this modification operates fully independently of PRDM9, and explaining how the H3K4me3 signal also increases with ATCCATG presence when PRDM9 does bind. This weak increase might reflect genuine partial co-occurrence of the two marks at the same locus (but possibly on different alleles, or in different cells), or in theory it could be explained by non-specificity of experimental antibodies for these two histone modifications.

We reasoned that we might more generally exploit the subtle H3K4me3 signal elevation (whatever its underlying cause) as a potential marker also of H3K9me3 elevation in germline tissues, by examining H3K4me3 in testes. We performed *de novo* motif finding to identify PRDM9-independent motifs associating with H3K4me3 in THE1B repeats definitively not bound by PRDM9 (Methods and Materials). This identified eighteen 7-bp motifs significantly associated with non-PRDM9 H3K4me3 (after Bonferroni correction, ***Figure 3a***). The motif ATCCATG remained the most strongly associated (p<10^-25^). The additional motifs occurred throughout the THE1B repeat, but notably eight clustered around this strongest signal.

Confirming that these motifs also predict H3K9me3 levels, we observed almost perfect positive correlation (r=0.93) between H3K4me3 signal strength in testes and H3K9me3 (as well as H3K4me3) in, for example, particular ROADMAP ESC cell-lines (***Figure 3***-S1d). 14 of the 18 motifs showed association with heterochromatin (p<2.5×10^-8^), in at least one cell type. Therefore, this represents a set of motifs for both H3K9me3 and H3K4me3, broadly observable across somatic cells and (at least for the latter mark) testes also, and so we refer to this set as non-PRDM9 H3K9me3/H3K4me3 motifs.

### Non-PRDM9 H3K9me3/H3K4me3 motifs completely coincide with recombination suppressing motifs

In addition to the top-scoring motif ATCCATG, many or all of the other 17 motifs for non-PRDM9 H3K9me3/H3K4me3 evidently impact meiotic recombination, and in the opposite direction. All four of the initial non-PRDM9 recombination-influencing motifs we found *de novo* overlap at least one of these 18 motifs (***Figure 3***a). In a joint test for association of the expanded set of 18 motifs with the occurrence of meiotic recombination hotspots given PRDM9 binding, their estimated effects on H3K4me3 were linearly correlated with both DMC1 and LD-based hotspot status, but with an effect direction opposite to that for H3K4me3 (***Figure 3***c; ***Figure 3***-source data 1; p<0.00036 in both cases). There was no impact on PRDM9 binding (p=0.25). Summing these motif influences to produce a score for each THE1B repeat using only its DNA sequence, we see >3-fold difference in the probability of observing a recombination hotspot across PRDM9-bound THE1B copies between the top and bottom 10% quantiles of the score (***Figure 3***d). Given we are only able to examine the region within each hotspot corresponding to the 354 bases of the THE1B element, it is likely this underestimates the true impact of local sequence on whether hotspots occur or not, and the 18 motifs we find collectively cover around 1/3 of the total THE1 bases near the hotspot center.

Notably, the *de novo* analysis identified many more motifs influencing histone-defined chromatin states in ROADMAP-studied cell types, including the binding targets of two proteins DUX4 and ZBTB33 previously shown to bind toTHE1B elements, with DUX4 showing strong expression in testes (***Young et al., 2013; Wang et al., 2012***). However, only those motifs associated with heterochromatin, and H3K9me3/H3K4me3, in somatic cells overlapped our new meiotic recombination associated motifs. Thus, only a particular subset of chromatin modifications correspond to suppressed recombination in THE1B repeats.

Overall, this analysis of thousands of human hotspots reveals that in *cis*, it is not simply PRDM9 binding that influences whether hotspots occur. Multiple sequence motifs exist that do not prevent PRDM9 binding, but instead modify the average amount of recombination that occurs *downstream* of binding, up to >2-fold for a single motif. Given this diversity even within THE1B-centered hotspots, completely different motifs might operate to modulate recombination activity in other hotspots, either centered in different repeats or in non-repeat DNA. In contrast to this complexity, examination of histone modifications reveals a common signature across motifs, with strong alterations in the specific histone marks H3K9me3 and weaker signals for H3K4me3. These occur in the opposite direction to recombination effects, and particularly strongly in embryonic stem cells. This suggests that the mechanism of action across motifs might share fundamental similarities. Both H3K4me3 and H3K9me3 marks correlate negatively with recombination across all human hotspots (***Figure 3***d; ***Figure 3***-S1), and reduced levels of non-PRDM9 H3K4me3 within hotspots has been observed in mice (***Brick et al., 2012; Davies et al., 2016***).

### KRAB-ZNF binding and TRIM28 recruitment suppress recombination

The large class of human KRAB-ZNF genes represent an obvious set of motif-binding candidates that might explain H3K9me3 deposition within THE1B repeats and more broadly. In many such genes, the KRAB domain recruits TRIM28, which in turn recruits histone modifying proteins including SETDB1, which lead to H3K9me3 deposition on nearby nucleosomes. We therefore examined recent data (***Imbeault et al., 2017***) measuring genome-wide binding of 222 KRAB-ZNF proteins in humans, and sites where TRIM28 is present in embryonic stem cells, for overlap with THE1B repeats (Methods and Materials). Three such proteins (ZNF100, ZNF430 and ZNF766), as well as TRIM28, are enriched for binding in THE1B repeats (***Imbeault et al., 2017***) and also associate genome-wide with H3K9me3 deposition. We identified binding motifs for each within THE1B repeats. Strikingly, ATCCATG overlapped the second most significant motif for TRIM28 recruitment, and additional motif analysis for TRIM28 revealed a large (51-bp) motif, fully spanning a cluster of eight motifs associated with H3K9me3/H3K4me3 and recombination rate, and presumably representing the binding target of one or more as-yet-unknown KRAB-ZNF protein(s). The three specific ZNF proteins also all bind sites overlapping those implicated in impacting H3K9me3/H3K4me3 and meiotic recombination, two in the same region as the TRIM28 motif, but with differing sequence specificity (***Figure 3***a). Thus, while not all human KRAB-ZNF proteins have yet been characterized, those that bind THE1B repeats consistently operate to reduce recombination, and TRIM28 recruitment can explain the strongest signals we see.

Across all our PRDM9 binding peaks, 36.5% fall within 500 bp of a binding site of at least one of the KRAB-ZNF proteins with available data (***Imbeault et al., 2017***), suggesting that such repression might be important in regulating recombination more generally. To test this, we individually analyzed the KRAB-ZNF proteins with at least 30 instances of a KRAB-ZNF binding peak occurring near a PRDM9 binding peak (after excluding DNase HS regions and promoters, which are often bound by multiple different proteins), for their effect on whether a hotspot occurs at these PRDM9 binding peaks (Methods and Materials). This revealed a universal negative trend (***Figure 3***e) typified by a >2-fold reduction in recombination locally atTRIM28-marked sites genome-wide, with every gene except one (ZNF282, which was non-significant) inferred to reduce hotspot odds. Binding of almost all KRAB-ZNF genes correlated positively with H3K9me3. Those genes with strongest H3K9me3 enrichment showed the strongest suppression of recombination locally (***Figure 3***e).

Together, our results indicate a mechanism of *cis* recombination repression affecting thousands of human PRDM9 binding sites. Binding of KRAB-ZNF proteins to specific sequence motifs within or nearby the PRDM9 binding site, followed by TRIM28 recruitment and H3K9me3 deposition, universally acts to strongly repress local recombination, at least sometimes without preventing PRDM9 binding or H3K4me3 deposition. In a conditional analysis to predict PRDM9 binding (***Figure 2***a,b), we found that the H3K9me3 mark associates negatively with the binding of human PRDM9 but positively with the binding of chimp PRDM9 in transfected human cells, although it was not among the top predictors of PRDM9 binding for either allele (***Figure 2***-S1). Many KRAB-ZNF genes bind to specific sets of retrotransposon repeats (THE1B repeats represent one example), so this repressive mechanism is likely to act to reduce recombination around many particular repeats.

### Comparing chimp PRDM9 and human PRDM9

In order to better understand the epigenetic predictors of binding, we next sought to explore the properties of a PRDM9 allele very different from the human B allele. We chose the chimpanzee reference allele (W11a, or Pan.t4,8,12,16), measured to be at roughly 13.4% frequency in wild chimpanzees (***Auton et al., 2012; Schwartz et al., 2014***). An LD-based genetic map of chimp recombination failed to identify definitive motifs at recombination hotspots, which tend to be weaker at the population level than those found in human populations (***Auton et al., 2012***). The chimp allele differs from the human B allele in having 18 canonical zinc fingers, as opposed to 12, with different predicted binding preferences.

*De novo* peak calling at the same thresholds (p<10^-6^) yielded 247,717 total chimp PRDM9 peaks, higher than the number observed for the human B-allele. Only 2% of chimp peak centers occurred within 1 kb of a human peak center, below chance expectation, so their ZF arrays have very different binding preferences (***Figure 4***). At broad scales, peaks for the human allele tend to be overrepresented in GC-rich regions and promoters, but peaks for the chimp allele show the exact opposite pattern, with overrepresentation in AT-rich regions, outside promoters. Because we have increased power to detect binding in these regions and have shown that the magnitude of human PRDM9 binding enrichment is lower at promoters, the lack of chimp PRDM9 binding sites in these regions is consistent with chimp PRDM9 failing to bind even weakly.

After running the same motif-finding pipeline used for the human allele, we identified a 17-bp, somewhat AT-rich motif, found at 60% of binding peaks and highly centrally enriched within peaks (***Figure 4***c). We compared this motif with published *in silico* binding predictions for this allele (***Auton et al., 2012; Schwartz et al., 2014***) and found a close match in the central region of the predicted motifs.

We plotted the chimp recombination rate (***Auton et al., 2012***) around the strongest ~40,000 FIMO matches for this motif as well as at the subset of 5,584 of these sites bound in our transfected cells. A modest increase in local recombination rate is visible at motif matches, with a much larger increase for those which are bound in our assay (***Figure 4***). Hence our binding sites overlap true chimp hotspots, but the association between binding and the population recombination rate is much smaller than for the human allele. This may owe to the fact that chimps possess a much greater diversity of PRDM9 alleles in their population (***Auton et al., 2012***), producing a large union set of hotspots, each of which only accounts for a small fraction of recombination in the population. Interestingly, the chimp PRDM9 motif almost exactly overlaps a subregion of the *in silico* binding motif that was identified as being common to many different chimp alleles (***Schwartz et al., 2014***), and we suggest natural selection might be a cause of this remarkable coincidence. In humans, a group of “C-like” alleles strongly bind a common motif also 17-bp long (***Hinch et al., 2011***), and again the zinc-fingers implicated as binding this motif are shared across the otherwise diverse alleles.

### PRDM9 can activate transcription of some genes, including *VCX* and *CTCFL*

Because the human B allele binds promoters, this raises the possibility that PRDM9’s H3K4me3 mark may play a role, whether as an accidental side effect of binding or specifically functional, apart from simply specifying the locations of meiotic DSB breakpoints. We therefore performed RNA-seq in cells transfected with the Human and Chimp alleles, and control cells that were either untransfected, or transfected with a construct containing only the human ZF domain (and incapable of H3K4me3 deposition; referred to as “ZFonly”, illustrated in ***Figure 6***a).

Seven transcripts showed overwhelming evidence of being differentially expressed in Human-transfected cells versus all other samples, all seven being upregulated in the Human sample. Five overlap known genes: *MEG3, ONECUT3, LGALS1, VCX*, and *CTCFL*. Interestingly, the latter two genes are normally expressed only in spermatogenesis. We validated expression induction at these two genes using qPCR (***Figure 5***).

**Figure 5.**
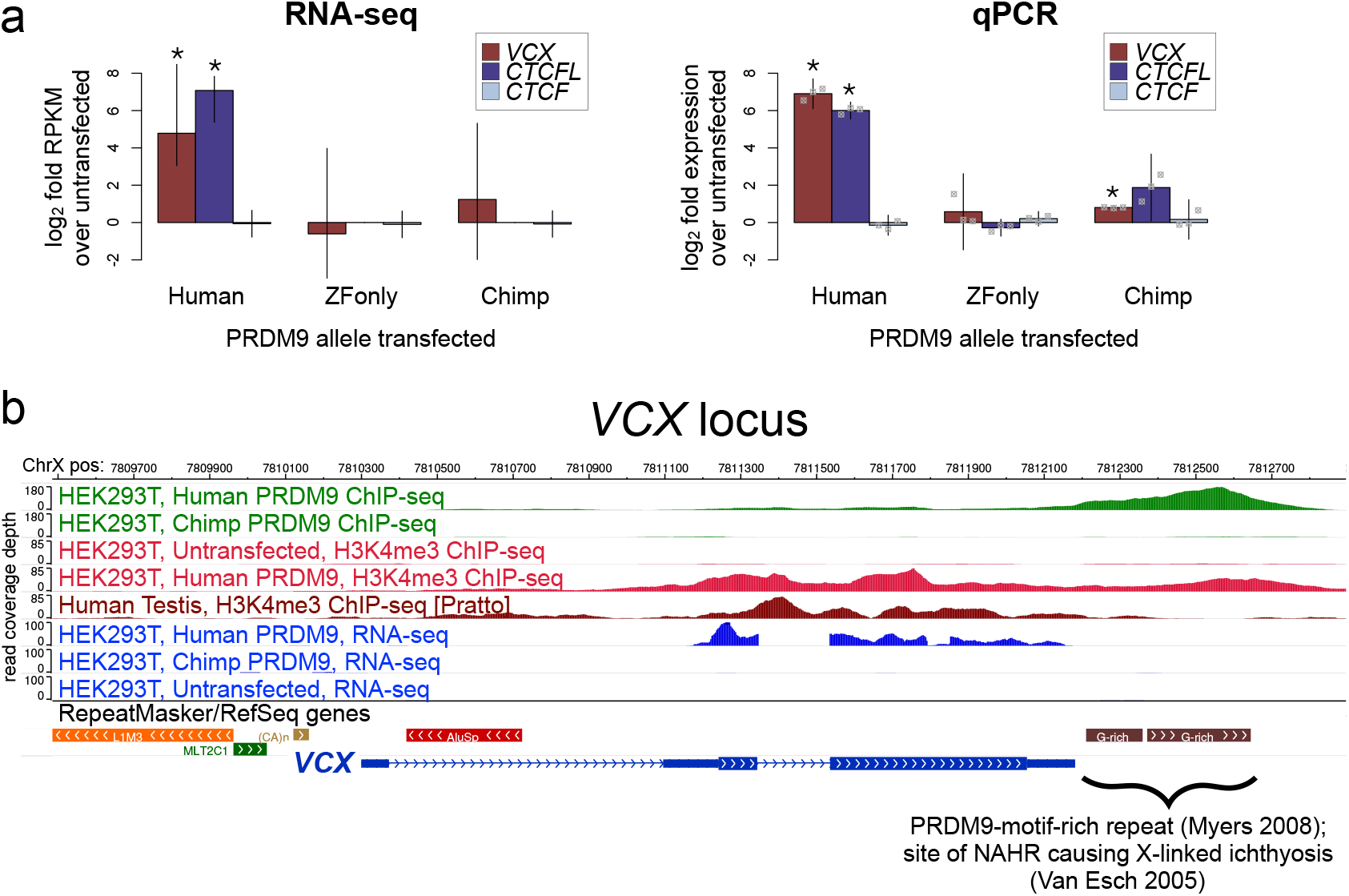
Spermatogenesis-specific genes VCX and CTCFL are activated by Human PRDM9. **a**: left: Bar plots showing the log_2_ fold change relative to untransfected HEK293T cells in computed FPKM values (fragments per kilobase of transcript per million mapped RNA-seq reads) for HEK293T cells transfected with the Human allele, the Chimp allele, or a construct containing only the human Zinc Finger domain, for *CTCFL* and VCX, with *CTCF* as a negative control. Error bars conservatively represent maximum ranges of the ratios given confidence intervals for FPKM values computed by cufflinks (***Trapnell etal., 2012***). Asterisks indicate significant differential gene expression, as reported by CuffDiff (p<0.0001). right: qPCR validation results for the same genes from independent biological replicates. Y-axis values are log_2_ ratios of ΔΔ *C_t_* values for each gene relative to the untransfected sample (normalized to the *TBP* housekeeping gene; see Methods and Materials). Error bars represent 95% confidence intervals from 3 biological replicates (t distribution; gray points represent individual replicate values), and asterisks indicate p<0.001 (one-tailed t test). **b**: A browser screenshot (***Zhou et al., 2011***) from ChrX containing the *VCX* gene with custom tracks indicating ChIP-seq and RNA-seq raw coverage data. Human PRDM9 (green) binds a G-rich repeat near the terminus of *VCX* as well as two loci in the middle of the gene, adding H3K4me3 marks (light red) where none were present in untransfected cells. RNA-seq coverage (blue) spikes in the coding regions in transfected cells, while it is nearly flat in untransfected cells. Testis H3K4me3 coverage (dark red, from ***Pratto et al., 2014***) also increases in the gene body, instead of near the annotated TSS. **Figure 5-Figure supplement 1**. See Figure Supplements **Figure 5-Figure supplement 2**. See Figure Supplements

**Figure 6.**
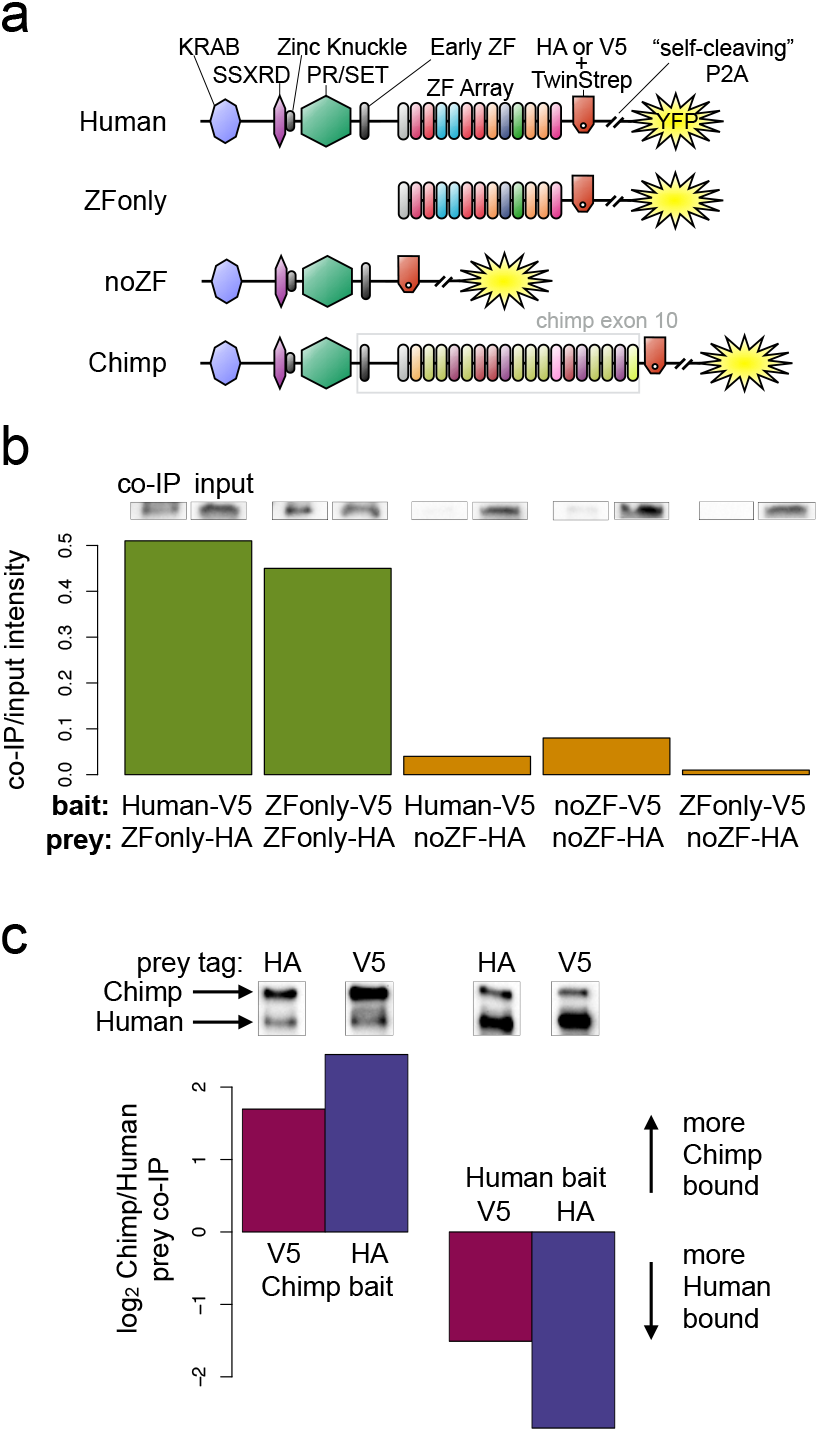
PRDM9 multimer formation is mediated by the ZF domain in an allele-biased manner. **a**: Overview of the different C-terminally tagged PRDM9 constructs used. Both an HA and a V5 version of each construct were generated for co-IP experiments. **b**: Barplot showing the relative intensity of western blot co-IP bands normalized to input bands (from 50-*μ*g of total lysate protein) for each combination of bait and prey constructs. Whenever both bait and prey contain the zinc finger domain (green bars), the co-IP signal is much stronger than when either or both constructs lack a ZF domain (orange bars). See Figures S9 and S10 for complete westerns with mock controls. **c**: Barplot showing the results of competitive co-IP experiments performed in cells transfected with both Human and Chimp as prey (with the same epitope tag) and either Human or Chimp as bait (with a complementary epitope tag). Bars indicate the relative co-IP band intensity for Chimp and Human prey constructs when pulled down with either Chimp or Human bait. When Human is used as bait, more Human prey is pulled down; when Chimp is used as bait, more Chimp prey is pulled down (and this holds for both directions of HA/V5 tagging). **Figure 6-Figure supplement 1**. See Figure Supplements **Figure 6-Figure supplement 2**. See Figure Supplements **Figure 6-Figure supplement 3**. See Figure Supplements

*VCX* encodes a small, highly charged protein of unknown function and has been previously studied for its involvement in PRDM9-related non-homologous recombination events and X-linked ichthyosis (***Myers et al., 2008; Van Esch et al., 2005***). We found that PRDM9 does not in fact bind near the annotated *VCX* Transcription Start Site (TSS), but instead in the middle of the gene and very strongly at a minisatellite repeated series of PRDM9 binding motifs (***Myers et al., 2008***) near the terminus of the gene. PRDM9 adds the H3K4me3 mark throughout the gene’s coding regions in a pattern similar to that seen in testes (***Figure 5***). RNA-seq coverage suggests normal splicing, but use of an alternative promoter that excludes the first, untranslated exon.

*CTCFL* is a variant of *CTCF* expressed exclusively in pre-leptotene spermatocytes. Male knockout mice show greatly reduced fertility due to meiotic arrest (***Sleutels et al., 2012***), and variants at CTCFL influence genome-wide recombination rates in human males (***Kong et al., 2014***). CTCFL may be involved in organizing the meiotic chromatin landscape and regulating the transcription of meiotic genes (***Sleutels et al., 2012***). CTCFL RNA levels increase 28-fold after transfection with the human allele, from a nearly undetectable baseline transcription level (***Figure 5***). PRDM9 binds strongly to a GC-rich repeat near the CTCFL TSS and creates H3K4me3, which is absent in untransfected cells (***Figure 5***-S1). The chimp PRDM9 allele does not bind near the TSS and does not show elevated transcript levels after transfection.

We note that this result does not establish whether human PRDM9 is necessary or sufficient for CTCFL and VCX expression *in vivo*, but still PRDM9 is demonstrably able to trigger the transcription of these genes in a way that depends on the binding of its zinc fingers. Recent work has shown that *Prdm9* expression begins in pre-leptotene cells in mice (***Sun et al., 2015***), concurrent with *Ctcfl* expression (***Sleutels et al., 2012***) and thus supports the possibility that PRDM9 may promote *CTCFL* transcription *in vivo*. The failure of the chimp allele to bind to or activate the expression of human *CTCFL* further suggests that this behavior may not be essential across organisms, although the chimp allele might in principle still bind the *CTCFL* promoter in the chimp genome. Similarly, there is not evidence that human PRDM9 alleles with very different binding preferences, such as the C allele, bind the same promoter. Also notably, the motif bound at the *CTCFL* promoter is Motif 7, so the A and B alleles may bind this locus with different affinities.

46 additional genes showed weaker evidence of being activated by human PRDM9 binding near their annotated transcription start sites, with 44 showing increases, as opposed to decreases, in expression (***Figure 5***-S2). We lack power to detect small changes in gene expression, especially decreases in expression (***Trapnell et al., 2012***). Nonetheless it is likely that effects of similar magnitude to *CTCFL* and *VCX* are quite rare. However, our data do make it clear that PRDM9 binding and trimethylation near a promoter can trigger or enhance gene expression in some cases. Furthermore, this effect on gene expression is not likely to result from PRDM9 binding alone but from its trimethylation activity, given that the ZFonly construct does not trigger expression. This is consistent with recent findings that tethering PRDM9 to other DNA-binding proteins can de-repress gene expression in a context-dependent manner (***Cano-Rodriguez et al., 2016***).

### Multimer formation is mediated primarily by the ZF array

We have studied properties of PRDM9’s zinc fingers in determining DNA binding targets, and the consequences of PRDM9 binding to DNA. At present, DNA binding is the only known role of PRDM9’s ZFs. There is evidence that PRDM9 as a whole can multimerize and that hetero-multimers of the human A and C alleles can bind the sequence targets of either allele and trimethylate surrounding histones (***Baker et al., 2015b***). However, it remains unknown which PRDM9 domain is responsible for this observed multimerization behavior. We sought to determine whether multimerization might involve PRDM9’s ZF domain in any way, given other examples of ZF domains mediating protein-protein interactions (***McCarty et al., 2003; Lee et al., 2007***). To do so, we co-expressed PRDM9 constructs with different ZF domain properties and performed co-ImmunoPrecipitation (co-IP) experiments, thus extending our study from PRDM9’s DNA-binding properties to its protein binding properties.

First, to confirm the ability of the PRDM9 alleles we study here to form multimers (***Baker et al., 2015b***), we performed co-IP experiments with full-length human B-allele PRDM9 constructs differentially tagged with HA and V5 epitopes and co-transfected into HEK293T cells. Following IP against the HA-tagged construct, we detected the V5-tagged construct very robustly; and conversely (***Figure 6***-S1). This is consistent with human PRDM9 binding strongly to itself, as demonstrated previously (***Baker et al., 2015b***).

To narrow the PRDM9 domain(s) responsible for this self-binding behavior, we split the full-length human B-allele PRDM9 cDNA into two pieces: one containing only the C-terminal Zinc Finger domain (the “ZFonly” construct), and one containing everything else (the “noZF” construct), and tagged with HA or V5 as above (illustrated in ***Figure 6***a). We co-transfected these constructs into HEK293T cells, in various combinations with each other and with full-length PRDM9. We found that the full-length human construct and the ZFonly construct localized to the nucleus, but the noZF construct localized throughout the cell, confirming a dominant role for the ZF domain in nuclear localization (***Figure 6***-S3, ***Collin et al., 2013; Wang et al., 2014***).

Interestingly, the ZF domain alone appears to be responsible for most of PRDM9’s self-binding activity (***Figure 6***b). Following co-transfection of noZF-HA and noZF-V5, and despite very high expression levels visible in the input, only a very faint co-IP band is visible in the absence of the ZF array. Because the mock control lane is clean (***Figure 6***-S2a), this band likely reflects a real but weak self-binding capability mediated by the non-ZF portion of PRDM9. In complete contrast, we saw an intense co-IP band when co-transfecting ZFonly-HA with ZFonly-V5. Therefore, the zinc finger domain of one PRDM9 protein can bind strongly to the zinc finger domain of another, while the rest of the protein interacts more weakly.

To confirm this, we co-transfected full-length, V5-tagged human PRDM9 with either noZF-HA or ZFonly-HA. Again, only a very faint co-IP band is visible with the noZF construct, and a very intense band is visible with the ZFonly construct (***Figure 6***b), so the ZFonly construct is sufficient to bind and pull down the full-length construct. This finding replicated in a repeat experiment, and when reversing the direction of the IP-western experiment (***Figure 6***-S2b). Finally, no co-IP band is seen in a negative control where we co-transfected the noZF construct with the ZFonly construct corresponding to the other end of the protein (***Figure 6***b), ruling out an interaction between the ZF domain and the rest of PRDM9 or any interaction between the epitope tags used. Taken together, these results demonstrate that PRDM9 multimerization depends strongly on the ZF array, and much more weakly on the rest of the protein.

Because these multimers were formed inside live cells and lysed in physiological salt concentrations and without DNase digestion, we cannot rule out a role for DNA in potentially mediating this observed interaction between ZF domains. However, a previous study identified PRDM9 multimerization even after benzonase digestion and confirmed the presence of biologically active hetero-multimers *in vivo* (***Baker et al., 2014***). In light of this, our failure to detect clear multimerization after deleting the ZF domain confirms a critical role for PRDM9’s zinc fingers in mediating multimerization, regardless of whether DNA plays a role.

### Hetero-multimers of divergent ZF arrays form less efficiently

Finally, to examine the specificity of ZF array binding, we generated HA- and V5-tagged constructs in which we replaced the final exon containing the human ZF array with a synthesized cDNA matching the final exon of the chimpanzee reference PRDM9 allele (W11a) and containing 18 zinc fingers, rather than 12. We refer to these as Chimp-HA and Chimp-V5 (illustrated in ***Figure 6***a). To test the relative efficiency of homo-versus hetero-multimerization in mixtures of Human and Chimp PRDM9, we performed direct competition experiments. We transfected cells with equimolar mixtures of DNA for three constructs, for example Chimp-V5 plus Chimp-HA plus Human-HA. In this case Chimp-V5 would be the “bait” pulled down by IP with anti-V5, and Chimp-HA and Human-HA would be the co-IP “prey” detected by western blotting with anti-HA (we replicated by reversing the tags). The results show that Chimp PRDM9 is >2-fold more efficiently pulled down, compared to Human PRDM9, by Chimp PRDM9. Conversely Human PRDM9 is >2-fold more efficiently pulled down than Chimp PRDM9, by Human PRDM9 (***Figure 6***c). Thus, PRDM9 preferentially forms homo-multimers rather than hetero-multimers, at least for ZF arrays as highly diverged as Human and Chimp.

## Discussion

Two striking properties of mammalian recombination that have been observed in multiple studies are that, although PRDM9 controls almost all hotspot positions (***Brick et al., 2012***), many apparent PRDM9 binding motifs are not in fact bound, while in mice at least, many PRDM9-bound sites do not become clear double-strand break hotspots (***Baker et al., 2014***). We identify factors responsible for these features, in part, and find that they differ even between humans and chimpanzees, in a manner dependent on the PRDM9 ZF-array.

The narrow widths and large number of our ChIP-seq peaks allowed us to recover no fewer than seven different modes of human PRDM9 binding with different internal spacings between several DNA-contacting zinc fingers (***Figure 1***), a pattern not detected in previous studies, and subtly distinguishing the PRDM9 “B” allele from the “A” allele. This revealed high binding specificity for many upstream fingers. Binding is strongly impacted by all zinc fingers—as for example directly seen in THE1B repeats—and involves extensive sequence specificity not captured by a single shared motif. This partially explains why using a single motif does not fully distinguish bound and unbound positions. Still, the strength of a match to our motif set correlates with but does not guarantee PRDM9 binding, and factors apart from the primary DNA sequence, including repeat context and local histone marks, influence human PRDM9 binding, with a preference for open chromatin regions including promoters, and H4K20me1 presence.

Compared to the human B allele, the chimp W11a allele shows different sequence preferences and its binding is associated with different epigenetic features (***Figure 2***), resulting in broad-scale binding differences between the human and chimp alleles (***Figure 4***). Other human ZF-genes show similar broad differences in binding ***preferences(Imbeault et al., 2017***), but it is interesting that this binding diversity is tolerated for recombination hotspot specification by PRDM9 (***Davies et al., 2016***).

Downstream of PRDM9 binding, hotspot presence/absence is subject to an additional level of regulation. At broad scales, recombination rates can be influenced by PRDM9-independent factors that increase the probability of DSB formation at PRDM9 binding sites across all levels of PRDM9 binding enrichment. For example, recombination rates increase at PRDM9 binding sites near telomeres in human male meiosis (***Pratto et al., 2014;Figure*** 1-S2). Here we show that, even outside of telomeric regions, broad-scale effects can influence recombination outcomes independently of PRDM9 binding and local sequence context (***Figure 3***-S2).

One strongly negative predictor of recombination outcomes is presence within an active gene promoter, marked by PRDM9-independent H3K4me3, an effect previously observed in mice (***Brick et al., 2012; Davies et al., 2016***). A recent study in mice (***Grey et al., 2017***) found that (like the chimp allele), two mouse PRDM9 alleles do not directly bind at promoters. When Spo11 was present to form DSBs, additional PRDM9 peaks appeared at a small number of promoters—hypothesized as due to indirect recruitment (***Grey et al., 2017***). In contrast, we observed human PRDM9 directly binding to many promoter regions, previously unobserved due to filtering of PRDM9-independent H3K4me3 peaks and the evident suppression of DSB formation at these sites (***Pratto et al., 2014***). Given the similarity of promoter composition and organization across cell types, the human A/B alleles likely bind to promoters *in vivo* as well. Thus, human PRDM9 might be unusual in this regard, and its properties imply that the direction of recombination away from promoters in humans does not simply owe to PRDM9’s binding preferences or creation of competitive H3K4me3 peaks, as has been suggested in mice with AT-rich PRDM9 binding motifs (***Brick et al., 2012***). Indeed if PRDM9’s binding preferences were responsible for keeping recombination away from promoters, one would expect PRDM9 alleles with promoter-enriched binding to be heavily selected against, but instead the nearly identical human A/B alleles have reached near-fixation in European populations.

Our analysis of thousands of hotspots centered atTHE1B repeats identified multiple sequence motifs, including the motif ATCCATG, which *in vivo* associates with >2-fold recombination suppression and acts downstream of PRDM9 binding. Therefore, DNA sequence outside PRDM9 binding motifs can strongly influence hotspot presence/absence in *cis*. Strikingly, these motifs do not impact PRDM9-dependent binding and resulting H3K4me3 deposition either in transfected cells (this study) or in testes (***Pratto et al., 2014***). They also map outside the PRDM9 motif region, while all motifs impacting binding fall within the motif region. Although diverse and spread throughout the center of these hotspots, these motifs instead share multiple features that overwhelmingly suggest a different, common causal mode of action, by impacting KRAB-ZNF protein binding. Several motifs overlap identified KRAB-ZNF binding target regions within THE1B and predict TRIM28 recruitment atTHE1B repeats; these motifs consistently associate with H3K9me3 deposition levels in a manner that linearly correlates with their impact on recombination. Interestingly, we also saw a weak increase in H3K4me3 signal whenever H3K9me3 increased, and this signal is also observed in testes, implying the motifs we find impact chromatin modifications in this tissue, and—unlike PRDM9—in many somatic cell types also. The motif ATCCATG consistently shows the most significant such associations and falls within a >50-bp motif for TRIM28 recruitment, likely by an unknown KRAB-ZNF protein, potentially with multiple ZNFs to specify this long target site. Indeed, examining all KRAB-ZNF proteins from a recent study (***Imbeault et al., 2017***) reveals a virtually universal pattern of local recombination suppression, particularly for those KRAB-ZNF proteins most strongly associated with H3K9me3 deposition at their binding sites. Thus, many of these KRAB-ZNF proteins are likely to exert functional influences during the early meiotic stages where recombination occurs.

Although in principle these effects on recombination might be due to KRAB-ZNF proteins protecting their underlying bound DNA from the meiotic DSB machinery, the recombination suppression impact of at least the motif ATCCATG operates even where PRDM9 does not bind the THE1B repeat in which it falls. Suppression occurs for >1 kb nearby, implying an ability to act at some distance and making direct physical action unlikely. Moreover, and interestingly, we observe no impact on recombination hotspots of the presence/absence of binding sites for other proteins such as DUX4 (***Young et al., 2013***), despite our observing clear impacts of DUX4 binding motif presence on local chromatin within THE1B repeats (***Figure 3***-source data 1). Instead, perhaps only certain chromatin modifications suppress recombination. At their binding sites, many KRAB-ZNF proteins recruit TRIM28 which in turn recruits histone remodeling proteins including SETDB1 and HP1, depositing the H3K9me3 modification, which has been associated with suppression of meiotic recombination in mice (***Buard et al., 2009; Walker et al., 2015***).

Promoters show strong PRDM9-independent H3K4me3 marks, while the recombination-suppressing motifs we identify are associated with PRDM9-independent H3K9me3, and weak H3K4me3, deposition. Interestingly, PRDM9 directly induces H3K4me3 at hotspot sites, and interacts with both readers and writers of H3K9me3 (***Parvanov et al., 2016***). It seems plausible that if placed independently of PRDM9 binding, these H3K4me3/H3K9me3 marks might disrupt co-ordination or timing of their placement, and hence in turn disrupt recombination. Indeed, our results (*e.g. **Figure 3***-S1a) show that pre-existing histone modifications correlate mainly negatively with recombination. The ideal nucleosome substrate for recombination hotspot formation might be devoid of specific histone modifications prior to PRDM9 binding, with PRDM9 able to produce and co-ordinate all required modifications.

Most KRAB-ZNF proteins bind repeats (***Imbeault et al., 2017***), and they constitute the largest family of transcription factors in mammals, with rapid evolution. Evidence suggests that the KRAB domain may have first evolved in an ancient ancestor of PRDM9 and then spread (***Birtle and Ponting, 2006***), so it is interesting that these partial descendants of PRDM9 appear to disrupt recombination. In general, KRAB-ZNF genes appear to emerge concomitantly with the spread of particular transposon families, and they play a role in repressing transposon activity (***Imbeault et al., 2017; Jacobs et al., 2014; Wolf et al., 2015; Rowe et al., 2013***). Paradoxically though, they often remain active long after their targets lose transpositional activity (***Imbeault et al., 2017***). Our results suggest that one possible reason might be an adaptive role for KRAB-ZNF genes in specifically suppressing meiotic recombination in and around repeats, which otherwise could be prone to mediating deleterious genomic rearrangements. If so, evolution of PRDM9 to bind new repeats might, in turn, lead on to co-evolution of ZNF genes, which contain KRAB domains that potentially evolved from PRDM9 itself.

Another consequence is that not only PRDM9 binding sites, but potentially many other sites within hotspots, are predicted to cause DSB initiation asymmetry, and thus to be influenced by biased transmission—as seen previously for PRDM9 motifs and GC-biased gene conversion in hotspots (***Boulton et al., 1997; Coop and Myers, 2007; Myers et al., 2010; Baker et al., 2015b; Smagulova et al., 2016; Davies et al., 2016***). Unlike self-destructive drive at PRDM9 motifs, such drive would bias the evolution of features with broad impacts across cell types, towards *increased* KRAB-ZNF binding and hence constitutive silencing of hotspot regions, even if this silencing is selectively disadvantageous.

The ability of PRDM9 to affect the transcription of bound promoters such as that of *CTCFL* may simply add another dimension to its pleiotropic effects across the genome, and this may even help to explain why a single PRDM9 allele predominates in humans. Speculatively, while a multitude of alleles may function equally well in specifying sites of meiotic recombination initiation, perhaps a subset can positively affect fertility by enhancing the expression of meiotic genes such as *CTCFL*, and these alleles are driven to high frequency by positive selection. A similar mechanism may also explain our finding that a predicted submotif shared by many chimp *PRDM9* alleles (***Schwartz et al., 2014***) corresponds to a group of chimp zinc fingers with a dominating influence on binding targets (***Figure 4***c).

Given DSB suppression at promoters, nearby PRDM9 binding sites might be immune from the effects of hotspot death, which would otherwise act to abolish its binding and drive potentially deleterious mutations—potentially including any which weaken the promoter—to fixation in these regions. Indeed, speculatively, this may even explain why recombination is actively suppressed at promoters in certain organisms.

We have also demonstrated that PRDM9’s ability to form multimers is mediated primarily by its zinc finger array, while two highly diverged human and chimp alleles form hetero-multimers less efficiently than homo-multimers. PRDM9’s zinc finger array has been regarded primarily as a DNA-binding domain with no other demonstrated functions, although studies of other zinc finger proteins have shown that ZF domains can participate in highly specific protein-protein interactions, including with each other (***McCarty et al., 2003; Lee et al., 2007***). The mammalian gene with the most similar ZF-array to PRDM9 is ZNF133, whose zinc fingers have an almost identical consensus sequence, apart from at DNA-contacting bases, to PRDM9. ZNF133 has been shown to interact with PIAS1 via its zinc fingers, and simultaneously bind its target sites (***Lee et al., 2007***). Thus, it seems credible that multimerization interactions involving PRDM9 might involve its zinc fingers. Interestingly, PIAS1 is recruited to DNA damage sites (***Galanty et al., 2009***). Currently, we can only speculate about what function PRDM9 multimerization might serve in meiosis. One intriguing hypothesis is that multimer formation may play some role in PRDM9-mediated homologue pairing, which we previously identified as a mechanism to explain the role of PRDM9 in fertility and speciation in mice (***Davies et al., 2016***). In this case, a preference for homo-multimer formation would have obvious advantages.

## Methods and Materials

### Cloning

A cDNA was custom synthesized to contain the full-length (2,685 bp) *PRDM9* transcript from the human reference genome (GRCh37), which is the B allele of *PRDM9*. 218 synonymous base changes were engineered into the exon containing the zinc finger domain in order to distinguish the synthetic copy of *PRDM9* from the endogenous copy and to facilitate proper synthesis of this highly repetitive region. We cloned this cDNA into the pLEXm transient expression vector (***Aricescu et al., 2006***) by ligation with a Venus (YFP) tag at its N-terminus, fused using an AgeI restriction site. A similar synthesized construct was designed to match exon 10 of the chimp PRDM9 reference allele (the “W11a” allele, 2022 bp, codon optimized for human expression and non-repetitiveness). Exons 1-9 were amplified from the human construct, and the chimp allele was fused at the N-terminus with an XbaI site. The ZFonly and noZF alleles were amplified using internal primers designed inside the full-length human construct. For the C-terminally tagged constructs, a 198-bp HA and 213-bp V5 linker were synthesized (having the sequence linker-TwinStrep-linker-HA/V5-linker-P2A) and cloned between each respective PRDM9 allele and a YFP tag using KpnI and AgeI sites, respectively. C-terminally tagged constructs were cloned into the pLENTI CMV/TO Puro DEST vector (Addgene plasmid # 17293; ***Campeau et al., 2009***), owing to its higher transient expression efficiency and to test the possibility of stable lentiviral transduction. Cloning into this vector was performed using the Gateway recombinase-based cloning system (Thermo Fisher Scientific). Constructs were cloned, amplified, and isolated using an Qiagen EndoFree Plasmid Giga Kit to yield transfection-quality DNA, which was verified by restriction digestion and Sanger sequencing.

### Transfection

HEK293T cells (ATCC CRL-3216) were chosen owing to their high transfection efficiency, rapid growth rate, and low-cost media requirements. Large-scale transfections of the N-terminal GFP-tagged Human PRDM9 construct were performed as described (***Aricescu et al., 2006***). Cells were grown in DMEM media (10% FCS, 1X NEAA, 2 mM L-Glut, Sigma D6546) in 200-ml roller bottles at 37°C/5% CO_2_. A transfection cocktail was prepared for each bottle by adding 0.5 mg of chloroform-purified construct DNA to 50 ml of serum-free DMEM (1X NEAA, 2 mM L-glut) and 1 mg polyethylenimine, followed by a 10-minute incubation, and then addition of 375 *μ*g of kifunensine. After the cells reached 75% confluence, the growth medium was removed from each roller bottle and replaced with 200 ml low-serum DMEM (2% FCS, 1X NEAA, 2 mM L-Glut) and 50 ml transfection cocktail. Cells were then incubated for 72 hours to enable expression of the transfected construct. Expression was verified by fluorescence microscopy.

We performed all subsequent smaller-scale transfections of the C-terminally tagged constructs in the pLENTI vector using the FuGENE-HD transfection reagent according to manufacturer instructions. HEK293T cells (ATCC CRL-3216) were thawed and incubated at 37°C with 5% CO_2_ in DMEM (Sigma D6546) supplemented with 10% fetal bovine serum (Sigma F7524), 1X L-Glutamine (Sigma G7513), and 1X penicillin/streptomycin (Sigma P0781). Confluent cells were split 1:10 and passaged for no longer than one month before transfection. The night before transfection, confluent cells were trypsinized (Sigma T3924), diluted in growth medium, and counted on an automatic hemocytometer (BioRad TC20). For each replicate, 15 million cells were seeded in 30 ml growth medium in a T175 cell culture flask. The following morning, cells were transfected by mixing 30 *μ*g total construct DNA into 800 *μ*l OPTI-MEM (Life Technologies 31985062), then carefully adding 90 *μ*l FuGENE-HD Transfection Reagent and flicking to mix, incubating at room temperature for 15 minutes, and then adding the mixture dropwise to each dish while swirling gently to mix. After 48 hours, cells were imaged briefly with a fluorescent microscope to confirm expression, and were subsequently harvested. As negative controls, additional cells were seeded at the same time but were not transfected.

### ChIP (N-terminal YFP-Human)

ChIP-seq was performed according to an online protocol produced by Rick Myers’s laboratory ***Johnson et al., 2007***), which was used to produce much of the ENCODE Project’s ChIP-seq data (***ENCODE, 2012***), with several optimizing modifications.

#### Crosslinking

Bottles were removed from the incubator and shaken vigorously to detach cells. Fresh formaldehyde was added to a final concentration of 0.75% and cells were incubated at room temperature for 15 minutes. The crosslinking reaction was stopped by adding glycine to a final concentration of 125 mM. Cells were aliquoted to 50 ml conical tubes, centrifuged (2000g, 5 minutes), resuspended in cold 1X PBS, and centrifuged again. Pellets were snap frozen with dry ice, and then stored at −80°C.

#### Lysis and Sonication

Frozen pellets were thawed and resuspended in cold Farnham Lysis Buffer (5 mM PIPES pH 8.0, 85 mM KCl, 0.5% NP-40,1 tablet Roche Complete protease inhibitor per 50 ml) to a concentration of 20 million cells per ml, then passed through a 22G needle 20 times to further lyse and homogenize them. Technical replicates were processed in parallel from this point forward (with only one replicate performed for transfected H3K4me3). Lysates were centrifuged and resuspended in 300 *μ*l cold RIPA lysis buffer (1X PBS, 1% NP-40, 0.5% sodium deoxycholate, 0.1% SDS, 1 tablet Roche Complete protease inhibitor per 50 ml) per 20 million cells to lyse nuclei. 300 *μ*l samples were sonicated in a Bioruptor Twin sonication bath in 1.5 ml Eppendorf tubes at4°C for two 10-minute periods of 30 seconds on, 30 seconds off at high power. Cell debris was removed by centrifugation (14,000 rpm, 15 minutes, 4°C), and supernatants were isolated and brought to a final volume of 1 ml with RIPA. These chromatin preps were snap-frozen in dry ice then stored at −80°C.

#### Immunoprecipitation

Magnetic beads were washed by adding 200 *μ*l Invitrogen Sheep AntiRabbit Dynabeads per sample to 800 *μ*l cold PBS/BSA(1X PBS, 5 mg/ml BSA, 1 tablet Roche Complete protease inhibitor per 50 ml, filtered with 0.45 micron filter). Solutions were placed on a magnetic rack and resuspended in 1 ml PBS/BSA four times. 5 *μ*l Abcam rabbit polyclonal ChIP-grade anti-GFP antibody (ab290) or rabbit polyclonal ChIP-grade anti-H3K4me3 antibody (ab8580) was added and solutions were incubated overnight at 4°C on a rotator. Antibody-coupled beads were washed three times with cold PBS/BSA and resuspended in 100 *μ*l PBS/BSA, then added to 1 ml chromatin preps thawed on ice. One tube was prepared in parallel without adding beads, to yield a genomic background control sample from total chromatin. Tubes were incubated for 12 hours on a rotator at 4°C, then washed 5 times for 3 minutes each with cold LiCl Wash Buffer (100 mM Tris pH 7.5, 500 mM LiCl, 1% NP-40,1% sodium deoxycholate, filtered with a 0.45 micron filter unit), then washed once with cold 1XTE (10 mM Tris-HCl pH 7.5, 0.1 mM Na_2_-EDTA). Bead pellets were resuspended in 200 *μ*l room-temperature IP elution buffer (1% SDS, 0.1 M NaHCO_3_, filtered with a 0.45 micron filter unit) and vortexed to mix.

#### Reverse crosslinking and DNA purification

Samples were incubated in a 65°C water bath for 1 hour with mixing at 15-minute intervals to uncouple beads from protein-DNA complexes. Samples were centrifuged (14,000 rpm, 3 mins) and placed on a magnet to pellet beads, and supernatants were isolated and then incubated in a 65°C water bath overnight to reverse crosslinks. DNA was purified using a Qiagen MinElute reaction cleanup kit and quantified using a Qubit High Sensitivity DNA kit.

### ChIP (C-terminal-tagged constructs)

Slight modifications were made for the smaller-scale transfection experiments with C-terminally tagged constructs. Crosslinking was performed in 1% formaldehyde for 5 minutes. Input chromatin was “pre-cleared” to remove chromatin bound non-specifically by the beads. For each sample, 50 *μ*l of equilibrated magnetic beads were resuspended in 100 *μ*l PBS/BSA and added to the chromatin samples for pre-clearing for two hours at 4°C with rotation. Beads were removed, and 100 *μ*l of pre-cleared chromatin was set aside for the input control. 5 *μ*l ChIP-grade rabbit polyclonal antibody (Abcam anti-HA ab9110, anti-V5 ab9116, anti-H3K4me3 ab8580, or anti-H3K36me3 ab9050) was added to the remaining pre-cleared chromatin and incubated overnight at 4°C with rotation. 50 *μ*l beads were washed and resuspended as before, then incubated with the chromatin samples for two hours at 4°C with rotation. After washing and decrosslinking, samples were further incubated with 80 *μ*g RNAse A at 37°C for 60 minutes and then with 80 *μ*g Proteinase K at 55°C for 90 minutes.

### ChIP sequencing, mapping, and filtering

DNA was submitted to the Oxford Genomics Centre for library preparation, sequencing, and mapping. For the N-terminal YFP-Human experiments, ChIP and input chromatin DNA samples from transfected and untransfected cells were sequenced in multiplexed paired-end Illumina HiSeq1000 libraries, yielding 51 -bp reads. Samples from transfected cells were multiplexed across 3 lanes, yielding roughly 77-101 million properly mapped read pairs (*i.e*. fragments) per replicate. Samples from untransfected cells (processed independently) were multiplexed across 2 lanes, yielding roughly 60-99 million properly mapped fragments per sample. For the C-terminal tag experiments, ChIP and input chromatin DNA samples from transfected and untransfected cells were sequenced all together in 6 lanes of paired-end Illumina HiSeq2500 libraries (rapid mode), yielding 51-bp reads with 37 to 64 million reads per replicate. Coverage was chosen in each experiment to exceed recommendations for doing ChIP-seq with sufficient power to detect the majority of true binding events (***Landt et al., 2012***).

Sequencing reads were aligned to hg19 using BWA (v0.7.0-r313, option -q 10, ***Li and Durbin, 2009***) followed by Stampy (v1.0.23-r2059, option -bamkeepgoodreads, ***Lunter and Goodson, 2011***), and reads not mapped in a proper pair or with an insert size larger than 10 kb were removed. Read pairs representing likely PCR duplicates were also removed by samtools rmdup (v0.1.19-44428cd, ***Li et al., 2009***). Pairs for which neither read had a mapping quality score greater than 0 were removed. For samples with only one replicate, fragments were split at random into two equally-sized pseudo replicates. Fragment coverage from each replicate was then computed at each position in the genome using in-house code and the samtools (v0.1.19-44428cd) and bedtools (v2.23.0, genomecov -d) packages (***Li et al., 2009; Quinlan and Hall, 2010***). Details of the ChIP-seq samples are listed below.

|  | Transfection with: | ChIP antibody | Proportion signal, rep1 | Proportion signal, rep2 | Fragment number, rep1 | Fragment number, rep2 | Peak number | Fraction of genome enriched |
| --- | --- | --- | --- | --- | --- | --- | --- | --- |
| N-terminal tag | YFP-hPRDM9 | GFP | 0.225 | 0.374 | 80,844,035 | 79,526,431 | 170,198 | 0.012 |
|  | YFP-hPRDM9 | H3K4me3 | 0.750 | n/a | 77,625,060 | n/a | 470,314 | 0.052 |
|  | YFP-hPRDM9 | none (Input) | n/a | n/a | 100,861,414 | n/a |  |  |
|  | Untransfected | H3K4me3 | 0.823 | 0.794 | 59,156,993 | 72,839,266 | 45,758 | 0.007 |
|  | Untransfected | none (Input) | n/a | n/a | 98,664,592 | n/a |  |  |
| C-terminal tags | cPRDM9-HAorV5 | HA/V5 | 0.443 | 0.394 | 36,662,728 | 44,594,666 | 247,717 | 0.011 |
|  | hPRDM9-HAorV5 | HA/V5 | 0.374 | 0.510 | 39,385,214 | 38,717,735 | 213,885 | 0.020 |
|  | hPRDM9-HA | H3K4me3 | 0.522 | 0.544 | 52,439,279 | 54,451,480 | 221,446 | 0.024 |
|  | hPRDM9-HA | H3K36me3 | 0.334 | n/a | 59,690,192 | n/a | 33,625 | 0.001 |
|  | hPRDM9-HA | none (Input) | n/a | n/a | 53,219,513 | n/a |  |  |
|  | Untransfected | H3K4me3 | 0.680 | 0.669 | 57,205,316 | 60,883,503 | 37,932 | 0.006 |
|  | Untransfected | H3K36me3 | 0.502 | n/a | 52,368,417 | n/a | 263,983 | 0.027 |
|  | Untransfected | none (Input) | n/a | n/a | 56,445,392 | n/a |  |  |

**ChIP-seq datasets generated in this study.** The datasets utilized in this analysis include the N-terminal YFP-tagged human construct used for most of the analysis as well as the C-terminal tagged constructs used in subsequent experiments. Columns 3 and 4 list the proportion of fragments estimated to arise from true signal genome-wide, as computed by our peak calling algorithm. Replicate 2 is assigned “n/a” when only one replicate was performed. Total peak numbers on the autosomes and on the X chromosome are listed in the second-to-last column (HEK293T cells lack a Y chromosome). The final column is an estimate of the proportion of 100-bp bins in the genome with evidence of enrichment at $p < 10^{-5}$ .

We compared the C-terminal Human-HA/V5 data with the N-terminal YFP-Human data and found strong overlap between the peak sets (60%) but a poor correlation in raw coverage values or in our computed enrichment values (r = 0.3). We explored this further and noticed that the newer sequencing run had a strong increase in coverage of GC-rich regions (nearly two-fold higher input coverage in regions with >60% GC), perhaps owing to differences in the ChIP protocol or to downstream differences in the library prep and sequencing steps (Illumina HiSeq 1000 versus Illumina HiSeq 2500). We also cannot exclude any effects due to the different placement of the tags. Due to this strong GC bias, we utilized the N-terminal YFP-Human dataset exclusively for most analyses of the human allele, except when directly comparing to data obtained using the C-terminal Human-HA/V5 constructs (ATAC-seq, RNA-seq, H3K36me3 ChIP-seq, Chimp ChIP-seq).

### Calling PRDM9 binding peaks

We developed a maximum-likelihood-based peak calling algorithm that takes as input the number of fragments overlapping a bin (a single base position or an interval) from two ChIP replicates and a genomic background control, as well as three constants describing the coverage ratios between these three inputs, which are estimated genome-wide in an initialization step. The Poisson distribution was chosen as a model of sequencing coverage given its support on all non-negative integers and simple parameterization. As specified, this model assumes that the coverage due to signal is proportional between the two ChIP-seq replicates across the genome and that the coverage due to background is proportional among all 3 lanes across the genome. We allow for local estimates of background and signal to account for sequence coverage biases and mappability differences across the genome. *Ab initio* single-base peak calling proceeds in three stages: 1) estimation of constants given coverage values in 100-bp non-overlapping bins genome-wide, 2) single-base maximum likelihood estimation given constants and single-base coverage values, 3) calling of peak centers in the likelihood landscape given a p-value threshold and a threshold on the minimum separation between peak centers.

#### Definitions

Let *D*_1_(*i*), *D*_2_(*i*) and *G*(*i*) be random variables representing the fragment coverage in bin *i* from the two ChIP-seq replicates and the genomic control, respectively (and let *d*_1_(*i*), *d*_2_(*i*) and *g*(*i*) represent the observed coverage in bin *i*). We model the coverage of each sequencing replicate *j* at bin *i* as a sample from a Poisson distribution with mean *λ_j_*(*i*),

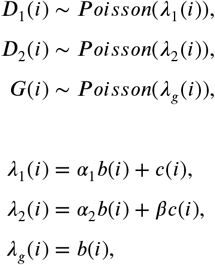

where *α*_1_ and *α*_2_ are constants defining how coverage due to background in the ChIP replicates compares to *b*(*i*), a parameter representing the mean coverage in the genomic control lane at bin *i*; and *β* is a constant defining how coverage due to binding enrichment in ChIP replicate 2 compares to *c*(*i*), a parameter representing the coverage due to binding enrichment in ChIP replicate 1 at bin *i*. We wish to test the hypothesis that *c*(*i*) ≥ 0 for each bin *i*.

#### Estimating constants

To speed up this step and to provide smoother coverage estimates, we first computed coverage values in 100-bp bins across the autosomes. One can estimate *α_j_* by assuming (conservatively) that when *d*_1_(*i*) = 0 or *d*_2_(*i*) = 0, *c*(*i*) = 0. That is, one can assume that if ChIP replicate *j* has coverage 0 at bin *i*, then any coverage in the other replicate (*j*’) arises purely from background. Thus for all *i* such that *d_j_*(*i*) = 0

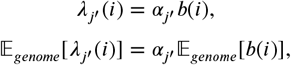

and thus one can estimate *α_j_*, as

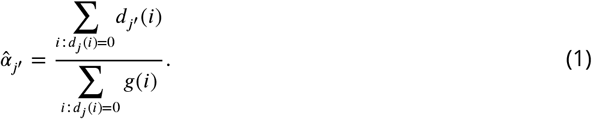

Now an initial estimate of *β* can be computed using genome-wide coverage means 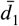, 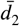, 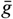 as follows:

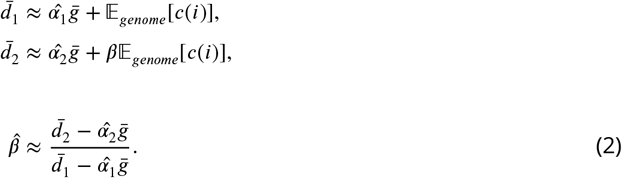

Next, maximum likelihood estimation and hypothesis testing are performed across all bins (see below), and 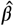 is re-computed as above, using coverage means from the subset of bins with p< 10^-10^, for which the ratio of coverage between the two replicates will be less affected by noise.

Finally, using the MLEs 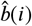 and 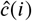 for each bin (see subsection below), a genome-wide estimate of the proportion of reads from signal is computed as

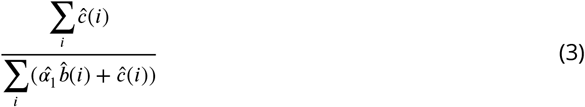

for replicate 1 and as

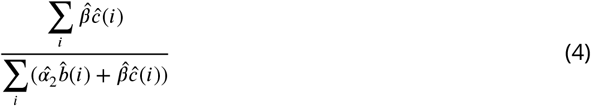

for replicate 2.

#### Hypothesis Testing

With these estimates of *α_j_* and *β*, one can compute Maximum Likelihood Estimators for the unknown parameters *b*(*i*) and *c*(*i*) at each bin *i* from the coverage data *d*_1_(*i*), *d*_2_(*i*) and *g*(*i*) (see below for derivation). Then, using these MLEs one can compute a log-likelihood ratio test statistic against a null model in which *c*(*i*) = 0:

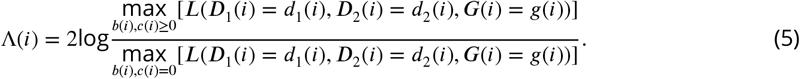

Under the null hypothesis, the test statistic Λ(*i*) is distributed approximately as a *χ*^2^ distribution (with 1 degree of freedom due to the parameter *c*(*i*) and an atom of probability at 0), yielding a p-value at each bin *i* indicating the probability that the observed likelihood ratio could arise from background alone.

#### Calculation of Maximum Likelihood Estimators

Recall that at each position the Poisson means for coverage in each lane are (dropping the *i* notation for succinctness)

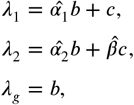

where 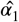, 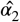, and 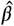 are constants estimated for the whole genome. To simplify calculations, we reparameterize using a new variable *y* = *c*/*b* and rewrite the above equations as

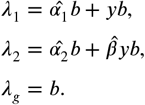

Given the observed coverage values *d*_1_, *d*_2_, and *g*, the Poisson log likelihood function can be written as

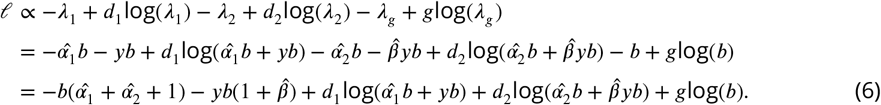

Now to maximize *ℓ* we first obtain the partial derivatives for *b* and *y*

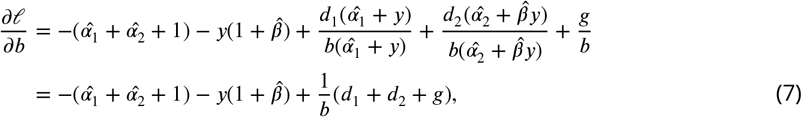

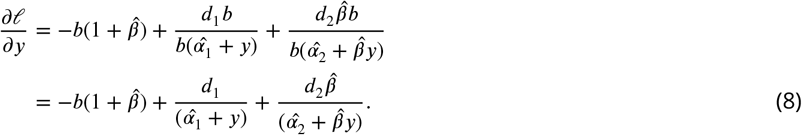

Next, we set the partials to 0 and solve them as a system to obtain any potential local maxima. We start by solving for *b* in *Equation 7* as follows:

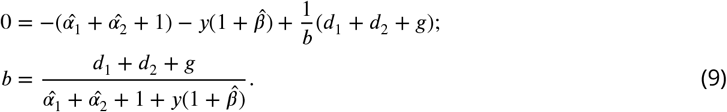

Then, we substitute it into *Equation 8* and rewrite it as follows, with the aim of simplifying it into quadratic form:

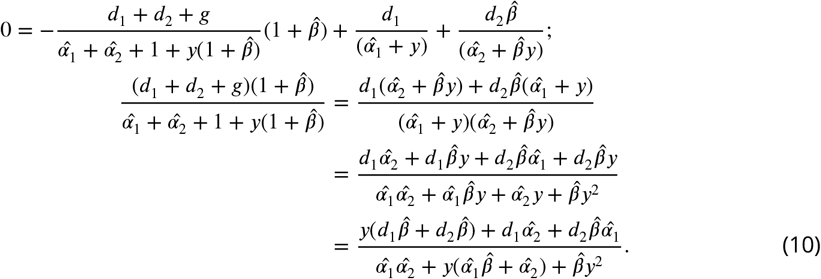

To shorten notation, we substitute in the following variables for constant terms in *Equation 10:*

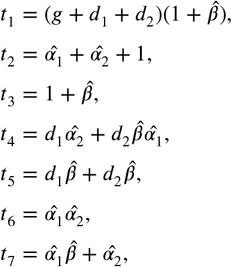

yielding

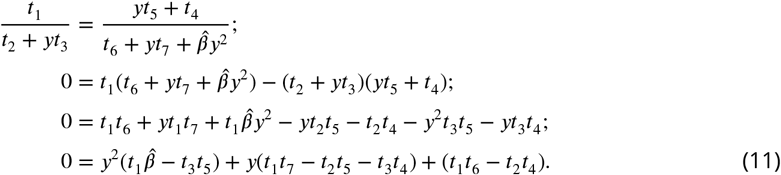

Now we can solve for *y* in ***Equation 11*** using the quadratic formula, taking the positive root to be 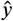, the MLE for *y*, which we report as the “enrichment” value for that bin. To obtain 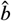, we simply substitute 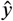 into ***Equation 9*** and, to return to the original paramaterization, 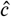 is simply computed as 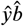. Finally, to obtain 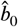, the MLE for *b* under the background model, we can simply set *y* to 0 in ***Equation 9***, yielding

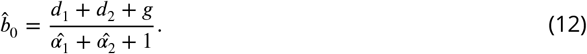

#### Peak calling and centering

Given a likelihood ratio value Λ(*i*) for each base *i* along a chromosome, along with a p-value threshold (which is converted to a lower bound on the likelihood ratio, *l*) and *m*, a threshold on the minimum separation between peak centers, initial peak centers are found by identifying all significant bases (bases for which Λ(*i*) > *l*) that are local maxima. Specifically, each significant base is scanned to test if

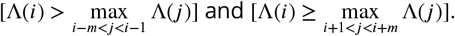

At each initial peak center satisfying these criteria, a confidence interval is computed by identifying the nearest position *j* to the left and to the right (by a maximum of 1000 bp) where (Λ(*i*)–Λ(*j*)) > 9.12, which defines a 99% confidence interval for the peak center (using 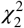, with one degree of freedom for the enrichment factor and one for the peak center position). All confidence intervals along a chromosome are then sorted from narrowest to widest, and in this order each confidence interval is added one at a time to the final peak set, provided it does not overlap any of the confidence intervals already included in the final peak set. This produces a final peak set with non-overlapping confidence intervals, favoring inclusion of stronger peaks with narrower confidence intervals. Finally, to refine peak centers in confidence intervals with multiple tied bases, the rounded mean position of all maximal bases is reported as the peak center. The resulting final peak set reports 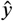 and the p-value for Λ at the peak center as the enrichment and p-value for that peak.

#### Force-calling

This algorithm enables maximum likelihood estimation and hypothesis testing at any arbitrary bin in the genome, when provided with coverage values and estimates of *α*_1_, *α*_2_, and *β*. This enables us to “force-call” enrichment and p-values at pre-specified locations in the genome, for example to determine what fraction of gene promoters show evidence of H3K4me3 enrichment in a 1-kb window centered on the transcription start site.

#### Overlap correction

When comparing peak sets to determine overlap proportions, one must account for chance overlaps owing to the width and number of peaks being compared. For comparisons between single-base peak centers and DSB hotspot intervals, for example, we computed the expected number of chance overlaps *c* between the *n* peak centers and the *t* hotspot intervals, each with width *w_i_*, in a genome of size *g* as

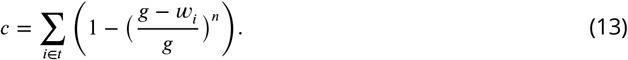

For more complicated comparisons, for example between two sets of intervals, we computed chance overlaps by randomly shifting the positions of one set of intervals uniformly in the interval [-60000, 60000], then counted the resulting overlaps to estimate *c*.

Given *f* observed overlaps between the sets of *n* and *t* peaks, we can compute the corrected overlap fraction, *o*/*t* as follows. Let *o*/*t* be the proportion of systematic overlaps, *c*/*t* be the fraction of chance overlaps, and *f*/*t* be the proportion of total overlaps. The probability of no overlap is simply the product of the complements of chance and systematic overlaps, as follows:

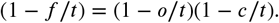

Solving for *o*/*t* then yields:

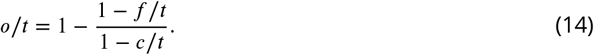

#### Motif finding

For each peak, a 300-bp sequence (centered on the called peak center) was extracted from the reference sequence (hg19). *Ab initio* motif calling was performed on sequences from the top 5,000 peaks (ranked by enrichment) that passed a set of stringent filters (p<10^-10^, enrichment >2, C.I. width ≤50, no bases overlapping annotated repeats, number of input reads between 10%ile and 90%ile, and ≥30 reads from ChIP rep1 + ChIP rep2). Motif calling proceeded in two stages: seeding motif identification, and joint motif refinement. Each seeding motif was obtained by first counting all 10-mers present in all input sequences, and from the top 50 most frequently occurring 10-mers, the one with the greatest over-representation in the central 100 bp of each peak sequence was chosen. This seeding 10-mer was then refined for 100 iterations as described in ***Davies et al.*** (***2016***), and all peak sequences containing matches to this refined motif were removed. From the remaining sequences, a new 10-mer was found and refined into a seeding motif, and this process was iterated up to 20 times. The 20 resulting seeding motifs were then refined jointly for 200 iterations as described (***Davies et al., 2016***). Three separate runs were performed for each sample to verify consensus. For the YFP-Human peaks, a run producing 17 final motifs was chosen, and of these the 7 motifs with ≥85% of matches occurring in the central 100 bp of each peak sequence were chosen as the final set in order to remove degenerate motifs (*i.e*. those with little base specificity at any position) as well as likely false positives (such as a match to the motif for the AP1 transcription factor). For the Chimp-HA/V5 peaks, only two motifs were produced, one of which was a degenerate CT-rich motif found in only 10% of peaks (but not centrally enriched), so it was filtered out (not shown). These final motifs were then force-called on the full set of peaks (without any peak filtering) by rerunning the refinement algorithm (***Davies et al., 2016***) with the option to not update the motifs with each iteration. The motif with the greatest posterior probability (of at least 0.75) of a match was reported for each peak, along with position and strand. For identifying motif matches genome wide, we used FIMO (version 4.10.0; ***Bailey et al., 2015***).

### GLM classifier of binding in 100-bp bins

To better understand the factors influencing PRDM9 binding at fine scales when expressed in HEK293T cells, we first split the autosomes (hg19) into non-overlapping 100-bp windows, then counted PRDM9 ChIP and Input fragments overlapping each bin and performed likelihood ratio testing as described (***Hinch et al., 2014***) to assign an Enrichment and P-value to each bin. We then determined whether each window overlaps peak regions from various histone mark ChIP-seq experiments carried out by the ENCODE project (***ENCODE, 2012***) in K562 cells: H2AZ, H3K27ac, H3K27me, H3K36me3, H3K4me1, H3K4me3, H3K79me3, H3K9AC, H3K9me1, H3K9me3, H4K20me1. Similarly, an indicator variable was created for DNase Hypersensitive Sites, as measured by the ENCODE project across many cell types (***ENCODE, 2012***). Indicator variables were also created to indicate whether a given bin overlaps an annotated repetitive sequence, and if so whether it overlaps a repeat of the L1, L2, Alu, orTHE1 classes. The proportion of GC bases within each window was also reported, along with the maximum PRDM9 motif score within each bin, as computed by FIMO software (***Bailey et al., 2015***). Bins were filtered to exclude those with fewer than 5 or greater than 50 overlapping Input fragments (removing the bottom 10% and top 0.1% of coverage to eliminate outlying repetitive regions or regions with poor coverage). Peaks were defined as bins with p<10^-5^ and enrichment >2 (~100k bins for human), and non-peaks were defined as bins with p>0.5 and 0 enrichment (~9.3M bins for human).

To set up a binary classification problem that could be easily modeled and interpreted, nonpeaks were subsampled to an identical number as the peaks dataset and merged to serve as the input for modeling. This dataset was randomly split into five subsets, and the fifth subset was reserved as a held-out test set for the final model. Iterative forward selection was carried out on the first three subsets, with an objective function equal to the model’s predictive accuracy on the fourth subset. That is, variables were incorporated into the model one at a time, choosing the variable that yielded the greatest increase in predictive accuracy on the held-out fourth subset at each step. The entire process, from subsampling non-peaks and test/training subsets to building a model by forward selection, was repeated ten times, and the relative order of incorporation of each explanatory variable was recorded in each replicate to ascertain model stability. The model used was a generalized linear model of a binomial family with a logit link function, as the dependent variable (peak/non-peak status) is binary. A predicted logit value of 0.5 was chosen as a threshold for classification, and classification error was determined by counting mismatches between predicted and observed classifications.

### ATAC-seq

ATAC libraries were prepared as described (***Buenrostro et al., 2013***). Briefly, 50,000 cells were lysed in 10 mM Tris-HCl pH 7.4, 10 mM NaCl, 3 mM MgCl_2_, 0.1% IGEPAL CA-630 and the nuclei were pelleted at 500*g* for 10 minutes. The transposition reaction was carried out for 30 minutes at 37°C using the Nextera DNA Sample Preparation Kit (Illumina) according to the manufacturer’s instructions. The libraries were purified using the MinElute PCR Purification Kit (Qiagen), PCR amplified, multiplexed, and sequenced by the Oxford Genomics Centre on an Illumina HiSeq2500 (rapid mode) to produce 60-77 million sequenced fragments (51-bp, paired-end reads) per sample. Reads were mapped to the hs37d5 reference (***Consortium, 2012***) using BWA (v0.7.0-r313, ***Li and Durbin, 2009***) followed by Stampy (v1.0.23-r2059, with option -bamkeepgoodreads, ***Lunter and Goodson, 2011***). PCR duplicates, mtDNA-mapped reads, reads not mapped in a proper pair, reads with mapping quality equal to 0, and pairs with an insert size larger than 2 kb were removed using samtools (v0.1.19-44428cd, ***Li et al., 2009***), leaving ~11 million fragments per sample. Using inhouse code, fragments were split by size into inter-nucleosome (51-100 bp) and mono-nucleosome fragments (180-247 bp), and the position of the central base in each fragment was reported, as described (***Buenrostro et al., 2013***). This yielded ~1 million inter-nucleosome and ~3 million mono-nucleosome fragments per sample. Fragment center coverage was computed genome-wide using bedtools (***Quinlan and Hall, 2010***).

### RNA extraction and RT-qPCR

Total RNA was extracted using the RNeasy kit (Qiagen) from three biological replicates (independently transfected in separate wells in parallel) per sample. For quantitative PCR analysis, RNA was reverse-transcribed using Expand Reverse Transcriptase (Roche), according to the manufacturer’s instructions. qPCR reactions were carried out in duplicate for each sample using Fast SYBR Green Master Mix (Applied Biosystems) on a CFX real-time C1000 thermal cycler (Bio-Rad), following the manufacturer’s guidelines. Data were analyzed using the CFX 2.1 Manager software (Bio-Rad) and normalized to the Tata binding protein (***TBP***) gene. Relative gene expression levels were calculated using the ΔΔC_*t*_ method, after averaging the two technical replicates for each sample. Statistical analysis was carried using a one-tailed t test. Primer sequences are given below.

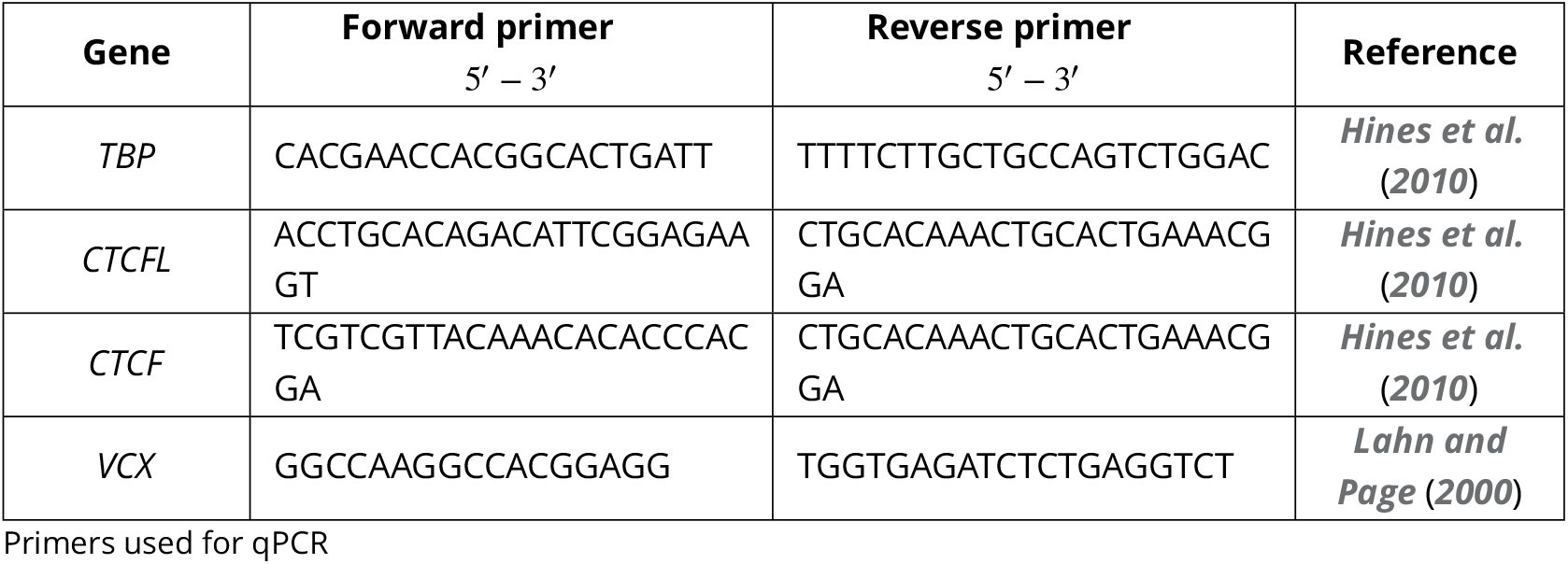

### RNA-seq

Total RNA was submitted to the Oxford Genomics Centre for mRNA enrichment, library preparation, and sequencing. Samples were multiplexed and sequenced on an Illumina Hi-Seq2500 (rapid mode), yielding 71-98 million 51-bp read pairs per sample. We created a custom reference sequence by merging the hs37d5 reference (used by the 1000 Genomes Project to improve mapping quality (***Consortium, 2012)***) with the construct and vector sequences transfected into our cells. Data were analyzed using the Tuxedo software package (***Trapnell et al., 2012***). Reads were mapped and processed using TopHat (version 2.0.13, options -mate-inner-dist=250 -mate-std-dev 80 – transcriptome-index=Ensembl.GRCh37.genes.gtf); followed by Cufflinks, CuffQuant, and CuffDiff (version 2.2.1); then analyzed using CummeRbund.

We searched for all genes with evidence of H3K4me3 within 500 bp of a TSS in the human-transfected sample (p<0.05, force-calling, requiring >5 input reads) and with defined FPKM values in the untransfected sample. Of the 14,667 genes passing these filters, 10,652 (73%) have a human PRDM9 binding peak within 500 bp of the TSS. Of these, 873 showed at least some evidence of differential expression between the human-transfected and untransfected samples (p<0.05), and of these 76 are significant after correction for multiple testing, with 46 significant only in the human-transfected sample (p<0.05 after Benjamini-Hochberg correction).

### Cell culture and transfection for co-IP experiments

For each experiment, 10 million cells were seeded in 20 ml growth medium in a 15-cm round cell culture dish. The following morning, cells were transfected by mixing 30 *μ*g total DNA into 800 *μ*l OPTI-MEM (Life Technologies 31985062), then carefully adding 90 *μ*l FuGENE-HD Transfection Reagent and flicking to mix, incubating at room temperature for 15 minutes, and then adding the mixture dropwise to each dish while swirling gently to mix. After 48 hours, cells were imaged briefly with a fluorescence microscope to confirm expression and were subsequently harvested. As negative controls, additional cells were seeded at the same time but were not transfected.

### Cell lysis and immunoprecipitation for co-IP experiments

Dishes were aspirated to remove media and cells were washed with cold PBS. 2 ml of cold lysis buffer (1% Triton X-100,150 mM NaCl, 50 mM Tris pH 8.0 plus 2X final concentration of Roche cOmplete Protease Inhibitor Cocktail Tablets) were added and cells were collected into 2 ml Eppendorf tubes using a cell scraper. Tubes were incubated on ice for 30 minutes and lysates were dounced 20 times in a 2 ml dounce homogenizer with a tight pestle to help shear nuclear membranes. Cells were spun at 2000g for 5 minutes to remove chromatin and cell debris. 100 *μ*l of lysate was set aside as an input control, and the remainder was split evenly among experimental and mock IP conditions. 2 *μ*g of primary antibody (Abcam ChIP-grade rabbit polyclonal anti-HA ab9110 or anti-V5 ab9116, or rabbit polyclonal IgG isotype control ab171870) was added and lysates were incubated for 1 hour at 4°C with rotation. For each sample, 25 *μ*l of magnetic beads (Invitrogen M-280 Sheep Anti-Rabbit Dynabeads) was equilibrated by washing 3 times in 1 ml cold PBS/BSA (1X PBS, 5 mg/ml BSA, filtered with 0.45-micron filter), then resuspending in 25 *μ*l PBS/BSA. Beads were added to the lysates and incubated for an additional hour at 4°C. Tubes were spun down and placed on a magnetic rack for 1 minute. Beads were pipetted up and down in 1 ml cold lysis buffer and rotated for 3 minutes at 4°C. Washing steps were repeated 4 more times, with all steps taking place in a cold room at 4°C.

### Western Blotting

Beads were resuspended in 20 *μ*l 2X Laemmli western loading buffer and boiled for 5 minutes at 100°C. Beads were removed on a magnetic stand and supernatants were diluted two-fold. The total protein concentrations of input lysates were estimated using a Pierce BCA Protein Assay Kit (Life Technologies 23227) and a NanoDrop spectrophotometer. 4X Laemmli buffer was added to 50 *μ*g of input protein to a final concentration of 1Xthen boiled for 5 minutes at 100°C. Samples were run on 10-well 7.5% BioRad mini-Protean TGX pre-cast gels at 150 Volts in standard TGX running buffer for approximately 1 hour, using 5 *μ*l of Full-Range Rainbow Ladder (VWR 95040-114) in one well. Gels were then assembled onto a BioRad mini Trans-Blot transfer pack (with PVDF membrane) according to manufacturer instructions and run on a Trans-Blot Turbo machine on the Mixed MW setting (2.5A, up to 25V, 7 mins). Membranes were quickly removed and transferred to 50 ml conical tubes, then blocked for 5 minutes with rotation in 10 ml Blocking Buffer (5% milk in PBS with 0.1% Tween-20), which was then poured off. Primary antibodies were diluted 1:5000 in 5 ml blocking buffer and added to the membranes and incubated for 1 hour at room temperature with rotation. Membranes were washed 3 times for 5 minutes each in PBST (PBS with 0.1% Tween). Secondary antibody (Amersham ECL Donkey anti-Rabbit IgG, HRP-linked, NA934) was diluted 1:30,000 in blocking buffer, then 5 ml was added to each membrane and they were incubated for 1 hour at room temperature with rotation. Membranes were washed an additional 3 times in PBST and one final time in PBS. Blots were imaged using a BioRad Clarity ECL kit according to manufacturer instructions and placed between sheets of transparency film to prevent drying during imaging. Imaging was performed using a BioRad ChemiDoc MP Instrument using chemiluminescence hi-sensitivity settings and signal accumulation mode for various exposure times. Image processing was performed in the BioRad ImageLab software, in which relative bands intensities were quantified by densitometry.

### Immunofluorescence detection of PRDM9 protein variants

HEK293T cells were seeded onto glass coverslips pre-treated with Poly-L-Lysine (SIGMA). Transfections with FL, ZF only and no ZF V5-tagged PRDM9 constructs were carried out for 24h, as described above. Cells were fixed for 20 min in chilled methanol, washed 3 times in PBS, permeabilized for 10min in PBS containing 0.1% Triton X-100, washed again, and blocked for 1h at RT in PBS supplemented with 0.1% Tween 20 and 1% BSA. Cells were immunostained with an anti-V5 antibody (Abcam ab9116) overnight at 4°C, washed, and incubated for 1h at RT with an appropriate secondary antibody conjugated to the Alexa Fluor 594 dye (Thermo Fisher Scientific). Coverslips were mounted in medium containing DAPI (Vectashield) and the cells were observed on a Olympus BX60 microscope for epifluorescence equipped with a Sensys CCD camera (Photometrics). Images were captured using the Genus Cytovision software (Leica Microsystems).

### Details of THE1B analysis

We developed an approach to identify motifs associating with various cellular phenotypes generated by or studied in this paper, specifically in and around THE1B elements. THE1B repeats are homologous repeat elements found across the genome, are non-genic in general, and are centers of hotspot activity. We sought to characterize how (and if) naturally arising DNA sequence differences across the 20696 autosomal THE1B copies impact both recombination and other measurable epigenetic features of them. Robustly identified associations are likely to be causal (*i.e*. identify DNA features influencing traits of interest), because the underlying DNA sequences are not in general believed to be specifically and consistently altered by the presence/absence of epigenetic features but, instead, can influence these features. Our approach used association testing to identify possible associations, and leveraged conditional testing to successively identify independent signals. This accounts for the fact that overlapping motifs, and even non-overlapping motifs, are correlated in which THE1B elements possess them. We performed testing based on the exact occurrence of 7-bp motifs. This length was chosen as a balance between specificity within the THE1B sequence, and occurring relatively commonly across THE1B elements. First, for the 20696 autosomal THE1B LTR elements annotated by the RepeatMasker software (hg19/Build 37, downloaded from the UCSC genome browser, and mapped to the positive strand relative to the THE1B consensus sequence) we produced a 20,696×16,384 matrix recording presence/absence of each motif of length 7 in each THE1B copy, across the genome. All subsequent analyses were then restricted to the 2021 such motifs present in at least 500 different THE1B elements (*i.e*. at least 2.5% of THE1B copies, aiding statistical power to detect potential associations). For each matrix row, we can view the set of motifs present as characterizing a single THE1B repeat copy in terms of common “variation” across such THE1B repeat copies. We annotated each THE1B repeat copy with various “phenotypes” - for example whether a recombination hotspot was present at that repeat copy. Then, we tested for association between each motif or groups of motifs, viewed as predictors, and the phenotype. This quantifies the impact of the set of common single or multiple base changes, against the 364-bp THE1B consensus sequence, on different recombination-related phenotypes. Motifs of interest were given a position relative to the 13-bp motif “CCTCCCTAGCCACG” previously identified (***Myers et al., 2008***) as predicting hotspot status in THE1B repeats, and closely matching the C-terminal end of the PRDM9 binding consensus sequence. This motif maps to position 261-274 in the THE1B consensus. To positionally map each motif, we used the mode of that motif’s first base position, relative to the first base of the motif CCTCCC[CT]AGCCA[CT]G, within THE1B repeat copies containing these two motifs. Phenotypes/annotations were either 0-1 (*e.g*. hotspot status, binding peak overlap), or quantitative (in the form of counts, for the H3K4me3 signal strength, specifically the number of reads observed). For the conditional testing we therefore used generalized linear models (GLMs) with either a binomial, or quasi-Poisson, underlying model as appropriate, as implemented in the “glm” library in R. For association testing we used Fisher’s exact test for association between 0-1 phenotypes and 0-1 motif occurrences, testing each motif separately. We performed different analyses catering for different phenotypes as appropriate, which we describe in subsequent sections.

#### Identifying motifs associated with PRDM9 binding to THE1B elements

We used our human PRDM9 ChIP-Seq data to annotate each THE1B element as bound or not bound by PRDM9. Specifically, an element was defined as bound if it overlapped an identified PRDM9 binding peak region (p<10^-5^). A substantial fraction of human THE1B elements (4392 of 20696,21%) were found to be bound. Recording binding across elements as a 0-1 vector, we successively fit GLMs of increasing complexity in a stepwise fashion, testing association between sets of motifs as regressors, and PRDM9 binding/non-binding as a response. In each model, we added a second matrix of regressors with entries defining which of the previously identified motifs CCTCCCCAGCCATG (matching the THE1B consensus sequence), CCTCCCTAGCCACG, CCTCCCTAGCCATG, or CCTCCCCAGCCACG, were present. These motifs are known to influence PRDM9 binding in THE1B elements (***Myers et al., 2008***). Including these additional regressors avoids false positive associations due to motifs whose presence/absence associates with these previously known determinants of PRDM9 binding. We restricted testing to only THE1B elements containing an exact match to one of these motifs, to avoid complexities due to cases of unusual PRDM9 binding to diverged THE1B sequences. Specifically, beginning with the model having only the 4 motifs above as predictors, we successively added in that new motif (of all 2021 possible motifs) maximally increasing the likelihood (as measured by the model deviance in the fitted GLM) of observed peak/non-peak status. We restricted the set of possible next motifs to those not strongly correlated (*r*^2^ <0.95) with the current set of included predictors, to avoid statistical artifacts due to near-complete motif co-occurrence and correlations, and to ensure a set of sufficiently independent predictors. Motifs were added in successively, until the conditional p-value of the next candidate motif was not significant (p<0.05) after Bonferroni correction for 2021 motifs tested. This yielded a final set of 17 motifs. We used the final joint GLM fit to estimate the joint effect of each motif on the probability of seeing a PRDM9 binding peak - in the binomial model, this is interpretable as the increase in the log-odds of a hotspot given each motif occurs, and taking into account the other motifs’ effects. We note that

1. Each of the 17 identified motifs by construction shows very strong evidence of influencing binding status, significant after Bonferroni correction for multiple testing (p<0.05).
2. All identified motifs map in - or close to - the predicted binding target region of PRDM9 based on our new set of motifs (***Figure 3***a). Different motifs act either to increase or decrease binding probability.

The estimated positions, effects and standard errors of each motif are shown in *Figure 3*a (top row). The full list of motifs themselves and estimated effect sizes is provided in ***Figure 3***-source data 1.

Identifying motifs impacting hotspot status conditional on PRDM9 binding presence/absence We annotated each THE1B element according to whether it overlapped a hotspot in a set of previously published human recombination hotspot positions (***Pratto et al., 2014***). That study examined meiotic DMC1 signal in male carriers of three different PRDM9 alleles labeled A-C. Alleles A and B bind similar target sites, and the B allele is studied here. We accordingly measured overlap only for hotspots detected in individuals whose PRDM9 alleles were both either A or B. We also annotated each THE1B element according to whether it overlapped an LD-based human hotspot (***HapMap, 2007***). These two annotations were highly correlated (p<10^-15^ by FET; odds ratio 25.6). Moreover, 1676 THE1B repeats overlapped Pratto *et al*. hotspots (2266 for LD-based hotspots), confirming that thousands of human hotspots localize in or near to THE1B elements. Having annotated THE1B repeats according to hotspot status, we used the same procedure as described above to test sequence motifs for association with hotspot status, separately for both hotspot sets. This analysis tests for evidence of association of different motifs with hotspot status, by influencing binding or other factors. We again used the same procedure, restricting to the set of THE1B elements defined as bound by PRDM9 above, to identify independent motifs associating with hotspot activity *conditional* on PRDM9 binding. We intersected motifs identified by these four analyses to identify a set of motifs robustly associating with hotspot occurrence, even given that measurable binding by PRDM9 occurs. (An initial comparison did not identify any evidence of motifs influencing one hotspot set differentially to the other, as might occur if *e.g*. female-specific influences on recombination rate exist within THE1B elements, and so we concentrate on this combined analysis.) First, we identified seven motifs with independent, significant evidence (p<0.05 after Bonferroni correction) of association with whether an LD-based hotspot was observed, conditional on binding by PRDM9 in our ChIP-Seq experiment. Separately, we identified four overlapping motifs with significant evidence of impacting the chance of being a Pratto *et al*. hotspot, conditional on binding by PRDM9 in our ChIP-Seq experiment. Using the set of 9 unique motifs, we then fit a series of generalized linear models to jointly test for association of a 0-1 matrix with 9 columns indicating motif presence/absence on (i) LD-based hotspot status, (ii) Pratto-based hotspot status in human males, and (iii)-(iv) the same conditional on PRDM9 binding, *i.e*. restricting testing to the set of THE1B elements overlapping a PRDM9 binding peak. In each model, we continued to include as regressors the previously identified 14-bp motifs influencing PRDM9 binding, and restrict testing to elements containing one of these motifs. Following this joint analysis, seven motifs show (a) p<0.05 (Bonferroni corrected p-value) for hotspot occurrence given binding, for at least one of the Pratto hotspot set and the LD-based hotspot set and (b) p<0.05 (nominal p-value) for all four tests, *i.e*. evidence of influencing hotspot status regardless of hotspot definition used, and both conditional and unconditional on PRDM9 binding. All but one of these motifs associate (p<0.05 after Bonferroni correction) with hotspot occurrence *unconditionally* also. We considered these seven motifs to form a set of independent, robust and consistently detected influences on hotspot status within THE1B repeats. For example, the motif “ATCCATG” shows p<0.05 after Bonferroni correction for all of (i-iv) above. Specifically, testing this motif (conditional on previously identified 14-bp PRDM9 binding motifs) at all THE1B repeats, without conditioning on PRDM9 binding, showed p=4.1×10^-11^ for association with DMC1 hotspots and p=5.9×10^-13^ for association with LD-based hotspots and odds ratios of around 0.5. This means that its impact on hotspots cannot be mediated via any biases in our ability to measure binding in HEK293T cells. The other two motifs of nine may associate with hotspot status, but were conservatively excluded because they showed no evidence (p>0.05) for unconditional evidence of association with hotspot status. They were removed in case their effect is mediated through properties of PRDM9 binding, specific to HEK293T cells. The detailed results of this conditional testing are given in *Figure* 3-source data 1, and were used to produce the first two rows of *Figure 3*a.

Identifying motifs associated with previously measured H3K4me3 signal strength in testes A previous human study measured levels of H3K4me3 in testes (***Pratto et al., 2014***). Although PRDM9 deposits H3K4me3 on binding, other proteins are capable of inducing this mark, and H3K4me3 occurs, for example, at many human promoters independently of PRDM9. We sought to identify sequence features impacting male meiotic H3K4me3 in THE1B elements, whether bound by PRDM9 or not bound. We “force-called” H3K4me3 as a quantitative phenotype at each THE1B element, and here test for association with the total number of reads observed across two replicates within 1kb of the center of the element. We split the THE1B elements into two sets, those with potential PRDM9 binding (the “bound set”) and a set robustly evidenced to not be bound by PRDM9 (the “unbound set”). For the bound set, we took the subset of THE1B elements containing an exact match to one of the 14-bp motifs CCTCCC[CT]AGCCA[CT]G, and overlapping a PRDM9 ChIP-seq peak. For the unbound set, we conservatively used the set of THE1B repeats remaining after removing as potentially bound by PRDM9 any repeat matching CCTCCC[CT]AGCCA[CT]G, or overlapping a PRDM9 binding site in our HEK293T cells, or overlapping an LD-based hotspot, or overlapping any Pratto *et al*. hotspot. The remaining THE1B elements contain no good match to the PRDM9 binding motif, and further show no evidence of any PRDM9-associated phenotype (binding or hotspot status). We then performed testing exactly as for the 0-1 annotations, to identify independent motifs associating with H3K4me3 level in each set. The only difference in each case was the GLM used (quasi-Poisson model). Notably, in the non-bound set of THE1B repeats, we are then testing for sequence features associating with H3K4me3 levels, independent of PRDM9. Similarly to PRDM9 binding motifs, the identified motifs are likely to causally influence histone modifications including H3K4me3 levels (and as described in the main text and below, they also associate with H3K9me3 and H3K4me3 in somatic cells, and potentially other modifications), but through initially unknown biological mechanisms. In the bound set, both PRDM9-dependent and PRDM9-independent sequence features might be identified. The testing of non-bound regions identified 18 distinct motifs after Bonferroni correction of significance level, mapping throughout the THE1B consensus sequence and associated with both increases and decreases in measured H3K4me3 signal. The estimated positions, effects and standard errors of each motif were used to construct ***Figure 3***a and ***Figure 3***d. The full list of motifs themselves and estimated effect sizes is provided in ***Figure 3***-source data 1. We note that all the motifs, except possibly one, map *outside* the PRDM9 target motif region, consistent with a role distinct from PRDM9. Further supporting this idea, 15/18 motifs show effects in the same direction for the “bound set” testing of the smaller, and so statistically less well powered, collection of PRDM9 bound repeats, suggestive of a continuing impact even if elements are also bound by PRDM9; although this reached significance in only 4 cases (p<0.05, with p<0.0001 for the strongest signalled motif), this can be explained by the dominant impact of PRDM9 binding on H3K4me3 for this set, as well as the smaller sample size.

#### Overlaps and correlations between recombination-related measures

The above procedures produced three partially overlapping sets of motifs that are highly significantly associated with PRDM9 binding, hotspot occurrence (measured by LD or DMC1) at sites bound by PRDM9, and H3K4me3 marks formed dependent and independently of PRDM9, respectively. We compared the sets of motifs identified – independently, using different phenotypic measures and often different sets of THE1B repeats – for overlaps. Given each set of motifs, we used the same procedures as described above to test the other measures, in order to examine whether the same features might have directional effects for the other measures and phenotypes. The results are shown in *Figure* 3-source data 1 and described briefly in the main text. Overall, we found the following:

1. The determinants of PRDM9 *binding* we identified are found exclusively within the region directly contacted by the zinc fingers of PRDM9, or immediately adjacent (<10bp). All influences on binding mapped within a region from −22 bp to + 14 bp relative to the motif CCTCCCTAGCCAC, in every case overlapping by the predicted PRDM9 binding motif within THE1B. While a previous report suggested influences on PRDM9 binding outside the binding region (***Grey et al., 2017***), these are not strongly evidenced here, although the motif from +14 bp to 22 bp inclusive extends slightly beyond the region bound by PRDM9. Finally, the motif CCTCCTT (p=9.94×10^-5^) is the most significant motif failing to reach Bonferroni significance, mapping just upstream of the region directly predicted to be within the binding region (−29 bp to −23 bp inclusive), suggesting there may be a weak role for sequence <10 bp away but not overlapping the identified motif itself.
2. Changes in DNA sequence throughout the roughly 40-bp PRDM9 binding target region (17 motifs) impact meiotic recombination, and recombination “heat” as well as H3K4me3 deposition seem to depend in a simple directional manner on binding. In general almost all (two exceptions discussed below) of 17 motifs impacting binding impact H3K4me3 at the bound sites in the same direction in human testes, *i.e*. during meiosis (where PRDM9 is expressed). Moreover, with the same 2 exceptions, all had a trend for measured recombination in the same direction when measured by LD and/or DMC1. For multiple motifs these associations were highly significant (***Figure 3***-source data 1). This finding is not unexpected but confirms the biological relevance of precisely and directly measuring binding, even in HEK293T cells.
3. As well as the above, and surprisingly, we identified a large number of motifs (18 reaching Bonferroni-corrected significance), associating with H3K4me3 signal strength in human testes at regions not bound by PRDM9. They map throughout the THE1B repeats, with only one overlapping the PRDM9-bound region. These motifs each have rather weak signals for the H3K4me3 signal compared to (for example) PRDM9 binding. However as we discuss below, the same motifs each show (stronger) impacts on H3K9me3 deposition within a large collection of cell types, and so it may be that histone modifications other than H3K4me3 drive the links between these motifs and meiotic recombination (see below), and our H3K4me3 signals appear as secondary biological markers of this stronger effect. We therefore call these “non-PRDM9 H3K9me9/H3K4me3” motifs.
4. We observed a strong, consistent, counter-directional correlation with non-PRDM9 H3K9me9/H3K4me3 motifs and hotspot activity. In THE1B elements, the sequence features increasing H3K9me9/H3K4me3 measured signals decrease recombination rate, in a seemingly simple linear fashion, and (less strongly) the opposite holds for decreases in H3K9me9/H3K4me3. First, of the seven new motifs identified to influence whether hotspots occur given binding in THE1B, three occur within the PRDM9 target motif, and are explained via direct changes on binding strength, in the expected direction. The remaining four motifs are outside the PRDM9 target motif. All of these are strongly associated (p<10^-60^ for ATCCATG) with non-PRDM9 H3K9me9/H3K4me3, in the opposite direction to the recombination association (***Figure 3***-source data 1). Conversely, testing influence of the 18 non-PRDM9 H3K9me9/H3K4me3 motifs on (i) PRDM9 binding, and (ii) LD/DMC1 hotspot formation, we found no particular association with PRDM9 binding itself, and no overlap with the set of motifs identified to influence PRDM9 binding. However, for 17/18 motifs we observed associated with increased/decreased H3K9me9/H3K4me3 levels, they were associated with decreased/increased probability of hotspot occurrence for each of LD-based hotspots and DMC1-based hotspots. The only exceptions in terms of direction showed non-significant trends, in different directions for the two sets of hotspots, so might be explained by statistical noise. Multiple motifs show significant evidence of significantly altering hotspot probability (***Figure 3***a; ***Figure 3***-source data 1). In particular, the most significant motif, associated with increased non-PRDM9 H3K9me9/H3K4me3, was again “ATCCATG” (p=4×10^-26^). This motif has no association with PRDM9 binding in our experiments (p>0.1) but overwhelming evidence of reducing hotspot probability at these binding sites and is in the motif set identified independently as associating with hotspot occurrence (p<10^-4^ for association with hotspot occurrence given binding, for each of DMC1 and LD hotspots).
5. As mentioned above, two motifs, “TTGTGAG” and “CCATGAT”, have significant impacts on both PRDM9 binding and meiotic recombination, but in *opposite* directions. This unusual property might in principle reflect subtle differences in binding properties between PRDM9 alleles A/B or in different cellular environments (HEK293T cells vs. cells where PRDM9 is natively expressed). However a simpler explanation given the above is offered by the fact that both motifs have a weak positive association with non-PRDM9 H3K9me9/H3K4me3 independent of PRDM9 binding (p<0.005 in each case). Thus there may be competition for these motifs involving an increase in PRDM9 binding, but within an environment where other histone modifications they cause make a hotspot less likely, plausibly resulting in a predicted decrease in hotspot probability *given* binding, as observed. Thus the complex patterns we observe comparing thousands of sequence motifs across thousands of THE1B elements for four different recombination-related phenotypes may actually be highly parsimoniously explained by a simple but surprising phenomenon: PRDM9 binding and PRDM9-induced H3K4me3 deposition dramatically increase hotspot probability, but PRDM9-independent H3K4me3 and/or H3K9me3 (see below) dramatically inhibit recombination, downstream, even where PRDM9 is able to bind the THE1B repeat.

#### Examining the impact on recombination of non-PRDM9 H3K9me3/H3K4me3 motifs in THE1B

To explore this signal, we plotted the estimated effect on H3K4me3 signal strength (log-fold increase on measured H3K4me3 signal) of each motif versus the average impact on recombination (measured by log-odds of a hotspot), in ***Figure 3***-S1c. This revealed a striking, essentially linear, negative trend (p<10^-16^ by rank correlation; rank correlation −0.85). Given these consistent marginal effects, we next examined how much influence these motifs have jointly, on whether hotspots occur or otherwise at THE1B repeats bound by PRDM9. Conceptually we imagine PRDM9-induced H3K4me3 increasing recombination, but other motifs that increase the non-PRDM9 H3K9me9/H3K4me3 signal, instead reducing recombination - in “opposition”. Although we can use the H3K4me3 data in the appropriate tissue (testes), the signals obviously and unfortunately conflate, and cannot separate whether these data measure H3K4me3 deposited by PRDM9. However, we *can* separate them by using our identified motifs. We used (only) the DNA sequence of each THE1B repeat to predict the non-PRDM9 H3K9me3/H3K4me3 for that repeat. This is expected to negatively correlate with recombination from the above findings. It appears as if PRDM9 binding in general does not alter the effect of non-PRDM9 H3K9me3/H3K4me3 motifs (***Figure 3***-source data 1), so this DNA-sequence-based measure is likely to remain relevant in those repeats also bound by PRDM9. Indeed: in the column “H3K4me3 at bound THE1B elements” of ***Figure 3***-source data 1, almost all the identified non-PRDM9 H3K9me9/H3K4me3 motifs have impacts in the same direction (rank cor-relation p=0.00036) for the unbound repeats, including *e.g*. the motif ATCCATG (p=2×10^-5^). In detail, for each element we calculated a separate “positive” and “negative” motif score (relative to a conceptual highly diverged THE1B element containing none of the motifs) for only motifs acting in those directions, summing the values given in column “N” of ***Figure 3***-source data 1 across motifs present in that repeat copy. We fit a regression model (Poisson GLM as above) and found both scores to be highly significantly associated with hotspot occurrence (p=9.9×10^-6^ and p=1.7×10^-7^ respectively) in opposite directions, though with slightly different coefficients. We combined the scores by adding them, downweighting/tempering the negative part of the non-PRDM9 H3K9me9/H3K4me3 signal by 2.3637/3.4842, the ratio of regression coefficients. This yields a single prediction value of the non-PRDM9 component of H3K4me3 per THE1B repeat. To visualize the impact of non-PRDM9 H3K9me9/H3K4me3 signal on hotspots (***Figure 3***d), restricting our analysis to the set of elements defined as bound by PRDM9 as above, we binned their predicted non-PRDM9 component of the H3K9me3/H3K4me3 signal into 10 equal quantiles. For each quantile, we plotted the (log-fold) mean H3K9me3/H3K4me3 change, against the probability of a hotspot given binding. It should be noted that these correspond to a rather modest range of predicted H3K4me3 changes - for example the 95% upper quantile of the summed positive influences on H3K4me3 corresponds to just a 1.3-fold increase in signal over background. It is difficult to quantify how strong this is biologically given noise in the H3K4me3 assay, but a helpful comparison might be that the single motif CCTCCCTAGCCAC confers a >2-fold increase in H3K4me3 signal in testes within bound PRDM9 repeats even conditional on binding occurring, so it seems likely that H3K4me3 differences made by these motifs are modest - and require caution in interpretation, given the same motifs also associate with much stronger H3K9me3 level differences (see below). Strikingly and nevertheless, as a group these motifs produce a very large and consistent impact on hotspot probability, almost identical for the DMC1 and LD-based hotspot sets. Hotspot probability reduced almost 3-fold, from 35% to 13%, as non-PRDM9 H3K9me3/H3K4me3 increased. Thus, complex non-PRDM9 sequence factors operate in combination to collectively determine whether hotspots occur at THE1B repeats.

#### General suppression of meiotic recombination but not PRDM9-associated H3K4me3 deposition, by the motif ATCCATG

We investigated whether non-PRDM9 H3K9me9/H3K4me3 sequence motifs reduce recombination by preventing PRDM9 from binding DNA and therefore recruiting DSBs, or instead act downstream of PRDM9 binding. For the most significant motif “ATCCATG” we were able to test this by plotting mean LD-based and DMC1-based recombination rate, and H3K4me3 level in human testes, for a 10-kb region (500 bp window slide 250 bp across region) centered on the THE1B repeat. We calculated and plotted each mean separately, grouping THE1B repeats according to whether they contain different PRDM9-bound motifs of the form CCTCCC[CT]AGCCA[CT] resulting in progressively stronger binding by PRDM9, and then either contain, or do not contain, the motif “ATCCATG” (***Figure 3***b). As expected, the recombination signal increases steadily with closeness of the match to the PRDM9 consensus sequence CCTCCCTNNCCAC. Conditional on this closeness, presence of the motif ATCCATG always and strongly reduces mean recombination rate by around 2-fold. Even where no PRDM9 binding motif is present inside the THE1B repeat itself (***Figure 3***b, cyan lines) there is a statistically significant (p<10^-10^) suppression of mean recombination rate *below background* when the motif ATCCATG occurs, at a scale of approximately 1-2 kb in each direction. Thus, the motif ATCCATG within THE1B repeats appears to be a strong general local suppressor of human recombination, and is able to suppress recombination when PRDM9 binds the usual motif in THE1B, and nearby hotspot occurrence more widely. Moreover, this suppression acts over reasonably broad scales. In contrast to their different effects in recombination, while the H3K4me3 signal consistently increases with closeness of the match to the PRDM9 consensus sequence CCTCCCTNNCCAC, it is also higher when the non-PRDM9 motif ATCCATG is present, with no evidence that this motif suppresses PRDM9-dependent H3K4me3 deposition *in vivo*. It appears that PRDM9 binding, and ATCCATG-driven histone modifications, act additively and perhaps independently. Therefore, this single non-PRDM9 motif must play a strong suppression role in a high proportion of the THE1B repeats where it is present. Likely, this suppression acts in both males and females, because DMC1 rate estimates are for males only, while LD-based rate estimates are sex-averaged and reflect mainly ancient crossovers.

#### Association testing the full landscape of histone modifications in THE1B repeats across ROADMAP cell lines

The ROADMAP consortium (***Kundaje et al., 2015***) previously measured multiple histone modifications and other molecular phenotypes across 125 diverse human somatic cell types. These were used to partition the genome into 15 different domains characterized by combinations of histone modifications: TssA, TssAFlnk, TxFlnk, **Tx, TxWk**, EnhG, **Enh, ZNF/Rpts, Het**, TssBiv, BivFlnk, EnhBiv, **ReprPC, ReprPCWk, Quies**. Eight of these states (in bold) occur over 8 times across the 20696 THE1B repeats on average and were examined. We first identified the ROADMAP domain inference for each THE1B repeat in each of the studied cell types. For each domain type and each cell type, we identified *de novo* a set of motifs associating with that domain in that cell type, by exactly repeating the analysis approach we used for hotspot status, as described above. We used a p-value cutoff of 2.5 × 10^-8^, to Bonferroni correct for the total of 125×8×2021 tests performed. The full resulting set of 1571 identified ROADMAP motifs and details is given in ***Figure 3***-source data 1. The motifs cover all 8 domain types, and every cell type has at least three, and up to 36, different motifs. Thus, as in meiosis THE1B repeats possess a diverse set of independent motifs associated with many different histone modifications (including H3K9me3, H3K27me3, H3K4me3, H3K36me3, among others) in THE1B elements. Although our main focus here is on correlating results with recombination rates, the collection of motifs is of biological interest in itself. We grouped highly co-occurring (and typically overlapping) motifs, collapsing motifs whose correlation (in which THE1B element each motif occurred in) was >50% until no further grouping was possible. This resulted in a set of 67 distinct “summary” motif groups, whose results are summarized in ***Figure 3***-source data 1, and which span much of the THE1B sequence. Previously, two papers have identified transcription factors DUX4 (***Young et al., 2013***) and ZBTB33 (***Wang et al., 2012***) as preferentially binding particular motifs within THE1B elements. Ordering motifs by how many cell types they are active in, of the top four motif groups identified, the top motif corresponds to the DUX4 consensus binding sequence and associates DUX4 binding with the two “Tx” (transcription) domains, associating the occurrence of this motif with only a signal of elevated H3K36me3 (***Kundaje et al., 2015***), ubiquitously across somatic cell types. Despite this, and interestingly, this motif was NOT identified as influencing H3K4me3 in testes, nor with any impact on meiotic recombination. Similarly, the fourth motif is a match to the ZBTB33 (Kaiso) target motif, associating this motif with the occurrence of both Tx (*i.e*. H3K36me3) and “ReprPCWk”; polycomb modifications, exhibiting enrichment of the H3K27me3 histone modification. The latter modification was previously associated with ZBTB33 binding, while the former represents a distinct modification associated with the same motif. The second motif group exactly matches the motif CCGCCAT which is the consensus binding target of YY1 and in THE1B repeats shows a similar Enrichment signal to the DUX4 motif. The final motif of the top 4 identified was precisely the motif ATCCATG, which we identified above and found to strongly reduce recombination rate where present. Across 110 categories and cell types, this motif was identified, and unlike the above motifs, showed enrichment for both the “Het” and “ZNF/repeats” categories. These are characterized by elevated H3K9me3, which marks “constitutive” heterochromatin or inactive DNA with widespread methylation of CpG dinucleotides, and in the second case, by elevated H3K36me3 also, which instead marks active regions, including transcribed regions. Given this, we compared all 18 motifs associating with H3K4me3 signal strength in human *testes* at regions not bound by PRDM9 (called non-PRDM9 H3K9me3/H3K4me3 motifs above) – and which show a consistent association with meiotic recombination in the opposite direction. Remarkably, 14 of the 18 motifs coincided with 14 of the 67 motif groups, indicating that these motifs (unlike PRDM9) appear to associate with histone modifications in somatic cells. Moreover, all 14 coinciding motifs lie within the subset of 34 motif groups associating with, in at least one cell type, the same heterochromatin category as the motif ATCCATG, a highly significant enrichment (p=2.3×10^-5^ by FET). This suggests a common cause for these diverse motifs - across many different cell types, they associate with increasing heterochromatin (H3K9me3, and as described above, and below, H3K4me3), while increases in H3K9me3 accompany increases in average H3K4me3 in testes, and decreases in meiotic recombination. Indeed, although we found 33 different motif groups associating with exclusively non-heterochromatin ROADMAP cellular domains, for example transcribed regions (Tx), or the polycomb repression-like state, none of these showed an impact on either H3K4me3 in testes, or meiotic recombination, despite (for example) high testes expression of DUX4 (***Young et al., 2013***). This implies a potential causal relationship between recombination and H3K4me3/H3K9me3, rather than the other marks studied by ROADMAP, within THE1B repeats. Looking across cell types, the overlap between motifs influencing THE1B H3K4me3 in testes and the heterochromatin state varies strongly between 0 and 10. The top cell types (***Figure 3***-source data 1) in increasing overlap were the following cell lines: ES-I3 Cells, hESC Derived CD184+ Endoderm Cultured Cells, hESC Derived CD56+ Mesoderm Cultured Cells, Primary monocytes from peripheral blood, Primary hematopoietic stem cells G-CSF-mobilized Male, Fetal Intestine Small, HUES48 Cells, HUES6 Cells, iPS-20b Cells and HUES64 Cells. This list is dominated by embryonic stem cells (ESCs), their derivatives, and induced pluripotent stem cells. These cell types therefore behave most similarly to the properties we observe for both meiotic recombination, and H3K4me3 in testes. Although the genomic “domain” annotation is informative, we further directly analyzed histone modification enrichment values for all seven core “ROADMAP” studied motifications (***Kundaje et al., 2015***) in two of the embryonic stem cell (ESC) types showing the strongest overlap; HUES6 Cells (E014 in ***Figure 3***-source data 1) and HUES64 cells (E016). Using each histone modification in turn as a phenotype, we tested jointly (using the same Poisson GLM framework as previously) for an association of the set of 18 motifs influencing meiotic H3K4me3 on that modification in the ES cells. We tested whether (i) each motif showed a significant impact in ESC cells, and (ii) for correlation in the estimated *effect size* in ES cells to that in testes H3K4me3, to examine whether there is a concordant effect across cell types. Results for both ESC types were highly concordant (***Figure 3***-S1d). For (ii), in HUES6 cells by far the strongest correlations in estimated effect size were seen with two marks; H3K4me3, and H3K9me3, with similar very strong positive rank correlations >90% (p<10^-16^). These correlations are so high that within noise, it appears many or most motifs have identical impacts across these cell types. Nominally significant negative correlations of around −0.5 were also seen for alternative histone modifications at the same residues: H3K4me1 and H3K9ac (0.01<p<0.05), potentially explained by their absence when the other modifications are present. 9 of the 18 motifs were significant at p<0.05 for H3K4me3, and remarkably 15 of 18 are significant for H3K9me3 in HUES6 cells, all in the same direction as testes H3K4me3 (***Figure 3***-S1d), from these fully independent data. Taken together, these results overwhelmingly imply that all, or almost all, the motifs which are responsible for elevated H3K4me3 in THE1B in testes, operate similarly or identically to elevate H3K4me3 in other tissues and cell types, particularly embryonic stem cells. Further, they are also - and considerably more strongly (***Figure 3***c) - associated with H3K9me3 elevation in the same cell types. Therefore, we describe these motifs as non-PRDM9 H3K9me3/H3K4me3 motifs to reflect this. We note that this does not directly imply these marks are *established* in ESCs and other cells and they might be inherited in these cell types from progenitors. However these non-PRDM9 influences on recombination, unlike PRDM9-induced H3K4me3, clearly operate rather widely across cell types. It is perhaps surprising that H3K4me3 and H3K9me3 should show these consistent impacts in the same directions, and across diverse motifs within THE1B repeats; such a pattern was though seen previously across human repeats (***Kundaje et al., 2015***) and so might operate more widely. Unsurprisingly given our results, across all 20696 THE1B repeats we studied, the enrichment for these two marks is highly correlated (rank correlation 61% in HUES6 cells, the highest for any pair of marks), so the same individual THE1B repeats show (often weak) enrichment for both marks, although this does not necessarily imply co-occurrence in the same individual cells. Potential causes of these histone modifications are discussed in the main text.

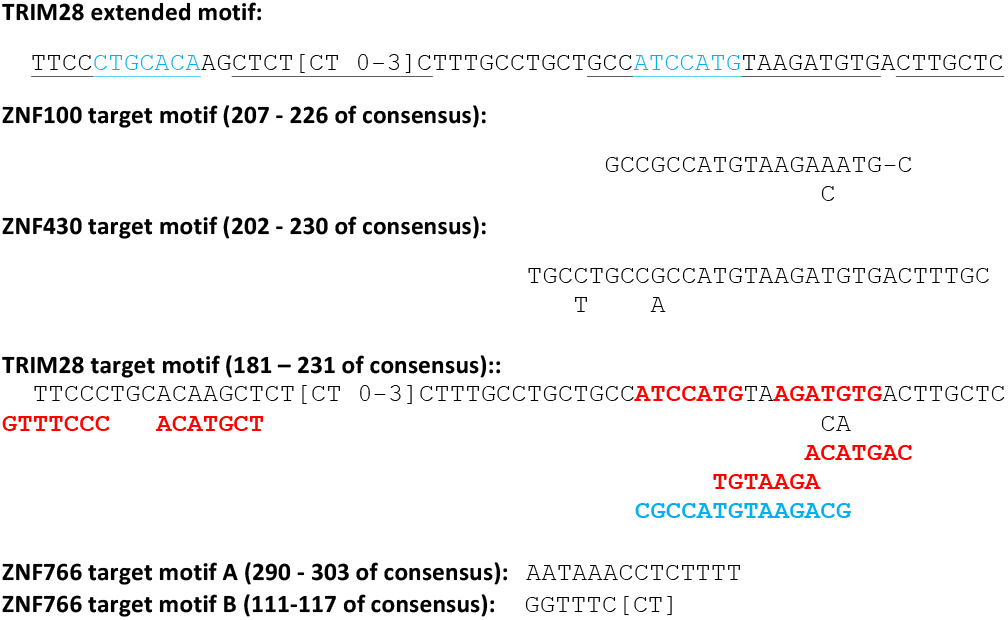

#### Identifying motifs associated with binding of KRAB-ZNF genes, and TRIM28 recruitment, atTHE1B repeats

The above approach describes a method to identify sequence motifs within all or a subset of THE1B elements influencing 0-1 hotspot status. We applied the identical approach to attempt to identify binding motifs for three KRAB-ZNF proteins enriched for PRDM9 binding (***Imbeault et al., 2017***; Michael Imbeault, personal communication): ZNF100, ZNF430 and ZNF766. For each we first identified instances of binding peaks of each protein within 500 bp of the centers of THE1B elements, and then identified motifs. We did the same for TRIM28, a protein recruited by the KRAB domains of many KRAB-ZNF proteins, and assayed in H1 human embryonic stem cells (***Imbeault et al., 2017***). In the first three cases, the identified motifs cluster and could be mapped to specific regions of THE1B, shown in ***Figure 3***a and also described below. In the case of TRIM28 the signal is expected to be a superposition of sites of binding by different KRAB-ZNF proteins, complicating interpretation; indeed we identified 16 motifs, mapping throughout THE1B elements. The top-scoring motifs were TCCCTGC and CCATGTA. These heavily overlapped 2 of the 4 motifs altering (and in both cases decreasing) the probability of hotspot occurrence, including the highly significant motif ATCCATG. Therefore, we conditioned on the latter motif occurring and repeated our motif-finding for the resulting subset of THE1B repeat elements, reasoning that such TRIM28 peaks might be bound by a single protein with a well-defined target motif. Indeed, this analysis revealed a set of 7 motifs, all within a contiguous region of length 57 bp and covering the 41 bases in bold and underlined below, mapping to the region 181-231 of the THE1B consensus sequence. The resulting extended “TRIM28” target motif is shown below. There is some spacing variability in the first half of this motif among bound copies because of the variable number of copies of “CT” found in this region. This motif incorporates and links the hotspot-influencing motifs ATCCATG and CTGCACA (highlighted in blue). Moreover, it overlaps several additional motifs associated with (increasing, red below) non-PRDM9 H3K9me3/H3K4me3. Finally this motif is disrupted by several motifs associated with decreasing (blue below) non-PRDM9 H3K9me3/H3K4me3. These overlaps are highlighted in the above figure, which gives results for all four motifs.

As shown in the above alignment figure, we also identified two similar target motifs for binding of ZNF766 mapping to different parts of the THE1B repeat consensus. The previously unknown extended “TRIM28” motif above is therefore a recombination coldspot motif, and simultaneously a motif, including the motif “ATCCATG” and others, for TRIM28 recruitment, H3K9me3 deposition, and weaker H3K4me3 deposition, at the same locations. Moreover it appears that binding in THE1B repeats and elsewhere by each of four further zinc finger proteins ZNF430, ZNF100, ZNF766 is recruited by other motifs for decreased recombination rates, in a manner highly dependent on the *cis* sequences near PRDM9 binding sites inside THE1B repeats.

#### Testing for a general association between KRAB-Zinc-finger protein binding and TRIM28 recruitment and recombination at PRDM9-bound sites

Given that binding by KRAB-ZNF genes and TRIM28 recruitment offers an explanation for the ability of particular sequence motifs in THE1B to increase H3K9me3 and H3K4me3 and yet decrease recombination rates, while not preventing PRDM9 binding, we tested if this property were more general. Across 235 recently studied KRAB-ZNF genes and TRIM28, we first identified their ChIP-seq binding sites falling within 500 bp of our PRDM9 binding sites, after excluding PRDM9 binding sites at pre-existing H3K4me3 peaks, near TSS, or overlapping DNase HS sites (where our other results show hotspots to be less likely; including these regions strengthened but did not alter the below results). We then studied those proteins with at least 30 peaks overlapping our binding sites (other proteins showed similar overall patterns though we lacked statistical power to examine them individually). We used the binary GLM framework described above to perform association testing for each protein separately between occurrence of that protein binding the genome within PRDM9 binding sites, and whether those binding sites become hotspots. We included our measured PRDM9 binding strength, and local GC-content within the PRDM9 binding site, as co-regressors. The results are shown in ***Figure 3***e; the estimated effect of KRAB-ZNF binding was negative in all but one case, and significantly negative impacts of binding on recombination (p<0.05) was seen for 27 proteins (TRIM28 being the most significant) examined despite the typical low overlap of individual KRAB-ZNF genes with PRDM9 binding sites. Among the genes with significant negative impacts were each of the four analyzed above that bind THE1B repeats, and where we were able to identify connections to their binding target sequences. For each protein we also tested for association with H3K9me3 in HUES-64 cells, with identical predictors. Instead of hotspot status, the response variable was now mean H3K9me3 enrichment in the 1 kb surrounding the PRDM9 binding peak center, after quantile normalization and now fitting an ordinary linear model. The resulting values were used to color ***Figure 3***e. The large majority of KRAB-ZNF genes examined show positive correlations between binding and H3K9me3 placement, as expected (***Imbeault et al., 2017***).

## Data Availability

Sequencing reads, genome-wide fragment coverage depth, peak calls, and differential gene expression files are available with GEO accession GSE99407.

## Acknowledgments

We would like to thank Jonathan Flint for providing bench space and reagents, as well as Julian Knight, Benjamin Davies, Peter Donnelly, Anjali Gupta Hinch, Robert W. Davies, and Catherine M. Green for their helpful guidance and feedback. We thank the Oxford Genomics Centre for generating the sequencing data.

## Supplementary Figures

**Figure 1-Figure supplement 1.**
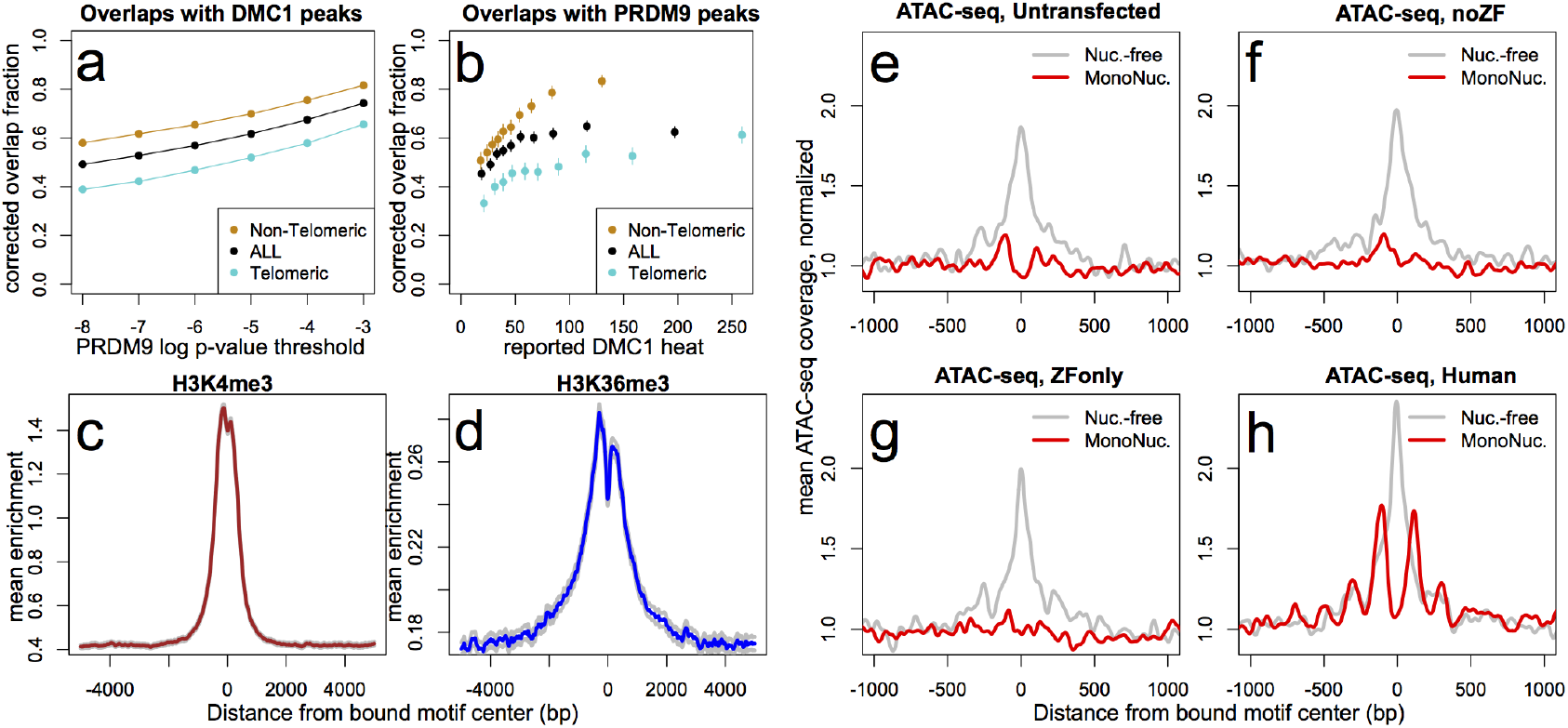
DMC1, H3K36me3, and ATAC-seq signals surrounding human PRDM9 peaks. **a**: A comparison our autosomal PRDM9 peaks, called at various p-value thresholds ranging from 10^-8^ to 10^-3^ (minimum peak separation 250 bp), to a set of published DSB hotspots corresponding to the human A allele (from a set of 18,343 “Intersect” DMC1 hotspots found in multiple individuals, filtered to remove hotspots wider than 3 kb ***Pratto et al., 2014***). Hotspots were further split into subsets occurring within 15 Mb of a telomere (turquoise) or not (orange). “Overlap” requires a PRDM9 peak center to fall within a reported DMC1 hotspot interval, and overlap fractions were corrected downward to account for chance overlaps (see Methods and Materials). **b**: DMC1 hotspots were split into decile bins by reported DMC1 heat, and the proportion of hotspots in each bin overlapping one or more of our PRDM9 peaks is indicated (error bars represent two standard errors of the proportion). **c**: Profile plot showing the mean H3K4me3 enrichment (measured in HEK293T cells transfected with human PRDM9) at bound human motifs conditioned not to have any H3K4me3 enrichment in untransfected cells. Grey lines indicate 2 standard errors of the mean. (smoothing: ksmooth, bandwidth 25) **d**: Profile plot showing the mean H3K36me3 enrichment (measured in HEK293T cells transfected with human PRDM9) at bound human motifs conditioned not to have any H3K36me3 enrichment in untransfected cells. Grey lines indicate 2 standard errors of the mean. NB: absolute enrichment values cannot be compared across samples. (smoothing: ksmooth, bandwidth 25) **e-h**: ATAC-seq profile plots surrounding a set of the ~15,000 strongest human PRDM9 ChIP-seq peaks (filtered to require a motif match and to not overlap an annotated DNase hypersensitive site), across 4 different transfection samples. “Coverage” here refers to the frequency with which an ATAC-seq fragment center occurs at each position, such that “Nuc.-free” coverage tracks the centers of nucleosome-depleted regions, and “MonoNuc.” coverage tracks the centers of single nucleosomes. Coverage values are normalized to the mean values observed between 1500 and 3000 bases away from each site, as a measure of background, and smoothed (ksmooth bandwidth = 50). The human-transfected cells show strongly phased nucleosomes centered at ~100 bp to either side of the motif and an elevated signature of nucleosome depletion at the center (**h**), when compared to the three controls (**e,f,g**). The ZFonly result (**g**) suggests that the ZF domain alone is insufficient to produce this nucleosome phasing. These data also suggest that PRDM9 binding is favored in nucleosome-depleted regions.

**Figure 1-Figure supplement 2.**
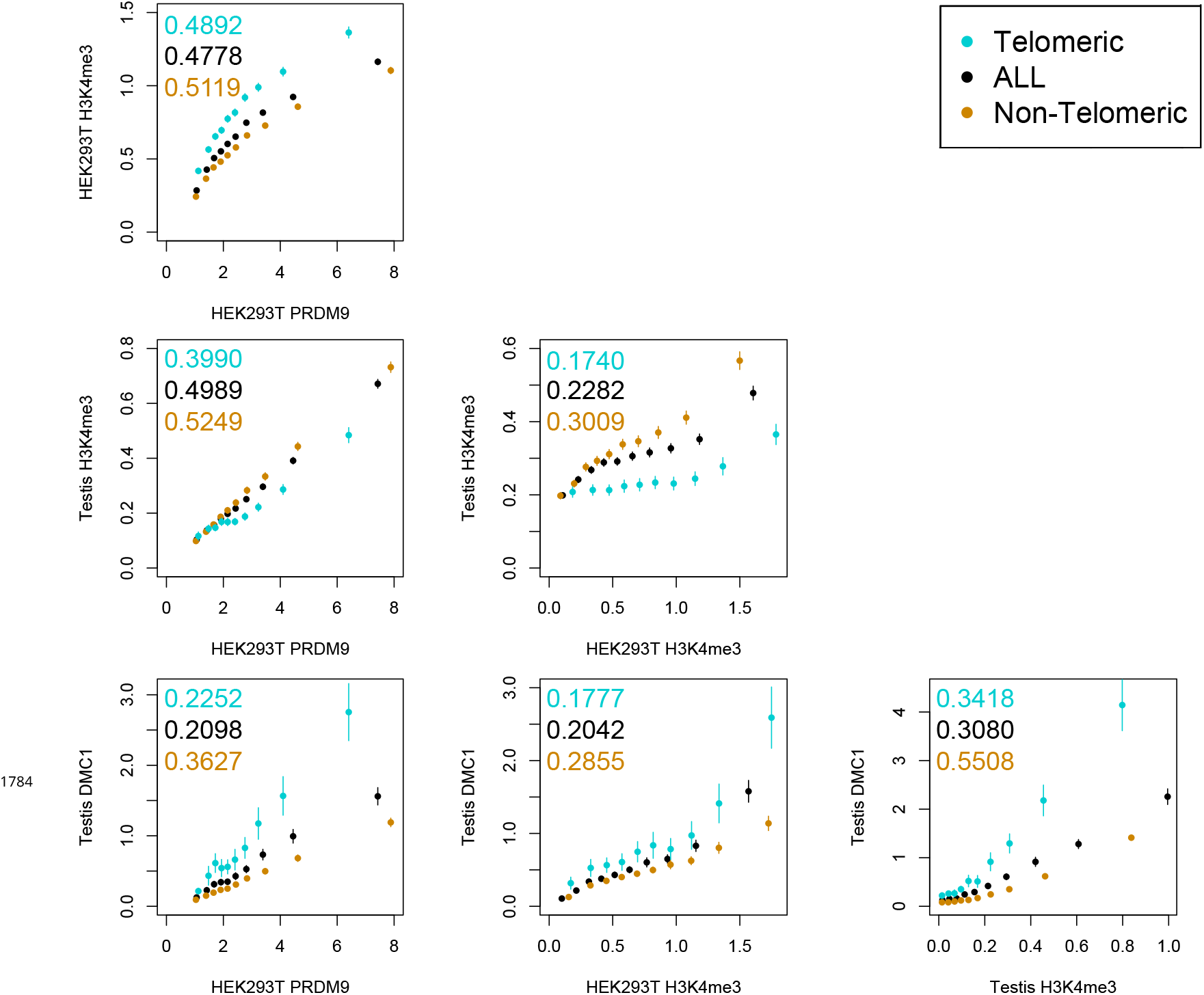
Comparison of PRDM9 and H3K4me3/DMC1 enrichment values. H3K4me3 ChIP-seq data from transfected HEK293T cells (this study) and H3K4me3/DMC1 data from testes (***Pratto et al., 2014***) were force-called in a 1-kb window centered on each PRDM9 binding peak center (p<10^-6^, minimum peak separation 1000 bp) to provide an enrichment value for each H3K4me3/DMC1 sample at each PRDM9 peak. Peaks were further split into subsets occurring within 15 Mb of a telomere (turquoise) or not (orange). Pairwise comparisons plot the mean force-called enrichment value of each sample (*y* axis) in each enrichment decile bin of each other sample (x axis). Points are positioned at the median value of each decile and error bars represent two standard errors of the mean. Raw Pearson correlation values are printed on each plot. All comparisons show a significant positive correlation (p<2×10^-16^). Peak windows with fewer than 5 input reads from cells or testes were filtered out, to improve enrichment estimates, and windows with excessive genomic coverage (in the top 0.1%ile) or IP coverage (>500 combined fragments) were removed to avoid outliers due to mapping errors. PRDM9 peaks overlapping H3K4me3 peaks from untransfected cells were removed, leaving 37,188 peaks passing all filters. Interestingly, we observe an enrichment of H3K4me3 in telomeric peaks in our HEK293T cells but not in testes.

**Figure 1-Figure supplement 3.**
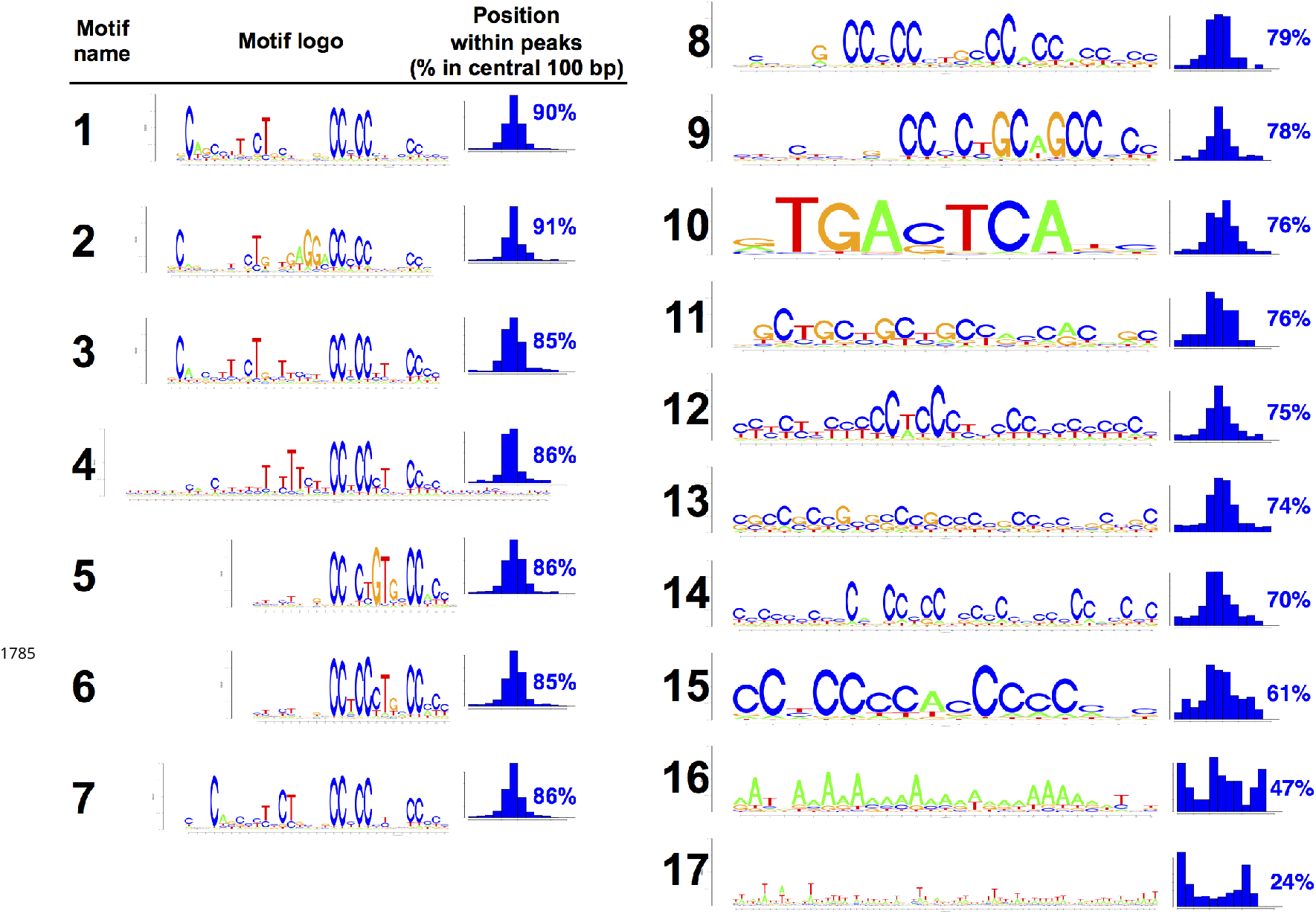
All motifs found in human PRDM9 peaks. All 17 motif logos returned by our motif-finding algorithm are listed, along with histograms indicating their positions within the central 300 bp of our human PRDM9 peaks, as a measure of how centrally enriched they are (and therefore likely to represent true binding targets). Only the seven motifs for which greater than 85% of occurrences within peaks are within 100 bp of the peak center were retained for downstream analyses. The remaining, less centrally enriched, motifs are either degenerate (as seen in mice containing the human allele: ***Davies et al., 2016***) or may arise as a consequence of PRDM9 binding to promoter regions (this would explain Motif 10, which is a near identical match to the binding motif for the transcription factor AP1).

**Figure 1-Figure supplement 4.**
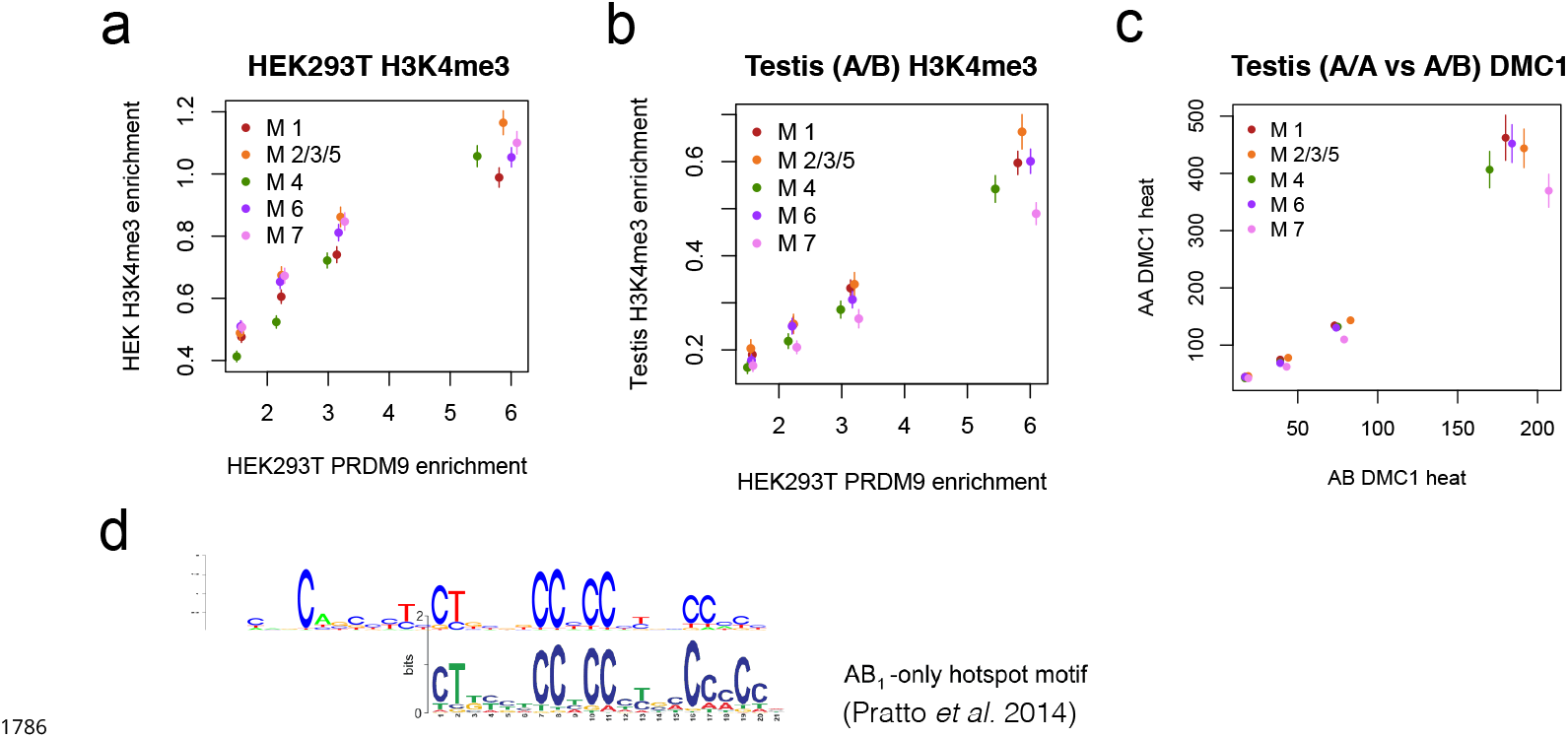
Motif 7 represents a binding mode favored by the B allele. **a**: Peak enrichment quartiles (filtered to remove promoters) were separated by motif type (Motifs 2, 3, and 5 were combined due to low abundance), and mean force-called H3K4me3 enrichment was plotted against median PRDM9 enrichment in each quartile. Error bars indicate two standard errors of the mean. This shows that the lower recombination rates for Motif 7 do not result from lower histone methylation activity of PRDM9 at those sites. **b**: Peak enrichment quartiles as in **a**, but with force-called testis H3K4me3 enrichment values from ***Pratto et al. (2014)*** in an individual with an A/B genotype. Motif 7 shows lower testis H3K4me3 enrichment for each level of PRDM9 binding, consistent with it being bound less efficiently by the A allele, **c**: At DMC1 hotspots found in both A/A and A/B individuals (from ***Pratto et al., 2014***), a comparison of mean reported heats in quartiles for each motif type. Motif 7 peaks are relatively hotter in the A/B samples than in the A/A samples. **d**: *A* comparison of Motif 7 to a reported motif obtained from A/B-only DMC1 hotspots (***Pratto et al., 2014***) shows a very close match.

**Figure 2-Figure supplement 1.**
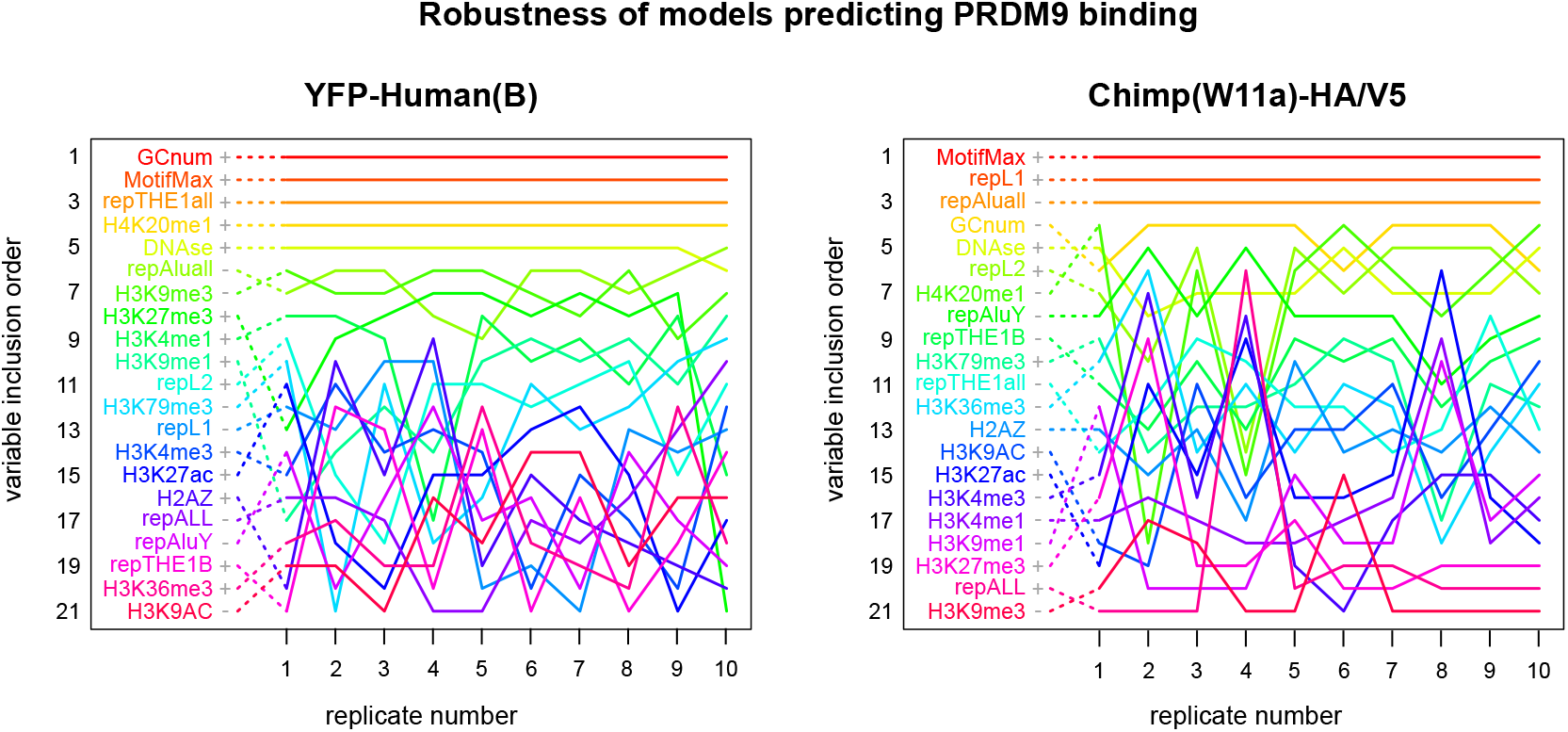
Robustness of features that predict PRDM9 binding across 10 resampled model replicates. These plots trace the forward regression inclusion order of each explanatory variable across 10 models trained on independently resampled data, as a measure of the stability of each submodel. Plus or minus symbols indicate the sign of each variable’s coefficient in the full model including all 21 variables. All features are significant in the full models (p<0.01), with the exception of H3K4me3 and H2AZ in the human model. Variables are listed in order of their mean rank across all 10 replicates, which represents their inclusion order in the final submodels evaluated on held-out test data. Dotted lines connect each variable name to its rank in the first replicate for ease of visualization. The top several features remain robustly stable across all models, while the remainder shift ranks moderately or dramatically. See Methods and Materials for a description of each explanatory variable.

**Figure 2-Figure supplement 2.**
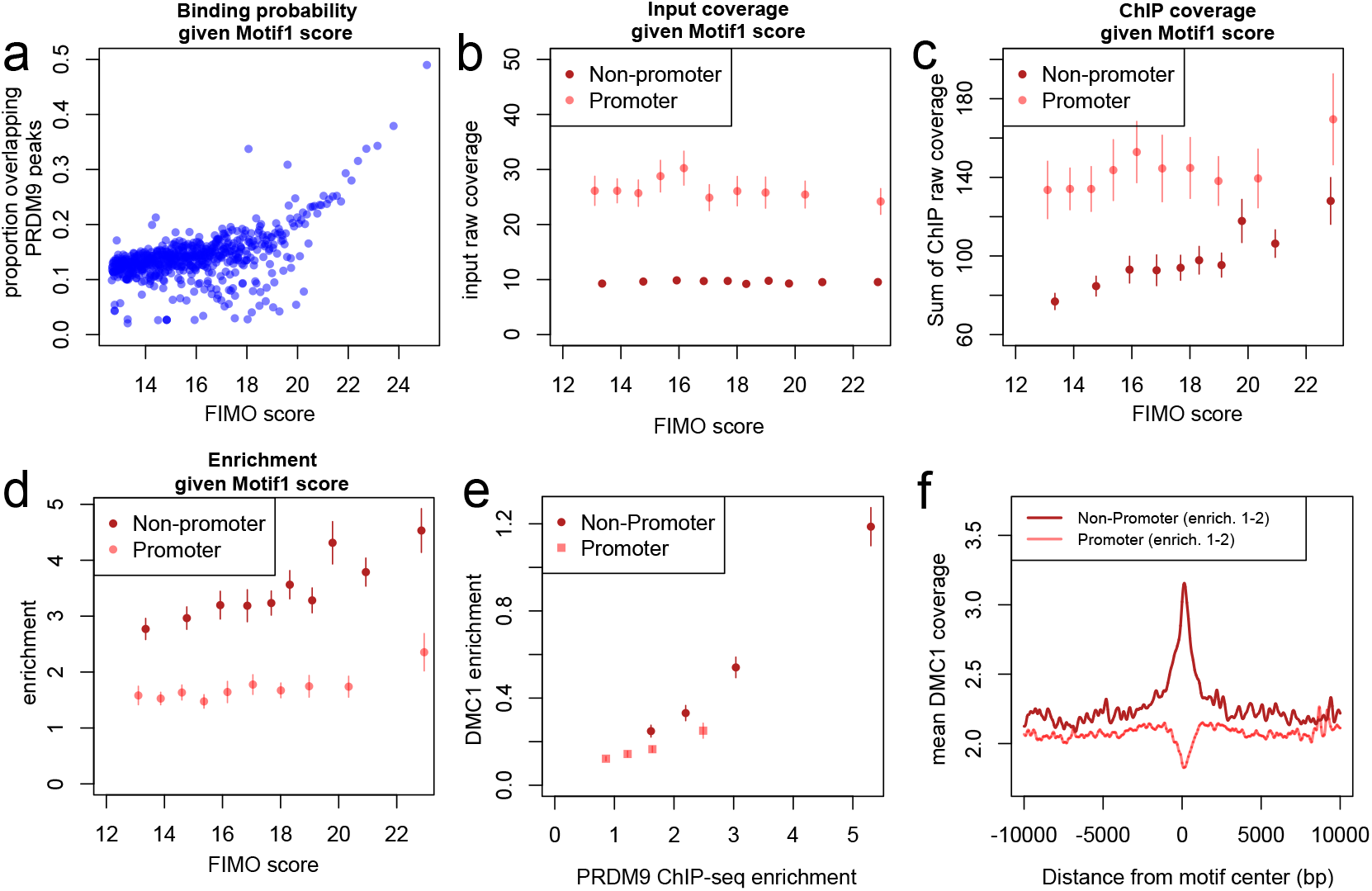
Human PRDM9 can bind promoters, though weakly, and DSBs do not occur. **a**: FIMO was used to identify the top 1 million matches for Motif 1 in hg19 (***Bailey et al., 2015***). For 0.1 percentile bins of increasing FIMO score, the proportion of motif matches occurring within 150 bp of a PRDM9 peak center is plotted (p<10^-6^, minsep 250). Even the strongest 0.1% of motif matches are only bound 50% of the time. **b**: PRDM9 peaks overlapping Motif 1 (and having more than 5 input reads overlapping the peak center) were divided into those overlapping promoters (stringently, those within 1 kb of a TSS, overlapping an H3K4me3 peak in untransfected cells, and overlapping a DNase HS site; red), and non-promoters (failing those criteria and further not overlapping an H3K4me3 peak reported by any ENCODE data; see Methods and Materials; pink). Mean raw input coverage values are plotted in decile bins of FIMO score, with error bars representing ± 2 s.e.m. **c,d**: Same as **b**, but with mean sum of raw ChIP fragment coverage values in each bin (**c**) or mean computed enrichment values in each bin (**d**). Overall, promoters show greater input sequencing coverage and thus we have greater power to detect weak binding in these regions. When corrected for this sequencing bias, we see that promoter binding sites tend to have weaker binding enrichment for a given FIMO score. **e**: Mean force-called DMC1 enrichment values (***Pratto et al., 2014***) are reported for promoter (pink squares) and non-promoter (red circles) human PRDM9 peaks split into quartiles of PRDM9 enrichment (filtered to not overlap repeats or occur within 15 Mb of a telomere; error bars represent two standard errors of the mean). Both median PRDM9 enrichment values and DMC1 enrichment values are greater for non-promoter peaks, even in overlapping ranges of PRDM9 enrichment. **f**: Mean raw DMC1 coverage in 20-kb windows centered on bound motifs, for promoter (pink) and non-promoter (red) peaks further filtered only to include peaks with PRDM9 enrichment values between 1 and 2 (smoothing: ksmooth bandwidth 200).

**Figure 3-Figure supplement 1.**
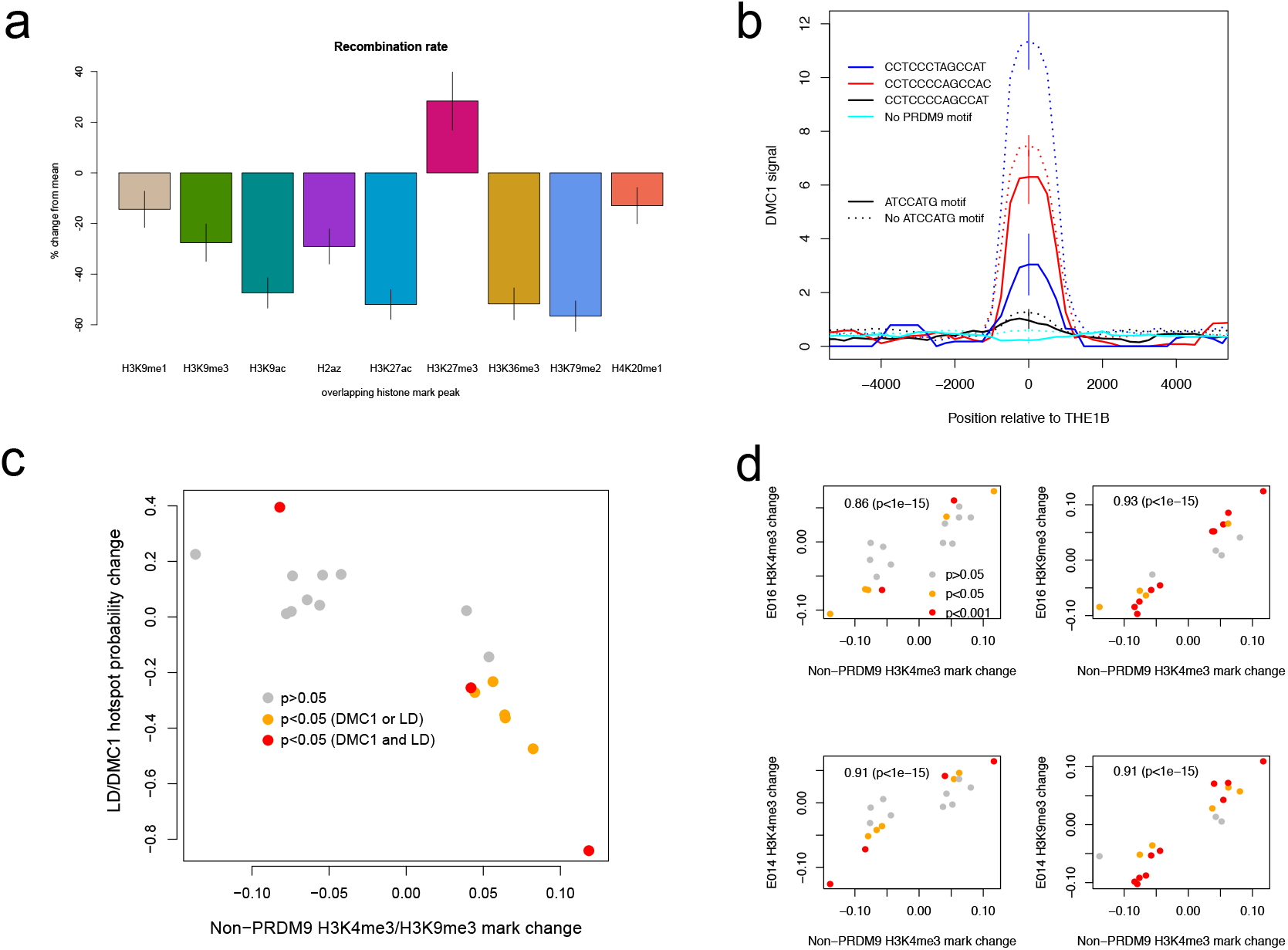
Features associated with recombination outcomes given PRDM9 binding. **a**: PRDM9 peaks were filtered (requiring each peak to: have an enrichment value in the range [1,2], have a motif match, not overlap promoters or DNase HS sites, not occur within 15 Mb of a telomere, not overlap repeats, not match Motif 7), then annotated with whether they overlap each of 9 reported histone variant peak sets reported for K562 cells (***ENCODE, 2012***). The marginal mean recombination rate is reported for peaks overlapping each histone variant type (categories are not mutually exclusive; error bars = ±2 s.e.m.; scale= % change relative to mean rate for all peaks: 2.62 cM/Mb). **b**: DMC1-based recombination rates around the centers of THE1B repeats containing different approximate matches to the PRDM9 binding motif CCTCCC[CT]AGCCA[CT] (colors) and the motif ATCCATG (lines dotted if absent). ATCCATG presence reduces recombination. Vertical lines: ± 2 s.e. **c**: For 18 motifs identified to influence H3K4me3 signal strength at THE1B repeats in testes (and H3K9me3 in other cell types, see d) but not PRDM9 (Methods and Materials) we fit a joint generalized linear model of each motif’s effect size on H3K4me3 in testes (x-axis). For the same set of motifs, we fit two joint generalized linear models to estimate each motif’s effect size on the probability a THE1B repeat overlaps respectively a DMC1 or LD-based hotspot, and average the estimated effect sizes, corresponding to an odds ratio for each motif (y-axis). Points are colored according to whether coefficients for the second linear models differ significantly from zero (legend). The strong negative correlation on the plot implies that motifs increasing H3K9me3/H3K4me3 associate with decreased recombination, and conversely. **d**: Four panels correspond to two different histone modifications H3K9me3 and H3K4me3, in two distinct somatic embryonic stem cell types (E014 and E016) studied by ROADMAP (***Kundaje et al., 2015***) and labeled accordingly on the y-axis. In each panel the x-axis is as for (c). Each y-axis gives estimated coefficients under a generalized linear model fitted in the same way as the x-axis (Methods and Materials), predicting enrichment of that particular histone modification in a particular cell type in THE1B repeats by presence/absence of each motif. Points are colored according to whether coefficients for this linear model differ significantly from zero (legend). Note strong positive correlations (each plot is labeled with rank-based correlation and p-value of rank-based correlation test) of 0.86 to 0.93, slightly higher for H3K9me3 than H3K4me3 and showing larger coefficients. The same motifs are then associated with both H3K9me3 and H3K4me3 changes across cell types including the cells lacking PRDM9 expression.

**Figure 3-Figure supplement 2.**
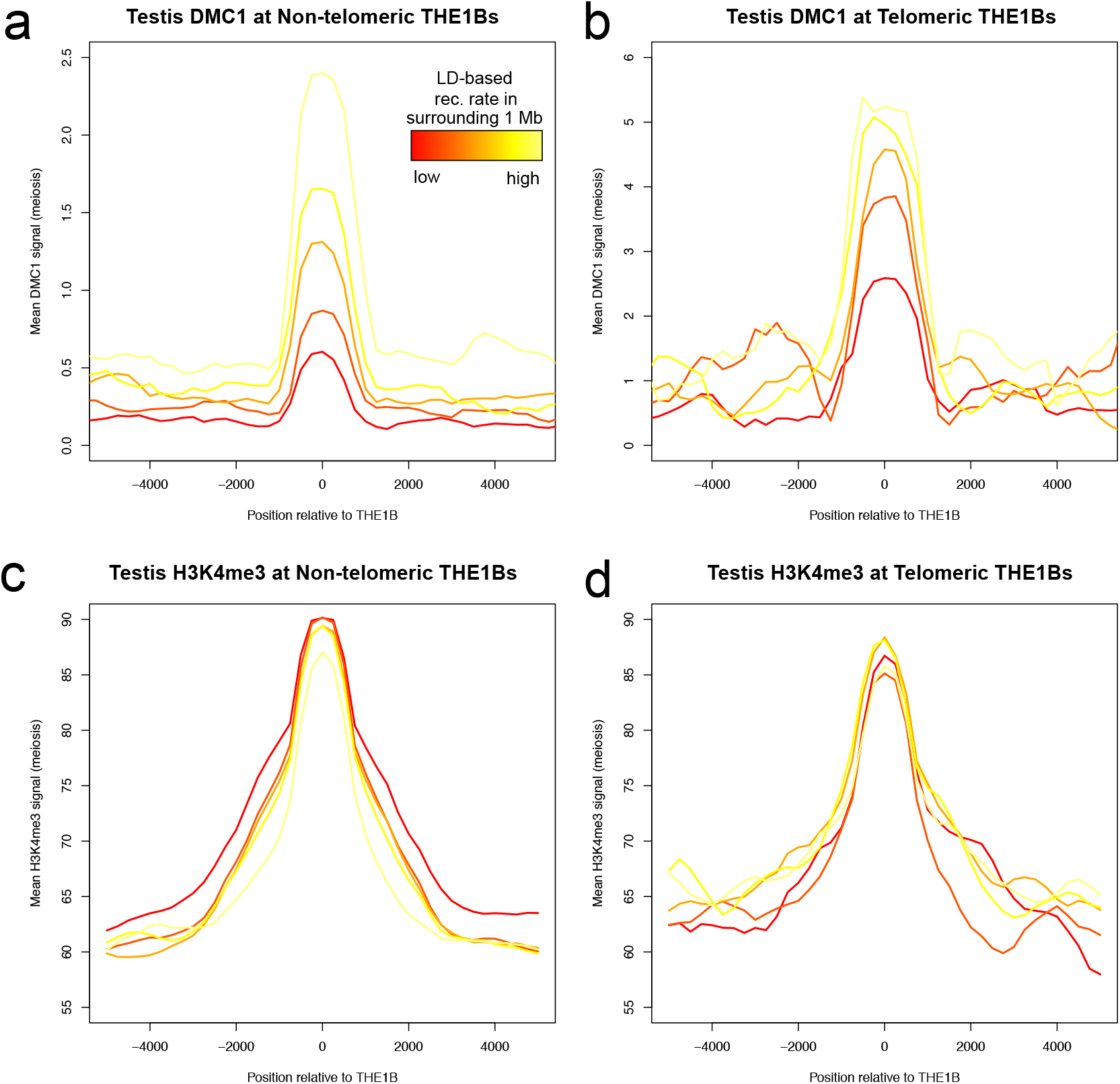
Large-scale recombination rate affects testis DMC1 but not H3K4me3. Profiles of mean DMC1 and H3K4me3 read coverage from human male testes (with a PRDM9 A/B genotype; ***Pratto et al., 2014***) around all THE1B repeats, stratified into quantiles based on the pedigree-based recombination heat in the surrounding 1 Mb of DNA (***Kong et al., 2002***), excluding the surrounding 20 kb and the repeat itself, by color (red to yellow are increasing 20% quantiles). H3K4me3 shows no impact whatsoever from surrounding recombination rate, implying PRDM9 binding is completely unaffected (c,d). However DMC1 signal increases dramatically (a,b), implying that broad-scale recombination control at these repeats occurs completely independently of PRDM9 binding or local sequence. Note the y-axes are different for telomere and non-telomere DMC1 (a,b) but not H3K4me3 (c,d). Telomeric sites were defined as those occurring within 10 Mb of a telomere, and H3K4me3 values were capped at 500 to reduce outlier effects.

**Figure 5-Figure supplement 1.**
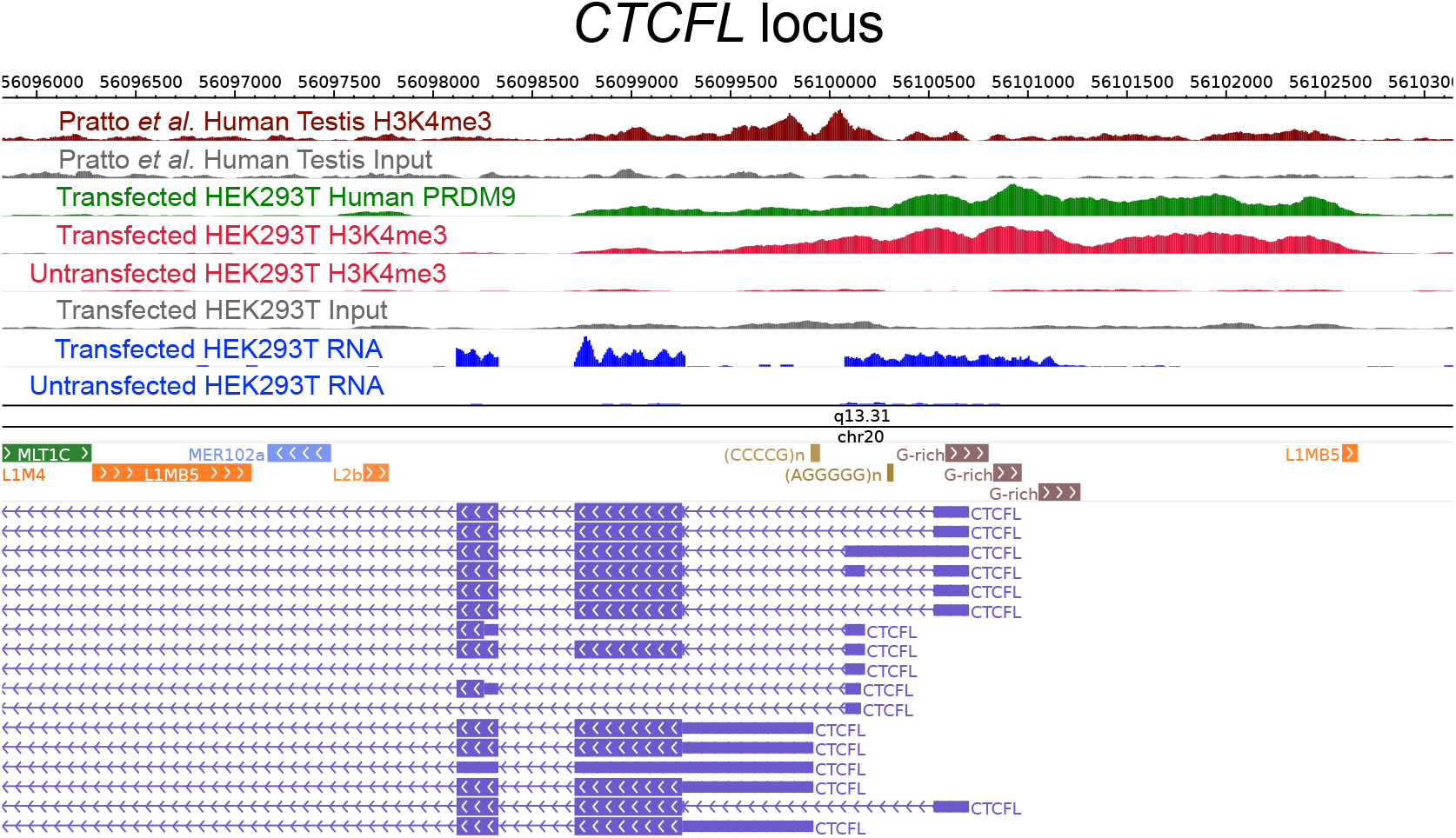
Raw coverage values surrounding the *CTCFL* promoter. A browser screenshot (***Zhou et al., 2011***) from Chr20 near the promoter region of *CTCFL* with custom tracks indicating ChIP-seq and RNA-seq raw coverage data. Human PRDM9 (green) binds a G-rich repeat near the TSS, adding an H3K4me3 mark (light red) where none is present in untransfected cells. RNA-seq coverage (blue) spikes in the coding regions in transfected cells, while it is nearly flat in untransfected cells. Testis H3K4me3 coverage (dark red, from ***Pratto et al., 2014***) peaks at a slightly different locus, corresponding to an alternative TSS. An LD-based recombination hotspot is visible in the HapMap CEU Recombination Rate track (top, black) near the promoter region.

**Figure 5-Figure supplement 2.**
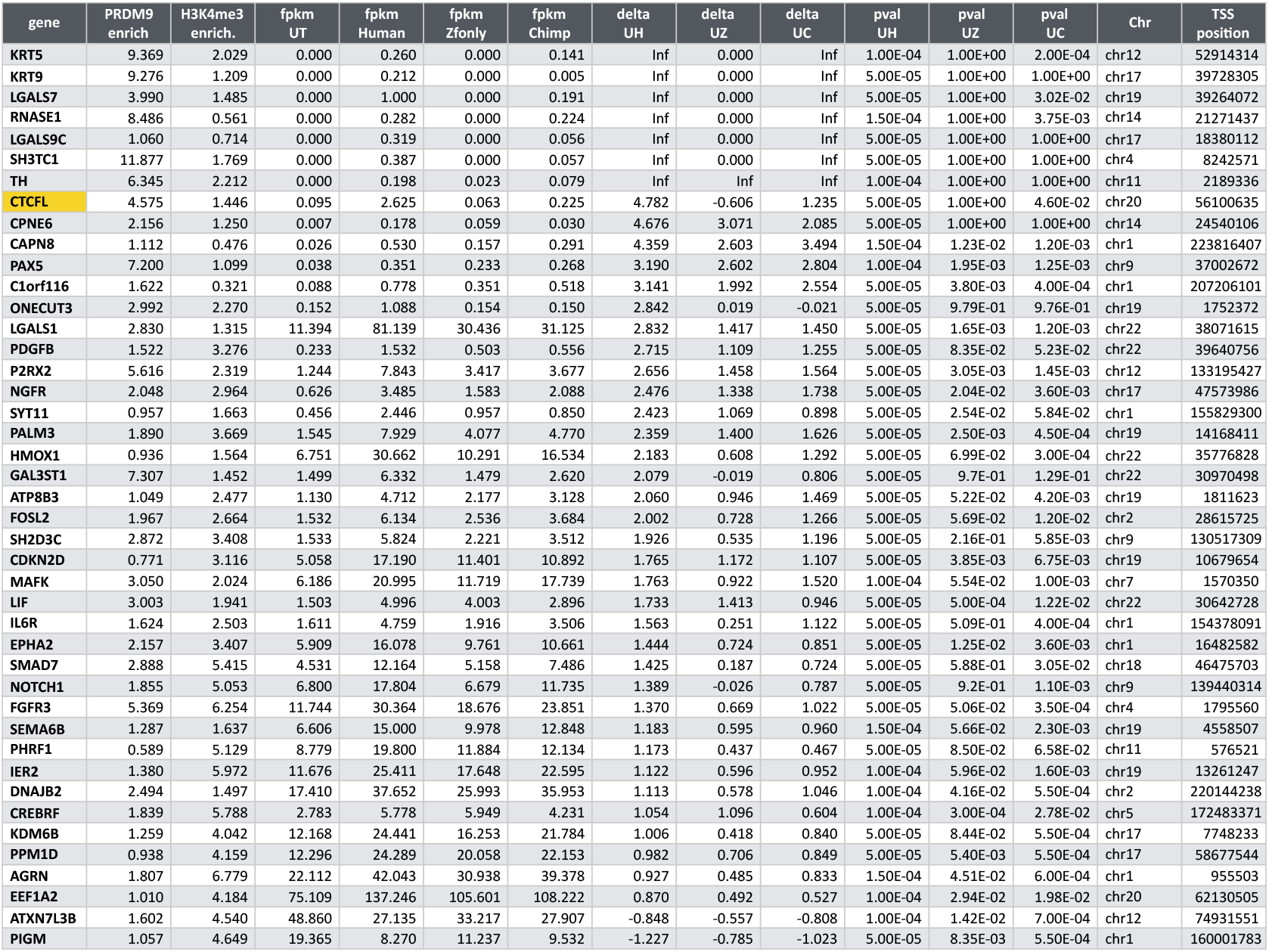
Raw coverage values surrounding the *CTCFL* promoter. Genes with significant expression differences in Human PRDM9 samples only. 46 protein coding genes with significant differential expression between human-transfected versus untransfected cells (but no significant expression change in the control transfections) are listed along with the enrichment value of the strongest PRDM9 peak within 500 bp of a TSS, the force-called H3K4me3 enrichment value around the TSS, and the RNA-seq values output by Cufflinks and CuffDiff (***Trapnell et al., 2012***). Genes are listed in reverse order of the fold expression change.

**Figure 6-Figure supplement 1.**
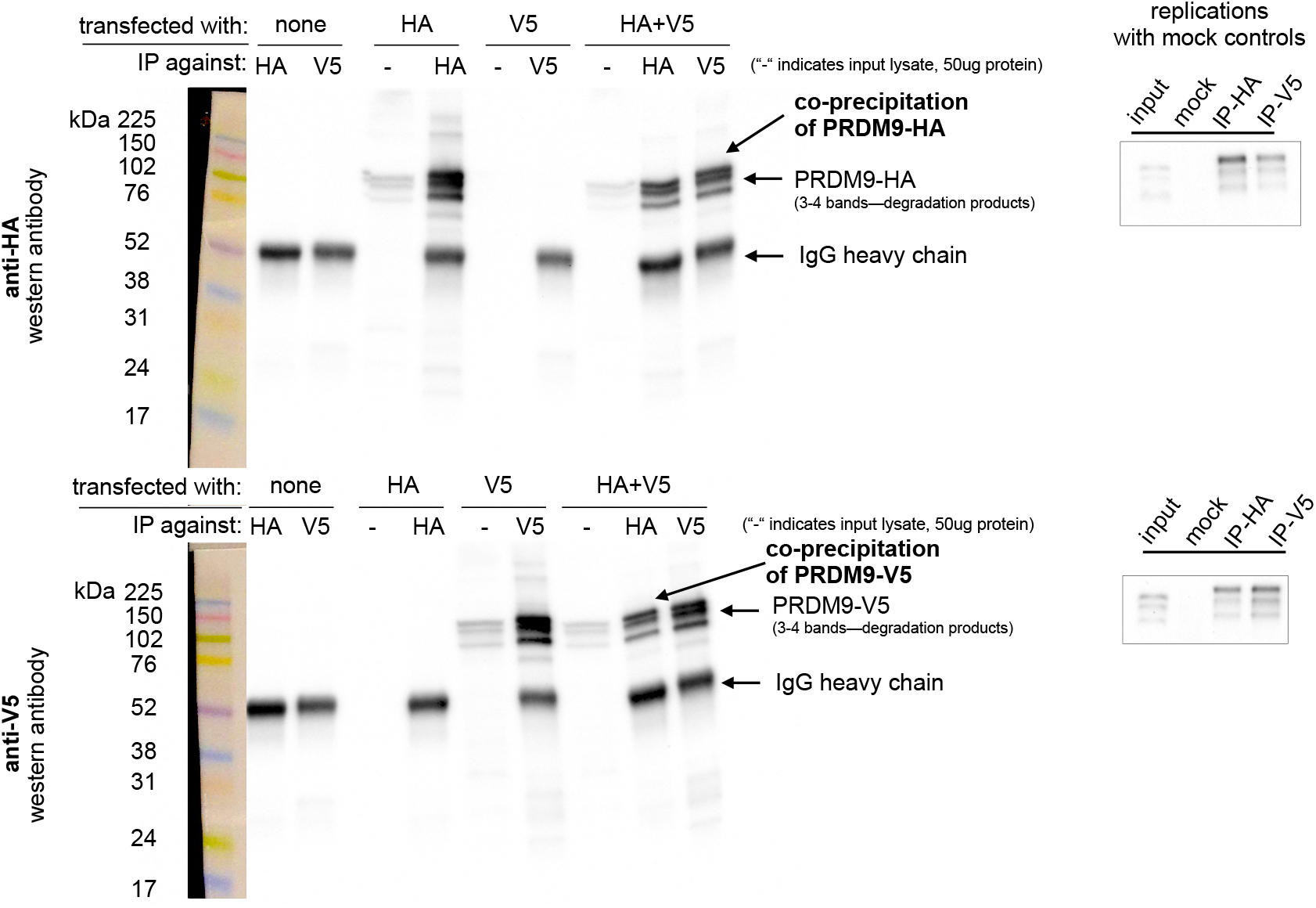
PRDM9 can form multimers when co-transfected in HEK293T cells. **Left**: Western blots illustrating controls and experimental results. Samples were split and run on two blots separately, one imaged using an anti-HA antibody (upper) and one using an anti-V5 antibody (lower). Exposure time was 4 minutes. Ladder lanes are overlaid on the left, with approximate sizes in kiloDaltons noted. Lanes are labeled according to which full-length Human construct (HA or V5) was used, as well as which antibody was used for immunoprecipitation. IgG heavy chains are visible around ~50 kDa, while the Human allele is visible as a band around ~100 kDA with two or three smaller bands beneath it, likely representing degradation products (***Grey et al., 2011; Cole et al., 2014***). “-” is a short-hand label for input lanes, for which 50 *μ*g of input chromatin was loaded in each well. The first six lanes demonstrate the specificity of the antibodies and their lack of cross-reactivity. The last two lanes show the co-IP experimental results confirming multimerization. **Right**: Two independent replicates were performed to confirm the formation of multimers with the full-length human constructs, using IgG mock control lanes to rule out nonspecific co-precipitation. Images were cropped to include only the PRDM9 bands. Input lane bands appear to have run lower than expected due to the use of a higher concentration of loading buffer in the IP lanes, an issue which was avoided in subsequent experiments.

**Figure 6-Figure supplement 2.**
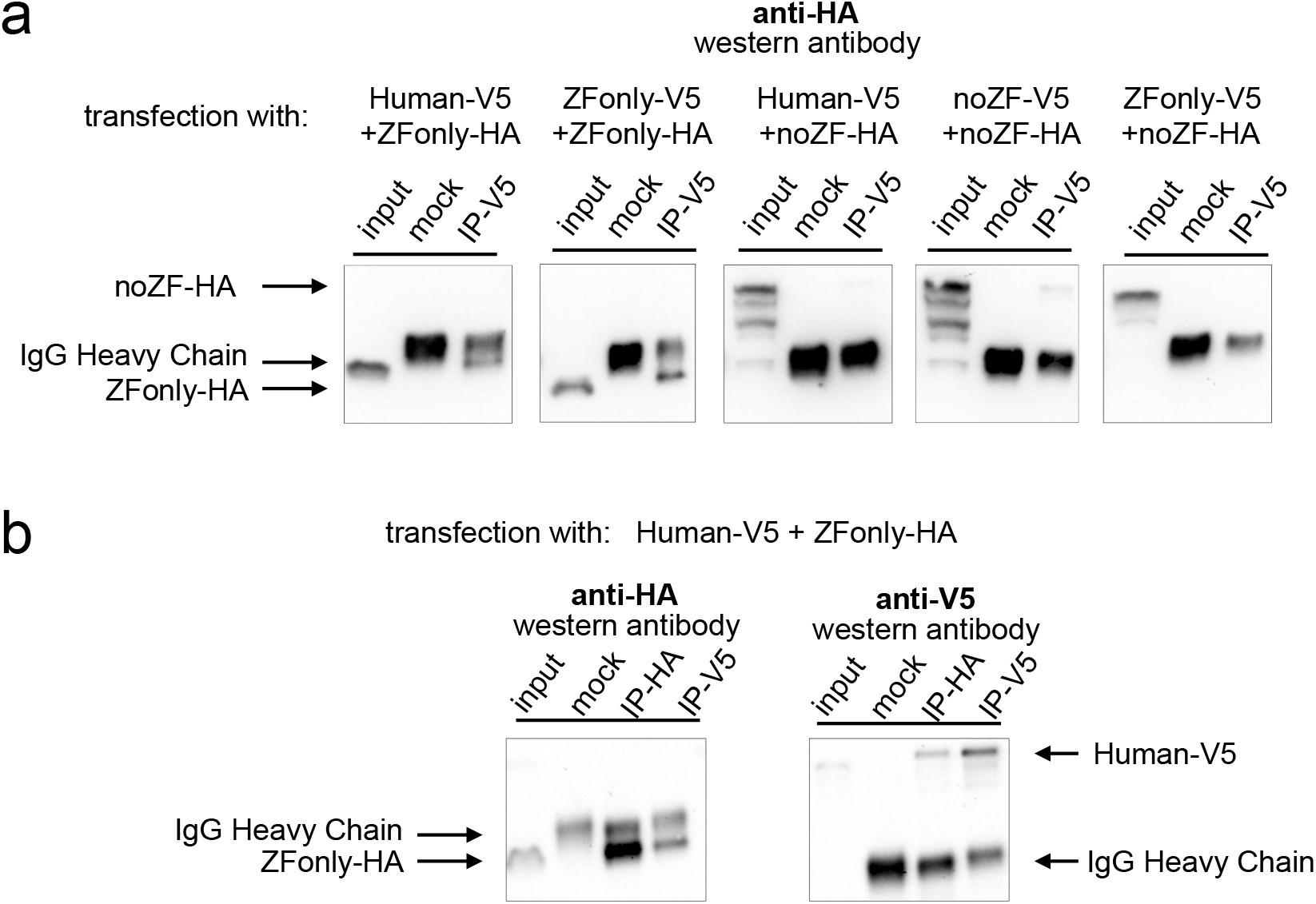
PRDM9 can form multimers when co-transfected in HEK293T cells. Multimerization is mediated primarily by ZF-ZF binding. Western blots illustrating co-IP results for various combinations of full-length human, noZF, and ZFonly constructs. **a**: The third and fourth blots show only a very faint co-IP signal despite strong input expression of the noZF construct, indicating that the non-ZF portion of PRDM9 cannot form mul-timers efficiently with itself or full-length PRDM9. The first and second blots show strong co-IP signals for the ZFonly construct, indicating that the ZF domain binds itself and binds the full-length Human construct. The fifth plot shows that the ZFonly and noZF constructs do not bind each other and confirms that multimerization is not mediated by the C-terminal tags. **b**: A replication of the experiment shown in the first blot above, but performing the IPs and western blots with both tag combinations. This confirms that the full-length Human construct can pull down the ZFonly construct, and the ZFonly construct is sufficient to pull down the full-length Human construct.

**Figure 6-Figure supplement 3.**
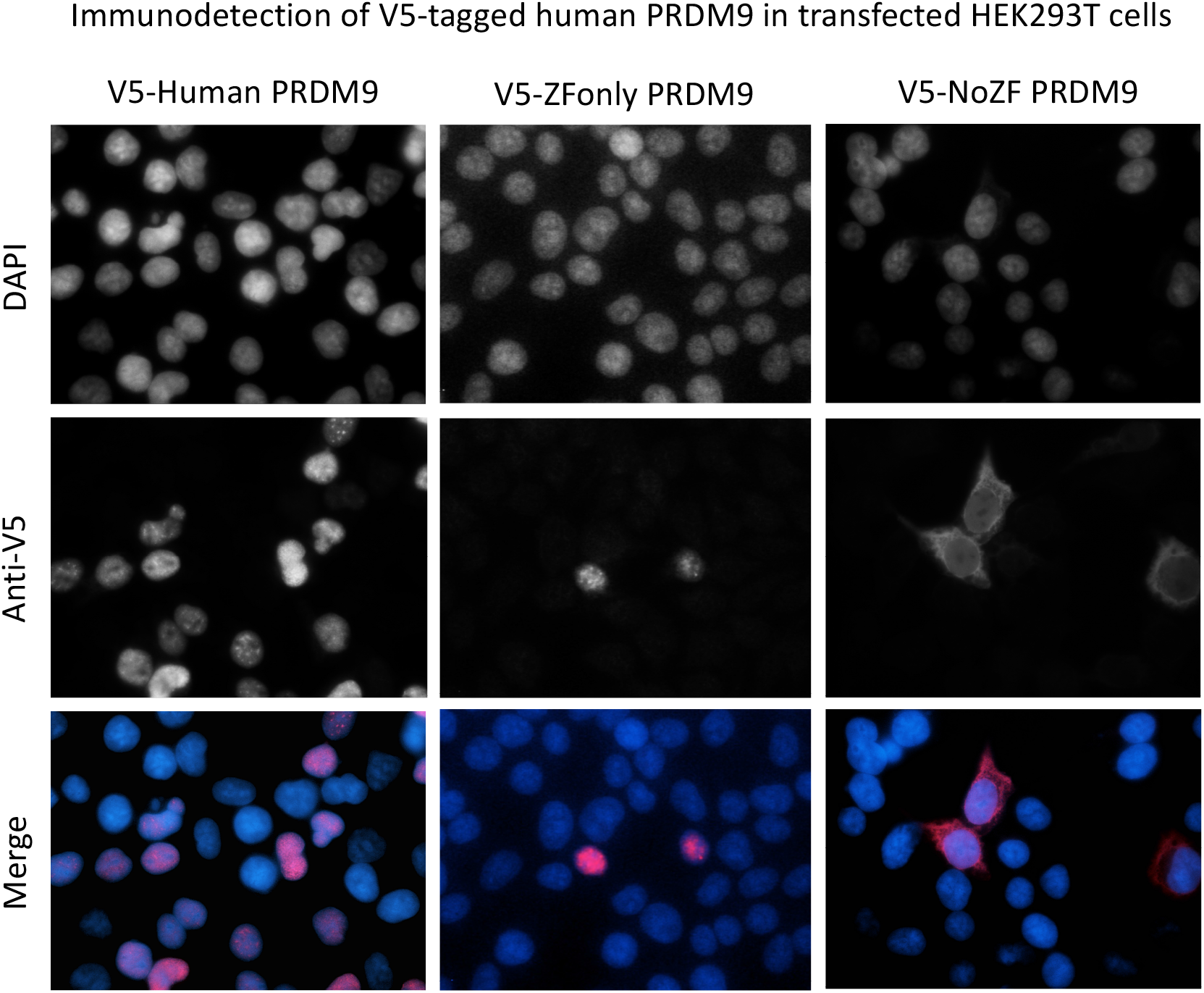
Human and ZFonly constructs localize to the nucleus.

## References

Aricescu AR, Lu W, Jones EY. A time- and cost-efficient system for high-level protein production in mammalian cells. Acta Crystallographica Section D. 2006; 62(10):1243–1250.

Auton A, Fledel-Alon A, Pfeifer S, Venn O, Segurel L, Street T, Leffler EM, Bowden R, Aneas I, Broxholme J, Humburg P, Iqbal Z, Lunter G, Maller J, Hernandez RD, Melton C, Venkat A, Nobrega MA, Bontrop R, Myers S, et al. A Fine-Scale Chimpanzee Genetic Map from Population Sequencing. Science. 2012; 336(6078):193–198.

Bailey TL, Johnson J, Grant CE, Noble WS. The MEME Suite. Nucleic Acids Research. 2015; 43(W1):W39–W49.

Baker CL, Kajita S, Walker M, Saxl RL, Raghupathy N, Choi K, Petkov PM, Paigen K. PRDM9 Drives Evolutionary Erosion of Hotspots in Mus musculus through Haplotype-Specific Initiation of Meiotic Recombination. PLoS Genetics. 2015; 11(1):e1004916–e1004916.

Baker CL, Petkova P, Walker M, Flachs P, Mihola O, Trachtulec Z, Petkov PM, Paigen K. Multimer Formation Explains Allelic Suppression of PRDM9 Recombination Hotspots. PLoS Genetics. 2015; 11(9):e1005512.

Baker CL, Walker M, Kajita S, Petkov PM, Paigen K. PRDM9 binding organizes hotspot nucleosomes and limits Holliday junction migration. Genome Research. 2014; 24(5):724–732.

Baudat F, Buard J, Grey C, Fledel-Alon A, Ober C, Przeworski M, Coop G, de Massy B. PRDM9 is a major determinant of meiotic recombination hotspots in humans and mice. Science. 2010; 327(5967):836–840.

Birtle Z, Ponting CP. Meisetz and the birth of the KRAB motif. Bioinformatics. 2006; 22(23):2841–2845.

Boulton A, Myers RS, Redfield RJ. The hotspot conversion paradox and the evolution of meiotic recombination. Proceedings of the National Academy of Sciences of the United States of America. 1997; 94(15):8058–8063.

Brick K, Smagulova F, Khil P, Camerini-Otero RD, Petukhova GV. Genetic recombination is directed away from functional genomic elements in mice. Nature. 2012; 485(7400):642–645.

Buard J, Barthès P, Grey C, de Massy B. Distinct histone modifications define initiation and repair of meiotic recombination in the mouse. The EMBO journal. 2009; 28(17):2616–2624.

Buenrostro JD, Giresi PG, Zaba LC, Chang HY, Greenleaf WJ. Transposition of native chromatin for fast and sensitive epigenomic profiling of open chromatin, DNA-binding proteins and nucleosome position. Nature Methods. 2013; 10(12):1213.

Campeau E, Ruhl VE, Rodier F, Smith CL, Rahmberg BL, Fuss JO, Campisi J, Yaswen P, Cooper PK, Kaufman PD. A Versatile Viral System for Expression and Depletion of Proteins in Mammalian Cells. PLoS ONE. 2009; 4(8):e6529.

Cano-Rodriguez D, Gjaltema RAF, Jilderda LJ, Jellema P, Dokter-Fokkens J, Ruiters MHJ, Rots MG. Writing of H3K4Me3 overcomes epigenetic silencing in a sustained but context-dependent manner. Nature communications. 2016 Aug; 7:12284.

Cole F, Baudat F, Grey C, Keeney S, de Massy B, Jasin M. Mouse tetrad analysis provides insights into recombination mechanisms and hotspot evolutionary dynamics. Nature Genetics. 2014; p. 1–11.

Collin RWJ, Nikopoulos K, Dona M, Gilissen C, Hoischen A, Boonstra FN, Poulter JA, Kondo H, Berger W, Toomes C, Tahira T, Mohn LR, Blokland EA, Hetterschijt L, Ali M, Groothuismink JM, Duijkers L, Inglehearn CF, Sollfrank L, Strom TM, et al. ZNF408 is mutated in familial exudative vitreoretinopathy and is crucial for the development of zebrafish retinal vasculature. Proceedings of the National Academy of Sciences of the United States of America. 2013; 110(24):9856–9861.

Consortium GP. An integrated map of genetic variation from 1,092 human genomes. Nature. 2012; 491(7422):56–65.

Coop G, Myers S. Live hot, die young: transmission distortion in recombination hotspots. PLoS Genetics. 2007; 3(3):e35.

Davies B, Hatton E, Altemose N, Hussin JG, Pratto F, Zhang G, Hinch AG, Moralli D, Biggs D, Diaz R, Preece C, Li R, Bitoun E, Brick K, Green CM, Camerini-Otero RD, Myers SR, Donnelly P. Re-engineering the zinc-fingers of PRDM9 reverses hybrid sterility in mice. Nature. 2016; p. doi:10.1038/nature16931.

ENCODE. An integrated encyclopedia of DNA elements in the human genome. Nature. 2012; 489(7414):57–74.

Eram MS, Bustos SP, Lima-Fernandes E, Siarheyeva A, Senisterra G, Hajian T, Chau I, Duan S, Wu H, Dombrovski L, Schapira M, Arrowsmith CH, Vedadi M. Trimethylation of Histone H3 Lysine 36 by Human Methyltransferase PRDM9 Protein. Journal of Biological Chemistry. 2014; 289(17):12177–12188.

Galanty Y, Belotserkovskaya R, Coates J, Polo S, Miller KM, Jackson SP. Mammalian SUMO E3-ligases PIAS1 and PIAS4 promote responses to DNA double-strand breaks. Nature. 2009; 462(7275):935–939.

Grey C, Barthès P, Chauveau-Le Friec G, Langa F, Baudat F, de Massy B. Mouse PRDM9 DNA-Binding Specificity Determines Sites of Histone H3 Lysine 4 Trimethylation for Initiation of Meiotic Recombination. PLoS Biology. 2011; 9(10):e1001176.

Grey C, Clément JAJ, Buard J, Leblanc B, Gut I, Gut M, Duret L, de Massy B. In vivo binding of PRDM9 reveals interactions with noncanonical genomic sites. Genome research. 2017; p. 1–12.

HapMap. A second generation human haplotype map of over 3.1 million SNPs. Nature. 2007; 449(7164):851–861.

Hayashi K, Yoshida K, Matsui Y. A histone H3 methyltransferase controls epigenetic events required for meiotic prophase. Nature. 2005; 438(7066):374–378.

Hinch AG, Altemose N, Noor N, Donnelly P, Myers SR. Recombination in the Human Pseudoautosomal Region PAR1. PLoS Genetics. 2014; 10(7):e1004503–e1004503.

Hinch AG, Tandon A, Patterson N, Song Y, Rohland N, Palmer CD, Chen GK, Wang K, Buxbaum SG, Akylbekova EL, Aldrich MC, Ambrosone CB, Amos C, Bandera EV, Berndt SI, Bernstein L, Blot WJ, Bock CH, Boerwinkle E, Cai Q, et al. The landscape of recombination in African Americans. Nature. 2011; 476(7359):170–175.

Hines WC, Bazarov AV, Mukhopadhyay R, Yaswen P. BORIS (CTCFL) is not expressed in most human breast cell lines and high grade breast carcinomas. PLoS ONE. 2010; 5(3):e9738.

Imbeault M, Helleboid PY, Trono D. KRAB zinc-finger proteins contribute to the evolution of gene regulatory networks. Nature. 2017; 543(7646):550–554.

Jacobs FMJ, Greenberg D, Nguyen N, Haeussler M, Ewing AD, Katzman S, Paten B, Salama SR, Haussler D. An evolutionary arms race between KRAB zinc-finger genes ZNF91/93 and SVA/L1 retrotransposons. Nature. 2014; 516(7530):242–245.

Jain D, Baldi S, Zabel A, Straub T, Becker PB. Active promoters give rise to false positive ‘Phantom Peaks’ in ChIP-seq experiments. Nucleic acids research. 2015; 43(14):6959–6968.

Johnson DS, Mortazavi A, Myers RM, Wold B. Genome-wide mapping of in vivo protein-DNA interactions. Science. 2007; 316(5830):1497–1502.

Jørgensen S, Schotta G, Sørensen CS. Histone H4 Lysine 20 methylation: key player in epigenetic regulation of genomic integrity. Nucleic acids research. 2013; 41(5):2797–2806.

Kong A, Gudbjartsson DF, Sainz J, Jonsdottir GM. A high-resolution recombination map of the human genome. Nature. 2002; 31(3):241–247.

Kong A, Thorleifsson G, Frigge ML, Masson G, Gudbjartsson DF, Villemoes R, Magnusdottir E, Olafsdottir SB, Thorsteinsdottir U, Stefansson K. Common and low-frequency variants associated with genome-wide recombination rate. Nature Genetics. 2014; 46(1):11–16.

Kundaje A, Meuleman W, Ernst J, Bilenky M, Yen A, Heravi-Moussavi A, Kheradpour P, Zhang Z, Wang J, Ziller MJ, Amin V, Whitaker JW, Schultz MD, Ward LD, Sarkar A, Quon G, Sandstrom RS, Eaton ML, Wu YC, Pfenning A, et al. Integrative analysis of 111 reference human epigenomes. Nature. 2015; 518(7539):317–330.

Lahn BT, Page DC. A human sex-chromosomal gene family expressed in male germ cells and encoding variably charged proteins. Human Molecular Genetics. 2000; 9(2):311–319.

Landt SG, Marinov GK, Kundaje A, Kheradpour P, Pauli F, Batzoglou S, Bernstein BE, Bickel P, Brown JB, Cayting P, Chen Y, DeSalvo G, Epstein C, Fisher-Aylor KI, Euskirchen G, Gerstein M, Gertz J, Hartemink AJ, Hoffman MM, Iyer VR, et al. ChIP-seq guidelines and practices of the ENCODE and modENCODE consortia. Genome research. 2012; 22(9):1813–1831.

Lange J, Yamada S, Tischfield SE, Pan J, Kim S, Zhu X, Socci ND, Jasin M, Keeney S. The Landscape of Mouse Meiotic Double-Strand Break Formation, Processing, and Repair. Cell. 2016; 167(3):695–708.e16.

Lee SJ, Lee JR, Hah H, Kim YH, Ahn JH. PIAS1 interacts with the KRAB zinc finger protein, ZNF133, via zinc finger motifs and regulates its transcriptional activity. Experimental and molecular medicine. 2007; 39(4):450–457.

Li H, Durbin R. Fast and accurate short read alignment with Burrows-Wheeler transform. Bioinformatics. 2009; 25(14):1754–1760.

Li H, Handsaker B, Wysoker A, Fennell T, Ruan J. The sequence alignment/map format and SAMtools. Bioinformatics. 2009; 25(16):2078–2079.

Lunter G, Goodson M. Stampy: a statistical algorithm for sensitive and fast mapping of Illumina sequence reads. Genome Research. 2011;.

McCarty AS, Kleiger G, Eisenberg D, Smale ST. Selective dimerization of a C2H2 zinc finger subfamily. Molecular Cell. 2003; 11(2):459–470.

Mihola O, Trachtulec Z, Vlcek C, Schimenti JC, Forejt J. A mouse speciation gene encodes a meiotic histone H3 methyltransferase. Science. 2009; 323(5912):373–375.

Myers S, Bottolo L, Freeman C, McVean G, Donnelly P. A fine-scale map of recombination rates and hotspots across the human genome. Science. 2005; 310(5746):321–324.

Myers S, Bowden R, Tumian A, Bontrop RE, Freeman C, MacFie TS, McVean G, Donnelly P. Drive against hotspot motifs in primates implicates the PRDM9 gene in meiotic recombination. Science. 2010; 327(5967):876–879.

Myers S, Freeman C, Auton A, Donnelly P, McVean G. A common sequence motif associated with recombination hot spots and genome instability in humans. Nature Genetics. 2008; 40(9):1124–1129.

Nakahashi H, Kwon KRK, Resch W, Vian L, Dose M, Stavreva D, Hakim O, Pruett N, Nelson S, Yamane A, Qian J, Dubois W, Welsh S, Phair RD, Pugh BF, Lobanenkov V, Hager GL, Casellas R. A genome-wide map of CTCF multivalency redefines the CTCF code. Cell Reports. 2013; 3(5):1678–1689.

Neale MJ, Keeney S. Clarifying the mechanics of DNA strand exchange in meiotic recombination. Nature. 2006; 442(7099):153–158.

Parvanov ED, Petkov PM, Paigen K. Prdm9 controls activation of mammalian recombination hotspots. Science. 2010; 327(5967):835.

Parvanov ED, Tian H, Billings T, Saxl RL, Spruce C, Aithal R, Krejci L, Paigen K, Petkov PM. PRDM9 interactions with other proteins provide a link between recombination hotspots and the chromosomal axis in meiosis. Molecular biology of the cell. 2016; 28(3):488–499.

Persikov AV, Singh M. An expanded binding model for Cys2His2 zinc finger protein-DNA interfaces. Physical Biology. 2011; 8(3):035010.

Persikov AV, Osada R, Singh M. Predicting DNA recognition by Cys2His2 zinc finger proteins. Bioinformatics (Oxford, England). 2009; 25(1):22–29.

Persikov AV, Singh M. De novo prediction of DNA-binding specificities for Cys2His2 zinc finger proteins. Nucleic acids research. 2014; 42(1):97–108.

Powers NR, Parvanov ED, Baker CL, Walker M, Petkov PM, Paigen K. The Meiotic Recombination Activator PRDM9 Trimethylates Both H3K36 and H3K4 at Recombination Hotspots In Vivo. PLoS genetics. 2016 Jun; 12(6):e1006146.

Pratto F, Brick K, Khil P, Smagulova F, Petukhova GV, Camerini-Otero RD. DNA recombination. Recombination initiation maps of individual human genomes. Science. 2014; 346(6211):1256442.

Quinlan ARA, Hall IMI. BEDTools: a flexible suite of utilities for comparing genomic features. Bioinformatics. 2010; 26(6):841–842.

Rowe HM, Kapopoulou A, Corsinotti A, Fasching L, Macfarlan TS, Tarabay Y, Viville S, Jakobsson J, Pfaff SL, Trono D. TRIM28 repression of retrotransposon-based enhancers is necessary to preserve transcriptional dynamics in embryonic stem cells. Genome research. 2013; 23(3):452–461.

Santos-Rosa H, Schneider R, Bannister AJ, Sherriff J, Bernstein BE, Emre NCT, Schreiber SL, Mellor J, Kouzarides T. Active genes are tri-methylated at K4 of histone H3. Nature. 2002; 419(6905):407–411.

Schwartz JJ, Roach DJ, Thomas JH, Shendure J. Primate evolution of the recombination regulator PRDM9. Nature Communications. 2014; 5:4370.

Sleutels F, Soochit W, Bartkuhn M, Heath H, Dienstbach S, Bergmaier P, Franke V, Rosa-Garrido M, van de Nobelen S, Caesar L, van der Reijden M, Bryne JC, van IJcken W, Grootegoed JA, Delgado MD, Lenhard B, Renkawitz R, Grosveld F, Galjart N. The male germ cell gene regulator CTCFL is functionally different from CTCF and binds CTCF-like consensus sites in a nucleosome composition-dependent manner. Epigenetics & Chromatin. 2012; 5(1):8.

Smagulova F, Gregoretti IV, Brick K, Khil P, Camerini-Otero RD, Petukhova GV. Genome-wide analysis reveals novel molecular features of mouse recombination hotspots. Nature. 2011; 472(7343):375–378.

Smagulova F, Brick K, Pu Y, Camerini-Otero RD, Petukhova GV. The evolutionary turnover of recombination hot spots contributes to speciation in mice. Genes & development. 2016; 30(3):266–280.

Striedner Y, Schwarz T, Welte T, Futschik A, Rant U, Tiemann-Boege I. The long zinc finger domain of PRDM9 forms a highly stable and long-lived complex with its DNA recognition sequence. Chromosome research: an international journal on the molecular, supramolecular and evolutionary aspects of chromosome biology. 2017; 112(695–708):2109.

Sun F, Fujiwara Y, Reinholdt LG, Hu J, Saxl RL, Baker CL, Petkov PM, Paigen K, Handel MA. Nuclear localization of PRDM9 and its role in meiotic chromatin modifications and homologous synapsis. Chromosoma. 2015; p. 1–19.

Trapnell C, Roberts A, Goff L, Pertea G, Kim D, Kelley DR, Pimentel H, Salzberg SL, Rinn JL, Pachter L. Differential gene and transcript expression analysis of RNA-seq experiments with TopHat and Cufflinks. Nature Protocols. 2012; 7(3):562–578.

Van Esch H, Hollanders K, Badisco L, Melotte C, Van Hummelen P, Vermeesch JR, Devriendt K, Fryns JP, Marynen P, Froyen G. Deletion of VCX-A due to NAHR plays a major role in the occurrence of mental retardation in patients with X-linked ichthyosis. Human Molecular Genetics. 2005; 14(13):1795–1803.

Walker M, Billings T, Baker CL, Powers N, Tian H, Saxl RL, Choi K, Hibbs MA, Carter GW, Handel MA, Paigen K, Petkov PM. Affinity-seq detects genome-wide PRDM9 binding sites and reveals the impact of prior chromatin modifications on mammalian recombination hotspot usage. Epigenetics & Chromatin. 2015; 8:31.

Wang J, Zhuang J, Iyer S, Lin X, Whitfield TW, Greven MC, Pierce BG, Dong X, Kundaje A, Cheng Y, Rando OJ, Birney E, Myers RM, Noble WS, Snyder M, Weng Z. Sequence features and chromatin structure around the genomic regions bound by 119 human transcription factors. Genome research. 2012; 22(9):1798–1812.

Wang W, Cai J, Lin Y, Liu Z, Ren Q, Hu L, Huang Z, Guo M, Li W. Zinc fingers function cooperatively with KRAB domain for nuclear localization of KRAB-containing zinc finger proteins. PLoS ONE. 2014; 9(3):e92155.

Wolf G, Yang P, Füchtbauer AC, Füchtbauer EM, Silva AM, Park C, Wu W, Nielsen AL, Pedersen FS, Macfarlan TS. The KRAB zinc finger protein ZFP809 is required to initiate epigenetic silencing of endogenous retroviruses. Genes & development. 2015 Mar; 29(5):538–554.

Wu H, Mathioudakis N, Diagouraga B, Dong A, Dombrovski L, Baudat F, Cusack S, de Massy B, Kadlec J. Molecular Basis for the Regulation of the H3K4 Methyltransferase Activity of PRDM9. Cell Reports. 2013; 5(1):13–20.

Young JM, Whiddon JL, Yao Z, Kasinathan B, Snider L, Geng LN, Balog J, Tawil R, van der Maarel SM, Tapscott SJ. DUX4 binding to retroelements creates promoters that are active in FSHD muscle and testis. PLoS genetics. 2013; 9(11):e1003947.

Zhou X, Maricque B, Xie M, Li D, Sundaram V, Martin EA, Koebbe BC, Nielsen C, Hirst M, Farnham P, Kuhn RM, Zhu J, Smirnov I, Kent WJ, Haussler D, Madden PAF, Costello JF, Wang T. The Human Epigenome Browser at Washington University. Nature methods. 2011; 8(12):989–990.

